# Non-canonical DNA in human and other ape telomere-to-telomere genomes

**DOI:** 10.1101/2024.09.02.610891

**Authors:** Linnéa Smeds, Kaivan Kamali, Iva Kejnovská, Eduard Kejnovský, Francesca Chiaromonte, Kateryna D. Makova

## Abstract

Non-canonical (non-B) DNA structures—e.g., bent DNA, hairpins, G-quadruplexes (G4s), Z-DNA, etc.—which form at certain sequence motifs (e.g., A-phased repeats, inverted repeats, etc.), have emerged as important regulators of cellular processes and drivers of genome evolution. Yet, they have been understudied due to their repetitive nature and potentially inaccurate sequences generated with short-read technologies. Here we comprehensively characterize such motifs in the long-read telomere-to-telomere (T2T) genomes of human, bonobo, chimpanzee, gorilla, Bornean orangutan, Sumatran orangutan, and siamang. Non-B DNA motifs are enriched at the genomic regions added to T2T assemblies, and occupy 9-15%, 9-11%, and 12-38% of autosomes, and chromosomes X and Y, respectively. G4s and Z-DNA are enriched at promoters and enhancers, as well as at origins of replication. Repetitive sequences harbor more non-B DNA motifs than non-repetitive sequences, especially in the short arms of acrocentric chromosomes. Most centromeres and/or their flanking regions are enriched in at least one non-B DNA motif type, consistent with a potential role of non-B structures in determining centromeres. Our results highlight the uneven distribution of predicted non-B DNA structures across ape genomes and suggest their novel functions in previously inaccessible genomic regions.

**Graphical Abstract:** 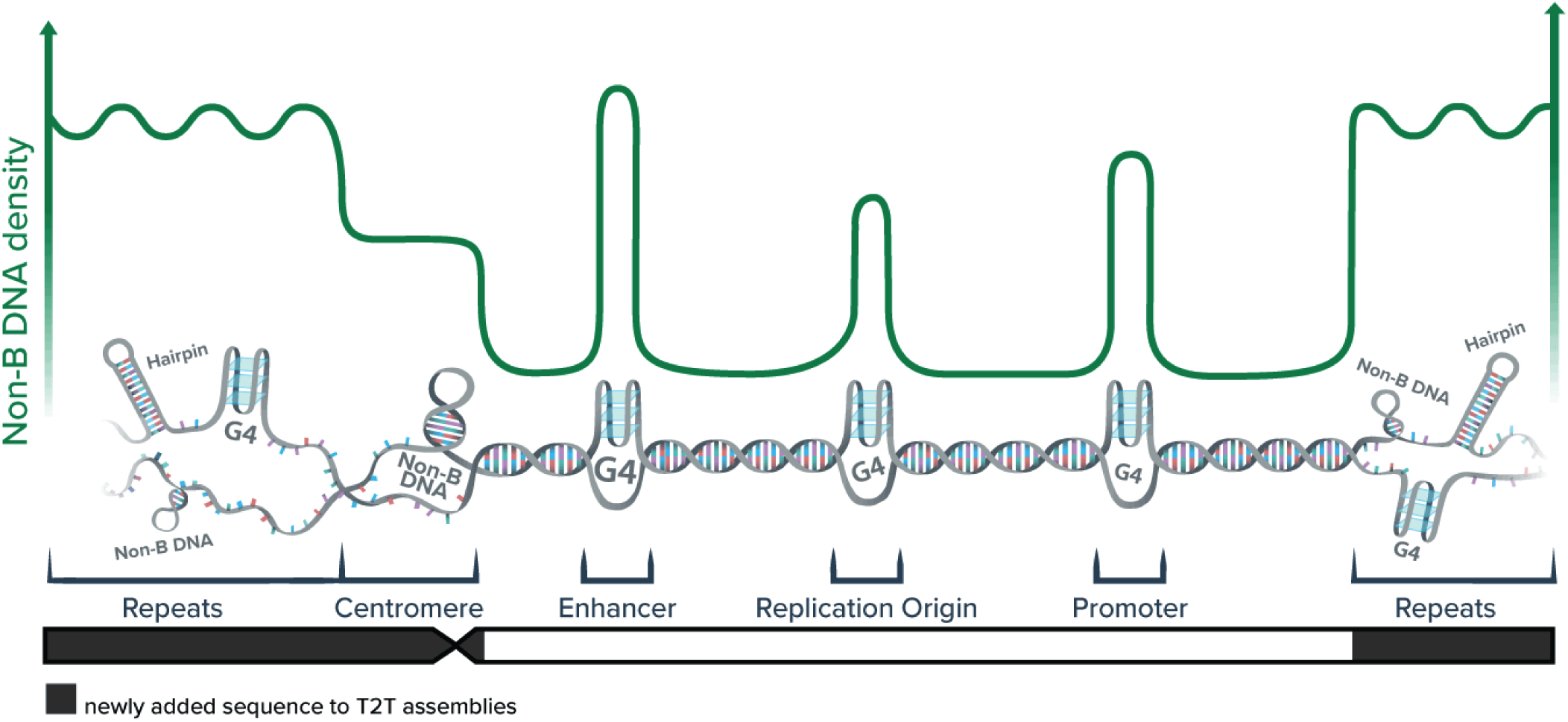

## Introduction

In addition to canonical B DNA—the right-handed double helix with 10 base pairs per turn (1)—an estimated 13% of the human genome has the ability to fold into non-canonical (non-B) DNA structures (2). Such non-B DNA conformations include cruciforms and hairpins formed by inverted repeats, triple helices (or H-DNA) formed by some mirror repeats, slipped strands formed by direct repeats, left-handed Z-DNA with 12 base pairs per turn formed by alternating purines and pyrimidines, and G-quadruplexes (G4s) formed by ≥4 ‘stems’ consisting of ≥3 guanines and ‘loops’ consisting of any 1-7 bases (Fig. 1). Non-B DNA sequence motifs range from tens to hundreds of nucleotides in length, and are present in tens of thousands of copies in the human genome (2). Non-B DNA structure formation depends on cellular conditions: DNA at a non-B motif can fold into either B or non-B form at a given time. For instance, folding into non-B forms is sensitive to oxidative stress (3, 4) and to temporal signals during cell differentiation and development (5, 6).

**Figure 1.**
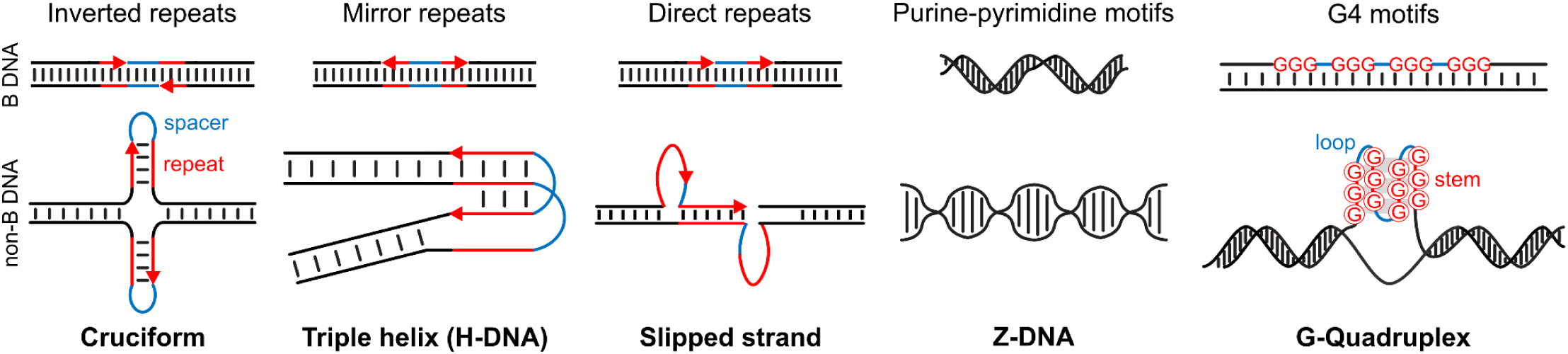
Types of non-B DNA. For structures formed from repeats: repeats are red, spacers are blue. For G4s: stems are red, loops are blue. Triple helix is formed by mirror repeats comprising predominantly purines or pyrimidines and having a short spacer (less than 9 bp) (7).

Non-B DNA is increasingly recognized as a major regulator of myriad processes in the mammalian cell. Non-B DNA structures are involved in replication initiation (8, 9). G4s affect the life cycle of L1 transposable elements (10) and protect chromosome ends at telomeres (11). Non-B DNA has been implicated in regulating transcription (12–24). G4s regulate chromatin organization (24–29) and methylation of CpG islands (30). The transcribed non-B DNA motifs can form structured RNA, which regulates alternative splicing (31), translation of mRNA (16, 32, 33), and function of non-coding RNA (34).

Non-B DNA has also been implicated in the definition and function of centromeres. For example, inverted repeats forming non-B DNA have been hypothesized to define centromeres (35), which would resolve the CENP-B paradox. Indeed, non-B DNA might play a role attributed to CENP-B, the highly conserved protein binding motif present at centromeres across a range of taxa and proposed to be involved in centromere formation—but paradoxically missing entirely on some chromosomes (36, 37). Recent studies also found enrichment at centromeres for G4s in Drosophila (38) and Z-DNA, and A-phased, direct, and mirror repeats in plants (39, 40) and argued that non-B DNA is important for centromere activity and stability.

Notwithstanding their important functions, non-B structures may impede replication and elevate mutagenesis and genome instability. For instance, they can increase pausing and decrease the accuracy of replicative DNA polymerases *in vitro* (41–43). The cell recruits error-prone specialized helicases (44) and polymerases (45–48) to handle non-B DNA structures during replication. Moreover, non-B DNA affects the efficiency of DNA repair pathways (30, 42, 49). Increased mutagenesis and genomic instability at non-B DNA are evident in cancers with mutated components of these pathways (42). In non-cancerous cells, the effects of non-B DNA on replication progression, mutation rate, and genome instability remain controversial (50). Nevertheless, non-B DNA has been recognized as an important driver of genome evolution (51).

Non-B DNA structures have been implicated in neurodegenerative diseases (e.g., amyotrophic lateral sclerosis (52) and fragile X syndrome (53)). They are also the preferential sites of genome rearrangements (54) and affect gene expression (53) in cancers. Some diseases result from mutations in genes encoding proteins processing non-B DNA (e.g., Werner syndrome) (55).

Despite its unequivocal importance for genome function, mutations, and diseases, studying non-B DNA has been challenging for several reasons. First, sequencing technologies, particularly the short-read Illumina technology, have increased error rates at non-B DNA motifs (56, 57). To overcome this limitation, the current recommendation is to use multiple long-read sequencing technologies as they differ in their biases at non-B DNA motifs (56). Second, incomplete genome assemblies have hindered the full characterization of non-B DNA motifs, particularly those in repetitive regions.

Here, we identify non-B DNA motif occurrences in the complete, telomere-to-telomere (T2T) genomes of human (58, 59) and several non-human apes—bonobo and chimpanzee (which diverged from each other ∼2.5 million years ago, Mya, and from the human lineage ∼7 Mya), gorilla (which diverged from human and bonobo/chimpanzee ∼9 Mya), Bornean and Sumatran orangutans (which diverged from each other ∼1 Mya, and from the previously mentioned species ∼17 Mya), and the lesser ape siamang (which diverged from great apes ∼20 Mya) (60, 61). This provides a comprehensive view of non-B DNA genomic distribution across most living great ape species and an outgroup. Importantly, the human and primate T2T assemblies employed in our study were produced with two long-read sequencing technologies, thus minimizing the effects of sequencing biases at non-B DNA motifs. Using this exhaustive dataset, we tackle several questions that could not be addressed prior to the availability of complete ape genomes, including the potential enrichment of non-B DNA at centromeres and satellites. We further investigate G4 formation at satellites using methylation data from two cell lines, and validate some commonly found G4 motifs experimentally with circular dichroism (CD) analysis.

## Materials & Methods

### Non-B DNA annotations

Non-B DNA motifs were annotated for bonobo (*Pan paniscus*), chimpanzee (*Pan troglodytes*), human (*Homo sapiens*), gorilla (*Gorilla gorilla)*, Bornean orangutan (*Pongo pygmeaus*), Sumatran orangutan (*Pongo abelii*), and siamang (*Symphalangus syndactylus*), as described in (61). In short, motifs of A-phased repeats (APRs), direct repeats, mirror repeats, short tandem repeats (STRs), and Z-DNA were annotated in each T2T genome with the software gfa (7) (https://github.com/abcsFrederick/non-B_gfa) with the flag *–skipGQ*. This predicts APRs with at least three A-tracts of length 3-9 bp and 10-11 bp between A-tract centers (it also looks for T-tracts that correspond to APRs on the reverse strand); direct repeats with unit lengths 10-300 bp and a maximum spacer length of 100 bp (we note that no DR spacer was longer than 10 bp, Fig. S1); inverted repeats with arms of 6 bp or longer (no upper cutoff) and a maximum loop size of 100 bp; mirror repeats with arms of 10 bp or longer (no upper cutoff) and a maximum loop size of 100 bp; STRs with repeated units of size 1-9 bp and a total length of at least 10 bp (no upper cutoff), and Z-DNA motifs (alternating purine-pyrimidine nucleotides) longer than 10 bp (no upper cutoff)). Triplex motifs were extracted from the mirror repeats (‘grep subset=1’, default parameters of minimum purine/pyrimidine content of 10% and maximum spacer length 8 bp were used to define the subset). G4s were annotated using Quadron (62) with default settings, which predicts standard G4s with at least four GGG-stems without bulges. The output from each motif type was converted to bed format, and any overlapping annotations of non-B DNA motifs of the same type were merged with mergeBed from bedtools v 2.31.1 (63). For G4s, motifs without scores were omitted from the analysis (less than 0.008% of the total; these were all located closer than 50 bp to a chromosome end, with flanks too short for Quadron to calculate a score). We did not include i-motifs as a specific non-B motif in this study because they are complementary to G4 motifs, and a test run with the tool iM-seeker (64) resulted in >99% overlap with Quadron annotations. Overlap between different motif types was retrieved using bedtools and in-house scripts (see github link below). For spacer length analysis, the spacers were extracted from the raw gfa output files.

### Alignments to previous assembly versions

Each of the T2T assemblies (CHM13v2.0 and v.2 assemblies for non-human ape genomes as available in (61)) for which there was an older non-T2T genome available, was mapped to its older counterpart (panPan3 for bonobo, panTro6 for chimpanzee, hg38 for human, gorGor6 for gorilla, and ponAbe3 for Sumatran orangutan) using winnowmap v2.03 (65). We followed the winnowmap recommendations and first generated a set of high-frequency *k*-mers with meryl v1.4.1 (66) using *k*=19.

Regions that did not map to the old assembly were extracted using bedtools complement, and assigned as ‘new’ (note that newly added regions that are duplicates of previously assembled sequence, e.g., previously unresolved multi-copy genes, repetitive arrays. etc., can align in a many-to-one fashion and will not be considered new). Densities of non-B motifs in ‘new’ and ‘old’ sequences (i.e., sequences in T2T genomes that did not align vs. aligned to the older assembly versions, respectively) were extracted with bash and awk scripts, and fold enrichment was calculated as density in ‘new’ divided by density in ‘old’. The numbers of non-B annotated base pairs in new and old sequences were compared with a chi-square goodness of fit test for each non-B motif type and chromosome type separately, and Bonferroni-corrected for multiple testing. To assess the results’ robustness, we randomly resampled half of the data 10 times and repeated the chi-square goodness of fit tests.

### Enrichment at functional regions

Gene and functional annotations for human (CHM13v2.0) were taken from (67). In short, these consist of gene annotations from NCBI (including enhancers from previous RefSeq versions), origins of replication from (9), and CpG islands annotations from the UCSC Table Browser. Promoters were defined as 1 kb upstream of gene transcription start sites, excluding overlaps with other genes. We considered G4s annotated on both strands. For the other non-B DNA types, the annotations are the same for both strands. Fold enrichment was calculated as non-B motif density for each region divided by the genome-wide non-B DNA density. Many regions were too large to perform random permutations and construct a background distribution for statistical comparison. Therefore, we assessed significance by randomly downsampling the data to either 10% (100 or 20 times) or 50% (20 times). The fold-enrichment was calculated for each subsample, and confidence intervals were constructed. When using 100 subsamples, the two extremes on each side of the distribution were removed. Since G4s are more likely to form in GC-rich regions, we corrected the enrichment in this motif category by multiplying it by a correction factor, following an approach used in (68).

### Enrichment at repetitive sequences and methylation analysis

RepeatMasker annotations were downloaded from https://github.com/marbl/CHM13 (for human version CHM13v2.0) and from https://www.genomeark.org/t2t-all/ (for all other apes). For human, also manually curated repeat annotations of new satellites and composite repeats (69) were downloaded from the same source and analyzed separately.

RepeatMasker output was converted to bed format and labeled according to the repeat class for all repeats except satellites, where both the class and the specific names were used. The repeats were intersected with each non-B motif type separately using bedtools, and non-B density in each repeat class was compared to the genome-wide density using python and bash scripts (provided on the github). We also calculated the fold enrichment in repeats compared to non-repetitive sequences, extracted using bedtools complement. When assessing significance for each non-B DNA motif type in repetitive versus non-repetitive sequence using a chi-square goodness of fit test, all comparisons were significant due to the large number of bases analyzed. Instead, we used the same downsampling approach described above for functional regions, taking 100 (or 20) random subsamples containing 10% (or 50%) of the data. If the distribution of subsamples (or 96% for the set with 100) did not overlap with 1, the comparison was considered significant. Methylation data for the H002 cell line, mapped onto CHM13v2.0, were downloaded from https://s3-us-west-2.amazonaws.com/human-pangenomics/T2T/CHM13/assemblies/annotation/regulation/chm13v2.0_hg002_CpG_ont_guppy6.1.2.bed, and methylation data for the CHM13 cell line were downloaded from <u>chm13v2.0_CHM13_CpG_ont_guppy3.6.0_nanopolish0.13.2.bw</u> (70). These files contain methylation scores for CpG sites, given as a fraction of methylated reads (0 means no methylation was detected in any reads, and 1 means all reads were methylated). We extracted G4s overlapping (partially or fully) with repeat classes that had shown an enrichment for G4s in the above analysis and compared the distribution of methylation scores within the G4 motifs with the distribution of methylation scores from all annotated repeats of each repeat class. Previously published experimental BG4 peak data indicating G4 formation in two cancerous cell lines (71) was translated from hg38 coordinates to CHM13 coordinates using the LiftOver tool (72). To assess to what extent the repeats and satellites with G4 enrichment were assembled in the hg38 version, we lifted over each region from CHM13 to hg38, and summed up all unique regions on anchored chromosomes (meaning if multiple repeat copies in CHM13 mapped to a single region in hg38, this was only counted once, and we did not consider hits on unplaced (“Un”) or random scaffolds as these were not used in (71)). To investigate the discrepancy in predicted G4s for the Walusat repeat in humans between our study and (69), we repeated their G4Hunter analysis (through the web application (73), https://bioinformatics.ibp.cz/#/analyse/quadruplex) on chr14:260778-634253 using default settings and downloaded the resulting G4 hits as a csv file. As G4Hunter only reports results for 25-nt windows, we parsed the Walusat fasta sequence and split it at a commonly occurring pattern (GGGGTCA, chosen so that the sequences start with the longest stretch of guanines). This resulted in the majority of sequences of 64 nt (the Walusat repeat length), and a minority of sequences longer than 64 nt (for diverged monomer copies lacking the aforementioned pattern). We cut all sequences at 64 nt, sorted them, and counted how many times each motif occurred. Out of a total of >5,800 copies on chr14, 1,186 shared the most common motif **GGGG**TCAGAGGAATAGAAA**GGG**ACA**GGG**CTGAAGAACACAGGTCGCTGCATTTAGAAAGGAGGC, which was subsequently tested experimentally (see below).

### Experimental validation of G4s in LSAU and Walusat

We aligned all G4 motifs overlapping with the LSAU motif to each other using mafft v7.481 (74), and observed several patterns of G4s. On chromosomes 4 and 10, there were many identical copies of several different motifs, while annotated LSAU regions on other chromosomes had more diverged sequences with many mismatches between the motifs. To group them together, we ran the sequence cluster algorithm starcode (75) separately on each strand. This clustered similar motifs and returned the consensus sequence. We then extracted the average methylation scores for all G4s in the top five clusters and visually inspected the distribution of these scores for each cluster. Three clusters that showed low methylation in combination with fairly high Quadron stability scores (Fig. S2) were selected for experimental validation. Two of them had uniform motifs (**GGGGG**C**GGGGGG**T**GGGGG**T**GGGG**A**GGGGG**CGGTCAGGCGGC**GGGG**T**GGG** with Quadron score 31.44, and **GGG**CGGCTGCA**GGGG**CCC**GGG**C**GGG**C**GGG**CGACGGTGGCGC**GGG** with Quadron score 19.76). The third cluster contained several very similar but not identical sequences, of which only one was chosen for validation (**GGG**T**GGGG**TGT**GGGGG**T**GGGG**A**GGGG**TGGTCAGGC**GGGGG**T**GGG**, Quadron score 31.01). Single-strand oligos were constructed from the above sequences and investigated by circular dichroism (CD), UV absorption spectra, and native polyacrylamide gel electrophoresis (PAGE), as described in (76). In short, CD measurements were performed at 23°C and samples were measured in potassium ion only (110 mM K^+^= 10mM K-phosphate, pH 7 + 95 mM KCl) in 1μM strand concentration, allowing 1 day to form due to many G-blocks. Isothermal difference spectra (IDS) were obtained by calculating the difference between the absorption spectra of the unfolded (1mM Na-phosphate) and folded (110mM K^+^) forms of samples during an increase of ionic strength. Thermal difference spectra (TDS) were calculated as the difference of the unfolded (95°C) and folded (20°C) forms from temperature dependences in the potassium environment. Temperature dependencies were measured repeatedly (up-down-up-down) one day after K^+^ addition. PAGE was run in 110 mM K^+^ (10mM K-phosphate, pH 7 + 95 mM KCl) at 23°C. Samples were prepared either immediately (loaded on the gel after adding K^+^) or 24 hours before loading onto the gel.

### Non-B distribution along the chromosomes and enrichment at centromeres

The density of each non-B motif type along the genome was calculated in 100-kb non-overlapping windows generated with bedtools makewindows. For heatmap visualization of acrocentric chromosomes, the Y chromosome, and centromeres, such densities were normalized by the highest value to the scale from 0 to 1. Centromeric regions were taken from the GenomeFeatures tracks downloaded from https://www.genomeark.org/ for each species and converted to bed format. For chromosomes with two or more annotated active centromere (‘CEN’) regions with a satellite (‘SAT’) in between, we combined the active centromeres for the enrichment analysis without including the intermediate regions. Fold enrichment was calculated as non-B motif density within centromeres divided by the genome-wide non-B DNA density. No GC correction was performed as the centromeres are large regions with GC content very similar to the genome-wide average (Fig. S3). To test for significance of non-B DNA enrichment at the centromeres, the non-centromeric parts of each chromosome were divided into 100 windows with the same size as the actual centromere (for most chromosomes, the windows had to overlap, however, if there were more than 100 possible non-overlapping windows, they were chosen randomly). Then, non-B DNA fold enrichment was calculated for each window separately, and the 100 values obtained for each chromosome were used as a null distribution against which we compared the centromere enrichment. If the centromere fell outside the 0.025th and the 0.975th quantiles, the enrichment was considered significant. For detailed figures of centromeric and acrocentric regions, tracks with centromeric satellite repeats were downloaded from GenomeArk and added using the color scheme from the UCSC Genome Browser(77). This included annotations of active alpha satellite (αSat, the parts that associate with the kinetochore proteins, usually the longest HOR array on each chromosome), inactive αSat (HOR arrays that do not associate with the kinetochore), divergent αSat (older HORs that have started to erode), and monomeric αSat (repeats not organized into HORs) (78). CENP-B binding motif annotation files were downloaded from GenomeArk (for non-human apes) and the supplementary database S15 in (78). Suprachromosomal family information was extracted from the centromeric satellite annotation.

Circular density plots were generated with Circos (79). Figures, as well as all statistical tests, were generated in R v4.4.0 (80) using the tidyverse (81), ggupset (82), patchwork (83), cowplot (84), ggtext (85) and ggh4x (86) libraries.

## Results

### Non-B DNA annotations

Most non-B DNA motifs—A-phased, direct, inverted, and mirror repeats, short tandem repeats, and Z-DNA—were annotated in the latest versions of human and non-human ape T2T genomes (61) with gfa (7). Some previous studies suggested that non-B DNA folds at inverted repeats only when the spacer length is below 15 bp (87–89); however, this has been debated. We used the default parameters for the gfa annotations that allow for longer spacers (up to 100 bp) because of this uncertainty and because most of our annotations had spacers below 15 bp (Fig. S1). We have also annotated a subset of mirror repeats with a high potential to form triplexes (i.e., mirror repeats comprising predominantly purines or pyrimidines and having a short spacer, see Methods) using gfa. G4s were annotated with Quadron (62).

### An overrepresentation of non-B DNA motifs at the newly added regions of the human T2T genomes

We observed an overrepresentation of most non-B DNA motif types at the newly added sequences of the T2T human genome (CHM13) as compared to the previous, non-T2T version (hg38; Table 1), demonstrating the power of T2T genome assemblies in resolving these motifs. In particular, A-phased, direct, inverted, and mirror repeats, as well as STRs, were substantially overrepresented at the newly added sequences for the autosomes. Direct, inverted, and mirror repeats, triplex motifs, G4s (although this was not significant after correcting for multiple tests), and STRs, were overrepresented at such sequences for the X chromosome, and inverted and mirror repeats were overrepresented for the Y chromosome. Similar results were previously obtained for great ape sex chromosomes (59, 60) and autosomes (61).

**Table 1.**
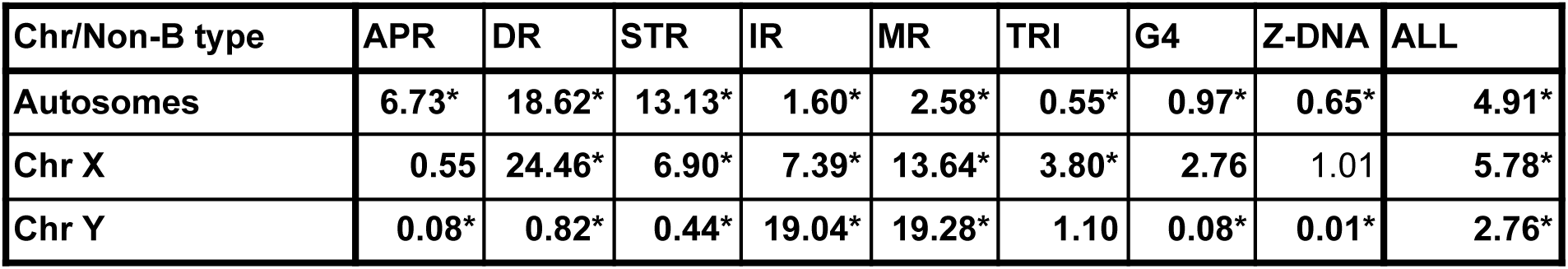
Non-B DNA motifs enriched at the newly added sequences of the human T2T genome (v2.0.CHM13+Y) as compared to hg38, shown as fold enrichment for unaligned vs. aligned sequences. Bold numbers indicate significantly different non-B DNA content in aligned vs. unaligned sequences (chi-square goodness of fit test with Bonferroni correction for multiple testing, *P*<0.01. Cells marked with ‘*” remained significant after the data were randomly subsampled down to half in 10 independent runs, see Methods). APR: A-phased repeats, DR: direct repeats, STR: short tandem repeats, IR: inverted repeats, MR: mirror repeats, TRI: triplex motifs (a subset of mirror repeats), G4: G-quadruplexes.

| Chr/Non-B type | APR | DR | STR | IR | MR | TRI | G4 | Z-DNA | ALL |
| --- | --- | --- | --- | --- | --- | --- | --- | --- | --- |
| **Autosomes** | **6.73\*** | **18.62\*** | **13.13\*** | **1.60\*** | **2.58\*** | **0.55\*** | **0.97\*** | **0.65\*** | **4.91\*** |
| **Chr X** | **0.55** | **24.46\*** | **6.90\*** | **7.39\*** | **13.64\*** | **3.80\*** | **2.76** | 1.01 | **5.78\*** |
| **Chr Y** | **0.08\*** | **0.82\*** | **0.44\*** | **19.04\*** | **19.28\*** | **1.10** | **0.08\*** | **0.01\*** | **2.76\*** |

### Distribution of non-B DNA motifs between sex chromosomes and autosomes, and among species

The non-B DNA motif annotations in the T2T ape genomes revealed the complete picture of the distribution of these motifs among different chromosome types, non-B DNA types, and species (Fig. 2 and Tables S1-S2).

**Figure 2.**
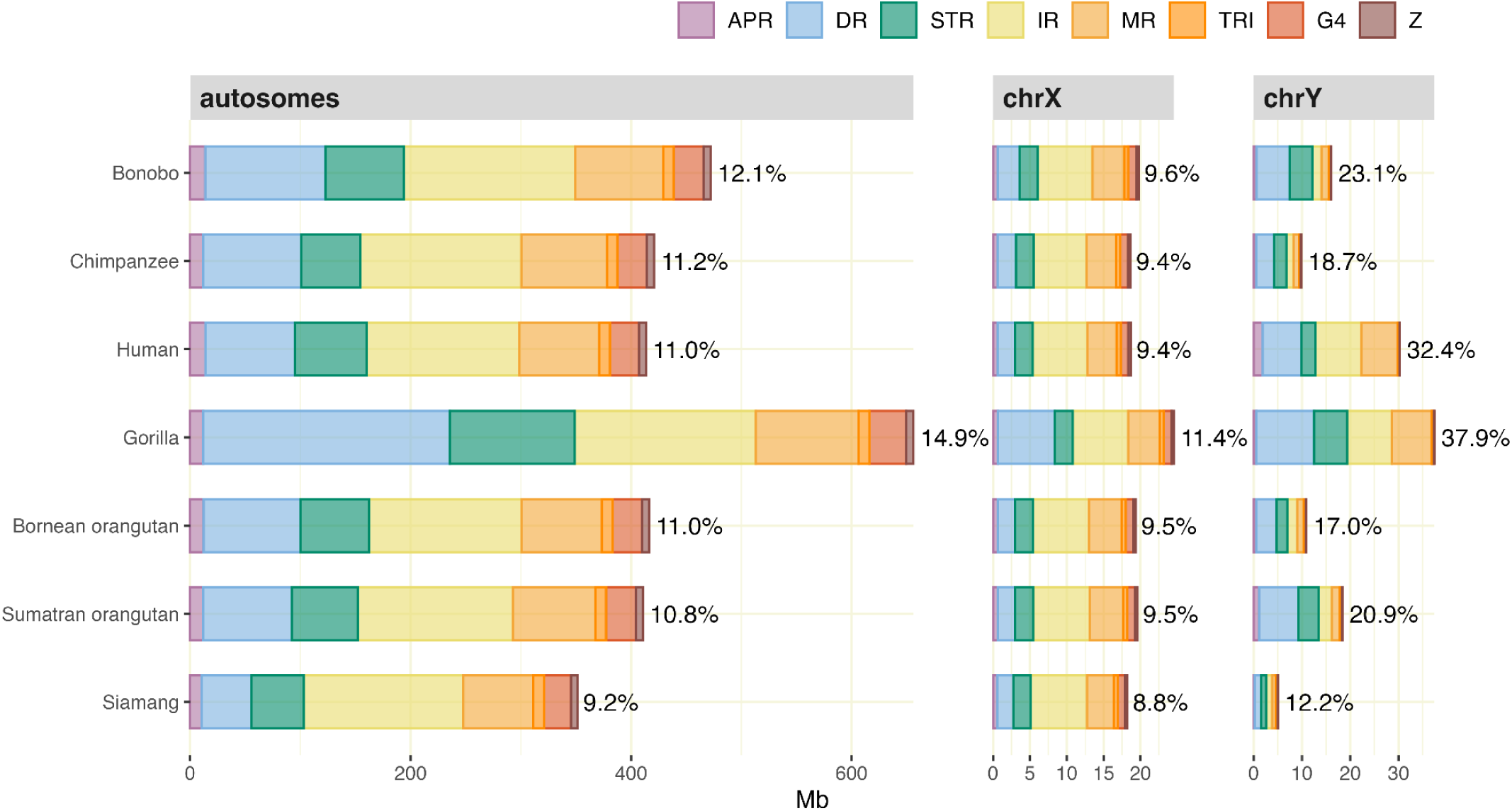
Non-B DNA motif annotations in T2T ape genomes (in Mb and percentage of total genome length), shown separately for autosomes and sex chromosomes. APR: A-phased repeats; DR: direct repeats; STR: short tandem repeats; IR: inverted repeats; MR: mirror repeats; TRI: triplex motifs; G4: G-quadruplexes; Z: Z-DNA. Note that the scale on the X-axis is different for each chromosome type. TRI is a subset of MR, which is plotted on top of MR. The data for this figure are in Table S1.

Depending on species, autosomes had 9.2-14.9% of their sequence annotated in non-B DNA motifs. The X chromosome had a lower percentage, and the Y chromosome had a higher percentage than that for the autosomes in each species (ranging across species from 8.8-11.4% for the X and from 12.2-37.9% for the Y). The number of predicted motifs ranged from 411,000 to 8 million, depending on non-B DNA type and species (Table S2). As a rule, inverted, mirror, direct repeats, and STRs were more abundant than the other non-B DNA motif types. G4s occupied 0.8-1.0% of autosomal sequences and usually a lower percentage of sex chromosomal sequences. A-phased repeats, triplex motifs, and Z-DNA each occupied a lower percentage of autosomal sequences than G4s. Among the species analyzed, the gorilla genome had the highest percentage of non-B DNA motif annotations, whereas siamang had the lowest. Some non-B DNA motif types were distinctly more abundant in some species than others. For instance, across chromosome types, direct repeats were more abundant in gorilla than in other species.

### Overlapping annotations of different non-B DNA motif types

We found substantial overlap among non-B DNA annotations of different types (Fig. 3, Fig. S4), suggesting alternative structure formation afforded by the same genomic sequence. For instance, ∼70%, ∼55%, and ∼48% of Z-DNA annotations on the human autosomes overlapped with STR, mirror repeats, and direct repeat annotations, respectively (Fig. 3B). The amount and types of motifs that overlapped differed between autosomes and sex chromosomes. For autosomes, the largest overlap was found between direct repeats and STRs, followed by the overlap between these two types and mirror repeats. For the Y chromosome, the largest overlap was found between mirror and inverted repeats, with overlapping annotations spanning more bases than non-overlapping annotations for these motifs (Fig. 3A). Non-human apes showed patterns of overlap similar to those observed in humans for both the autosomes and the X chromosome. However, differences were observed for the Y chromosome (Fig. S4A-L). For example, the overlap between mirror and inverted repeats on chromosome Y was less pronounced in non-human apes than in humans. This observation appears to be driven by species-specific patterns of satellite content on the Y chromosome (see next section).

**Figure 3.**
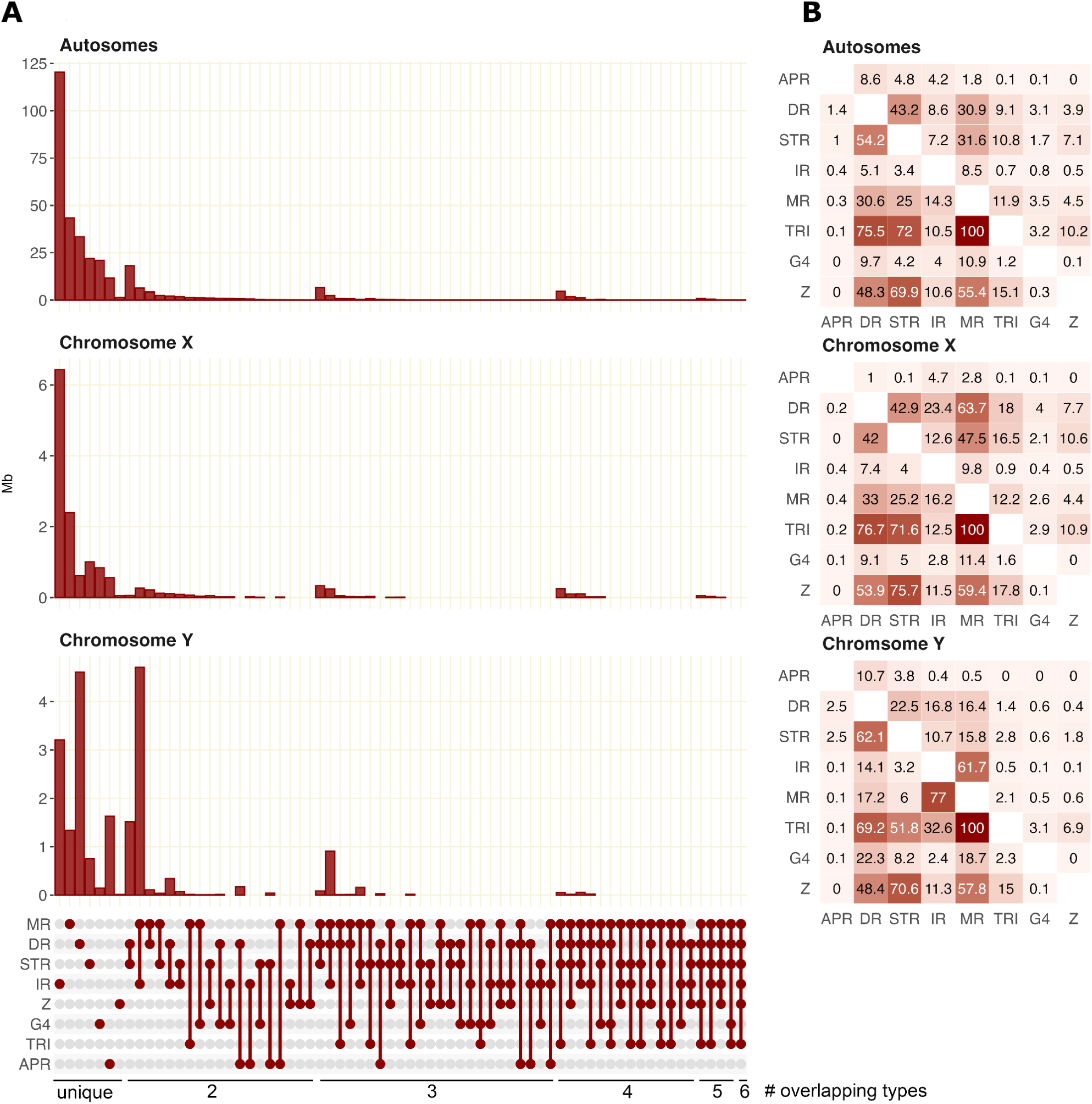
Non-B DNA motif types annotations and their overlaps (i.e., the same bases annotated) in human T2T autosomes, chromosome X, and chromosome. **Y.** (**A**) Number of Megabases and upset plot, comprising all combinations with a total overlap >10 kb, (**B**) Pairwise overlap given as the percentage of the row type (indicated on the left) that overlaps with the column type (indicated at the bottom). APR: A-phased repeats; DR: direct repeats; G4: G-quadruplexes; IR: inverted repeats; MR: mirror repeats; TRI: triplex motifs; STR: short tandem repeats; Z: Z-DNA. Note that TRI overlaps with MR to 100% as the former is a subset of the latter. See Fig. S4 for non-B DNA motif type annotations and overlaps in the other species.

### Distribution of non-B DNA motifs along the chromosomes: General trends

A visual inspection of the density of non-B DNA motifs along ape chromosomes (Fig. 4, Fig. S5, and Fig. S6) suggested the following trends. In humans, all non-B DNA motif types have high density on the short arms of acrocentric chromosomes (chromosomes 13, 14, 15, 21, and 22), and A-phased, direct, short tandem, inverted, and mirror repeats have high density in the heterochromatic region of the Y chromosome (Fig. 4C). We note that triplex motifs do not show the same high-density peaks as mirror repeats, despite being a subset of the latter. A >25-Mb region on the q-arm of human chromosome 9 shows high enrichment of A-phased, direct, short tandem repeats (Fig 4C); this corresponds to a large HSat3 satellite that was missing from the previous human reference genome (hg38). The acrocentric chromosomes in non-human great apes showed similarly high densities of non-B DNA motifs on the short arms, especially for direct repeats and STRs in gorilla and orangutans (Fig. 4D-F). The patchwork of different non-B motifs corresponded very well to the centromeric satellite repeat annotation, with, for example, HSat1 enriched in inverted and mirror repeats and HSat3 enriched in A-phased, direct, and short tandem repeats (see Fig. S7A-F for detailed figures of the acrocentric chromosomes, Fig. S8 for the Y chromosome, and Fig. S9 for enrichment per satellite). rDNA was enriched in all non-B types except A-phased and inverted repeats. Subtelomeric regions were frequently enriched in G4s. An interesting pattern was observed at and around the centromeres, with some centromeres showing high densities for certain non-B DNA motifs, while others having higher densities in the flanking regions. We investigated some of these patterns in more detail below.

**Figure 4.**
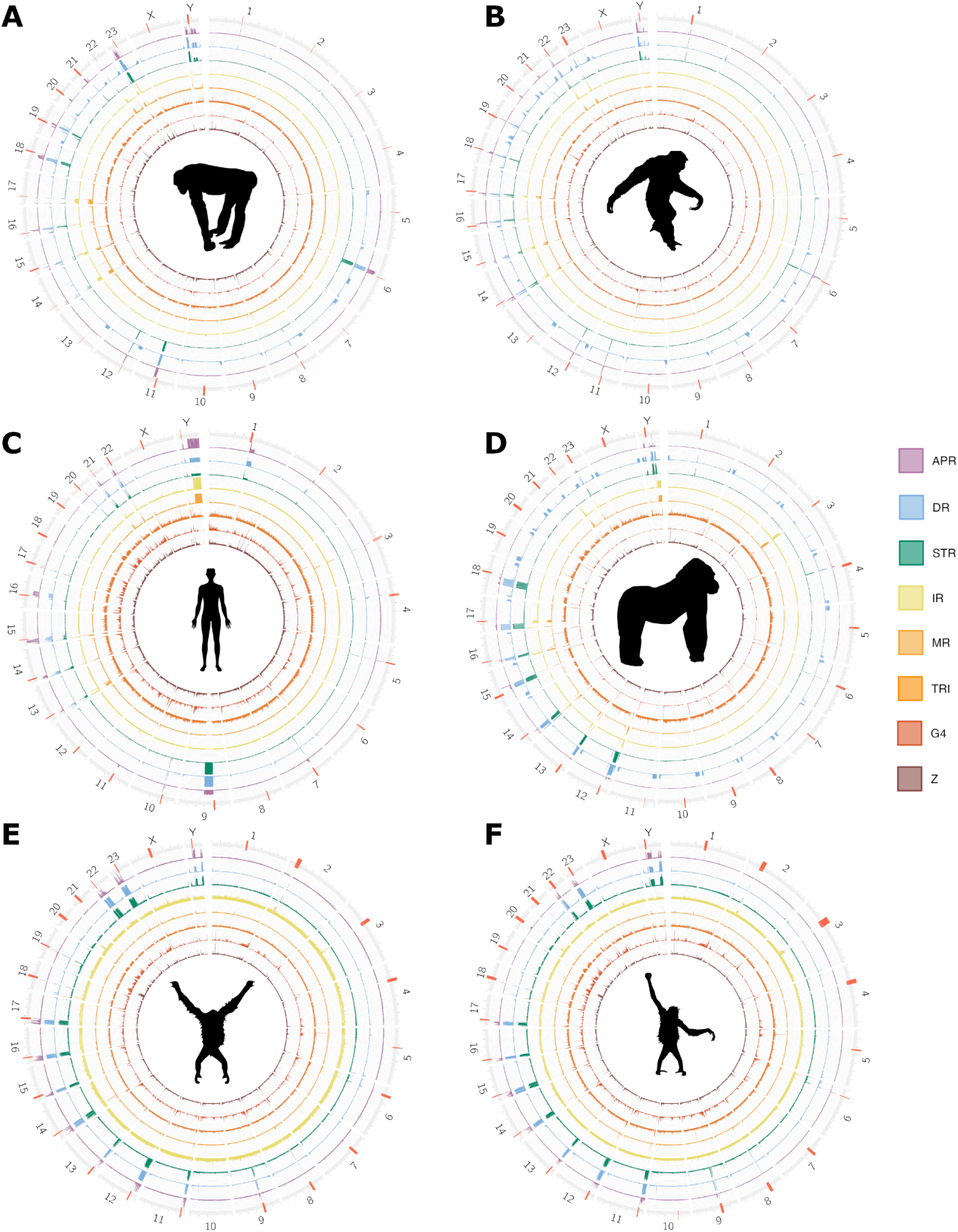
Non-B DNA density along the primary haplotype chromosomes for (A) chimpanzee, (B) bonobo, (C) human, (D) gorilla, (E) Bornean orangutan, and (F) Sumatran orangutan. See Fig. S5 for non-B DNA density in siamang. Active centromeres are marked with red stripes along the chromosomes. Note that an active centromere was not found for chr10 in Bornean orangutan. Abbreviations for non-B DNA are as in Fig. 2. The alternative haplotypes are shown in Fig. S6. Animal silhouettes are from https://www.phylopic.org.

### Enrichment of non-B DNA motifs at genes and regulatory elements

To perform a more rigorous analysis of non-B DNA enrichment, we evaluated it in different functional regions, repeats (based on RepeatMasker annotations), and the remaining, presumably non-functional non-repetitive regions of the human genome (similar analyses were not performed in non-human ape genomes due to incomplete annotations of functional sequences). Many types of non-B DNA were previously implicated in the regulation of transcription (see references in the Introduction). Consistent with these studies, but now analyzing the complete, T2T human genome, we found enrichment of G4s and Z-DNA at promoters and enhancers, as well as at origins of replication (Fig. 5, Fig S10). 14.4% of the promoters and 4.1% of the enhancers contained both G4 and Z-DNA motifs, but typically not at the same position (only 0.07% of sites were annotated as both motifs). CpG islands were enriched in all types of non-B DNA motifs except for A-phased and inverted repeats. 5’ untranslated regions (UTRs) and, to a smaller degree, 3’UTRs, as well as coding sequences, were enriched in G4s. This was still true after correcting G4 fold enrichment for GC content using a simple conversion factor (see Methods, Fig. S11, and Discussion).

**Figure 5.**
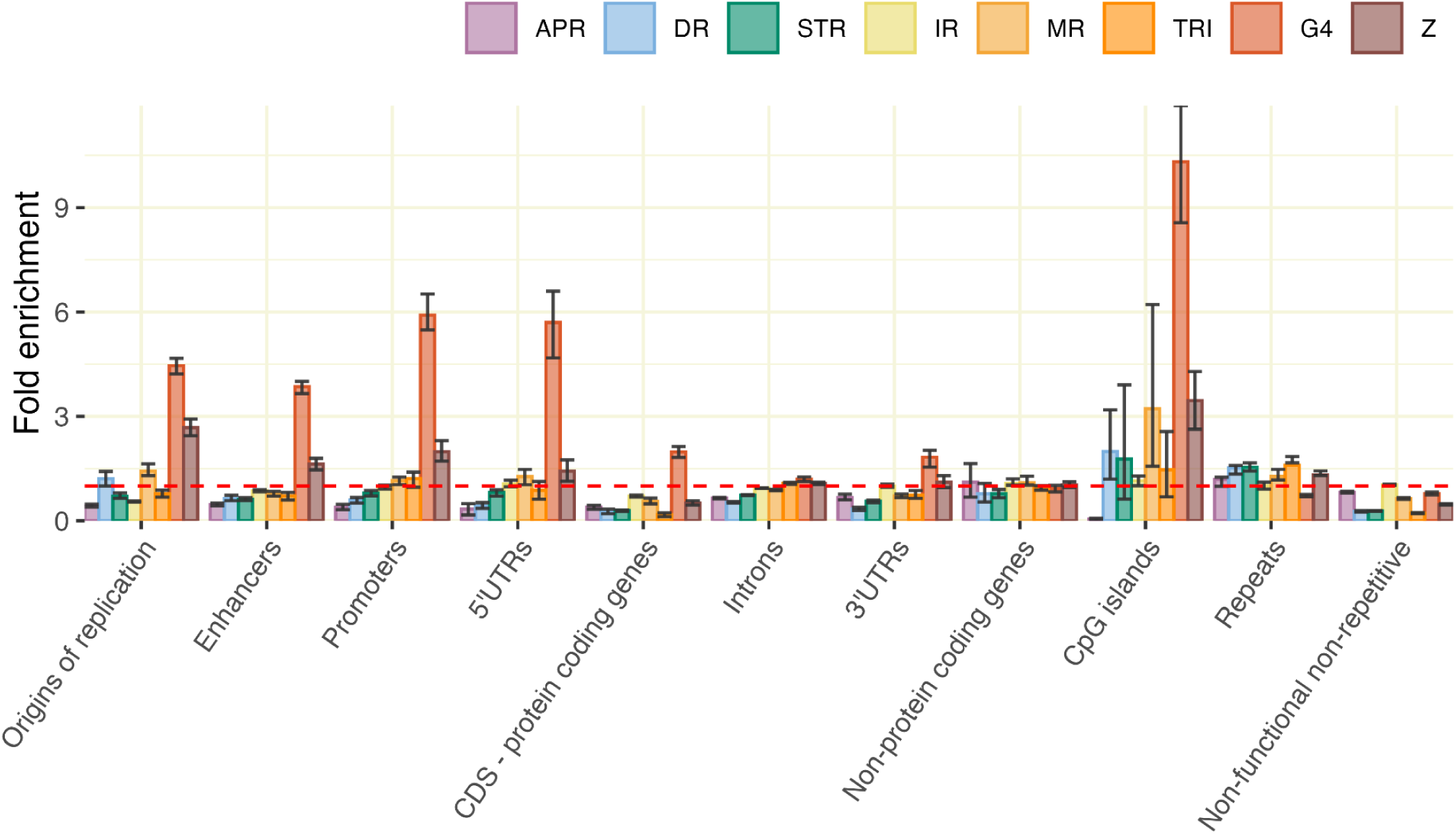
Enrichment of non-B DNA motifs at different functional regions of the genome. Fold enrichment is calculated compared to genome-wide density. Non-functional non-repetitive regions represent sequences that do not belong to the other categories. Red dashed line (fold enrichment=1) represents the genome-wide average. The data were randomly downsampled to 10% 100 times to construct 96% confidence intervals. Bars with confidence intervals overlapping the red line are not considered to be significantly different from the genome-wide average. See Figure S10 for different downsampling approaches. Abbreviations for non-B DNA are as in Fig. 1.

### Enrichment of non-B DNA motifs at repeats and satellites

The T2T genomes provided a complete resolution of repeats in the ape genomes, including transposable elements and satellites, allowing us to comprehensively evaluate non-B DNA present at such genomic regions. In general, repetitive sequences harbored more non-B DNA than non-repetitive sequences (for example, 1.4× more in human, and 2.0× more in gorilla, Table 2, Table S3). Simple repeats and low-complexity regions were strongly enriched in most types of non-B DNA motifs. Considered together, transposable elements were not enriched in non-B DNA (Fig. 6, Table S4). However, some transposable elements showed enrichment in inverted and A-phased repeats, and SVA was enriched in G4s and direct and mirror repeats. Interestingly, RNA as a group was enriched in G4s (Fig. 6A), a signal driven by ribosomal RNA (rRNA, Fig. 6B).

**Figure 6.**
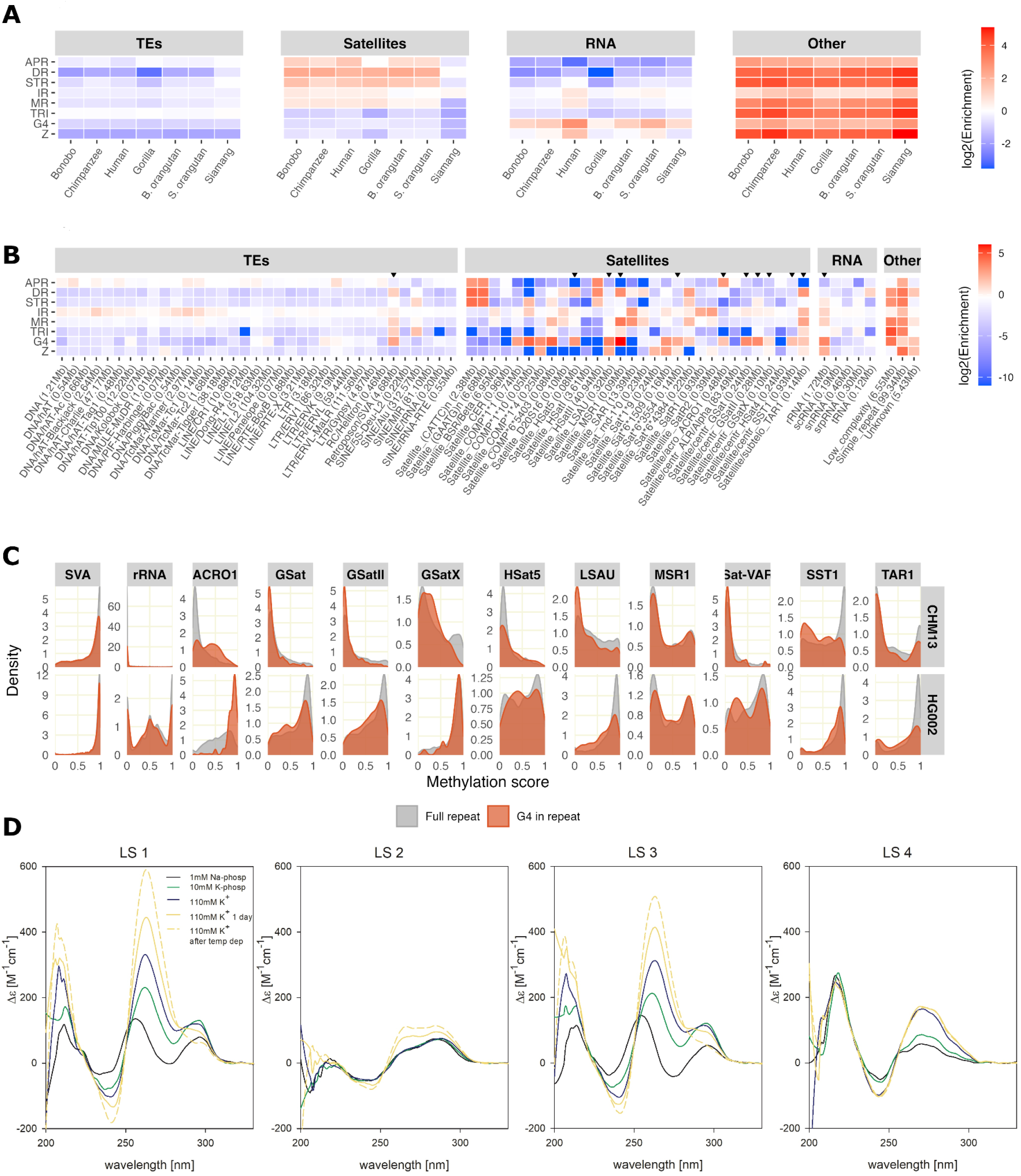
Non-B DNA at repeats and satellites. (**A**) Fold enrichment (given as log-fold densities compared to genome-wide densities) in the four repeat groups for all seven species, centromeric satellite repeats (shown in Fig. S9) are included in the Satellite group. Underrepresentation of non-B DNA (values below 1) is shown in blue, whereas enrichment (values above 1) is shown in red. (**B**) Enrichment of non-B DNA motifs (compared to genome-wide) at repeats, for the human genome. Long repeat names were shortened for visualization purposes (marked with *). Total repeat lengths are given after the names; repeats with a total length shorter than 50 kb are not shown. Abbreviations for non-B DNA are as in Fig. 2. Repeats marked with a black arrow are further investigated in part C. (**C**) Distribution of methylation scores at selected repeats (gray) and G4s (vermilion) overlapping with the repeat annotations, obtained from CpG sites in the human lymphoblastoid cell line (HG002) and hydatidiform mole cell line (CHM13). A score of 1 means that all reads were methylated at this site, whereas a score of 0 means no methylation. (**D**) Circular dichroism spectroscopy of the four experimentally tested G4 motifs: LS1-LS3 from LSAU and LS4 from Walusat. Samples were measured in the potassium chloride buffer (110 mM K^+^= 10mM K-phosphate, pH 7 + 95 mM KCl) in 1μM strand concentration.

**Table 2.**
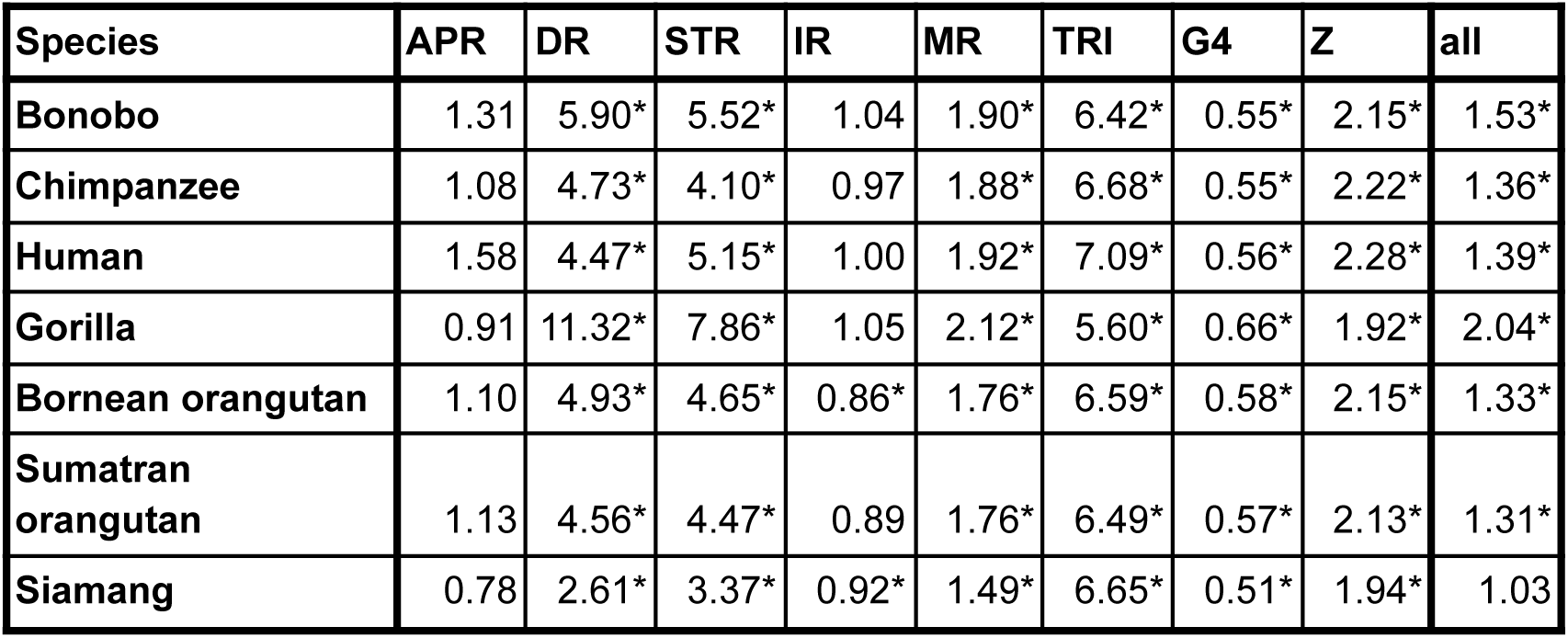
Enrichment of non-B DNA motifs at repetitive sequences. Fold enrichment is calculated as non-B motif density at repeats compared to density in non-repetitive sequence, using only the primary haplotypes. Due to a large number of bases examined, all comparisons are significant in a chi-square goodness of fit test performed as for Table1. Cells for which the 96% confidence interval (based on subsampling the data randomly down to 10% in 100 separate runs) were exclusively enriched or depleted are marked with “*”. See Table S3 for different downsampling approaches. Abbreviations for non-B DNA are as in Table 1.

Considered together, satellites were enriched in A-phased, direct, inverted, and mirror repeats, as well as in STRs, but not in G4s and Z-DNA, for all great apes studied but not for the siamang (Fig. 6A, Table S4). The patterns of non-B DNA enrichment at particular satellites (see Fig. 6B and Fig. S12 for human, and Fig. S13 for non-human apes) were often shared across species. For instance, the LSAU and MSR1 satellites were enriched in G4s in all studied species. The Nereid, Neso, and Proteus satellites were enriched in direct, inverted, and mirror repeats in all studied non-human apes (Fig. S13A-F). The human repeat annotations have been recently updated with manually curated satellites and composite repeats (69). Many of these satellites were enriched for direct repeats. However, surprisingly, we found no enrichment of G4 motifs in Walusat (Fig. S12A), which was reported previously (69). Composite repeats were more often enriched in G4 and Z-DNA motifs than in the other non-B DNA types (Fig. S12B).

We further analyzed a subset of human satellites and repeats enriched in G4 motifs (Table S5). These included retrotransposon SVA, rRNA, and satellites ACRO1, GSat, GSatII, GSatX, HSat5, LSAU, MSR1, Sat-VAR, SST1, and TAR1. Whereas G4 motifs were enriched in these cases, we had no evidence of G4 formation vs. non-formation. As a proxy of such formation, we evaluated methylation profiles (based on modified base calling from Oxford Nanopore Technologies, ONT, sequencing) for G4s and repeats/satellites harboring them in the HG002 lymphoblastoid and the CHM13 hydatidiform mole cell lines (70), as methylation was shown to be antagonistic to G4 formation (90, 91). In the SVA retrotransposon, both cell lines were methylated at G4s as well as at the full repeat region, suggesting that G4 structures do not form at this retrotransposon. In most of the other cases we considered, G4 motifs enriched at repeats and satellites were unmethylated in CHM13 and methylated in HG002, reflecting the overall methylation trends at these genomic regions for these cell lines (70) (Fig. 6C). However, for certain satellites (e.g., LSAU and TAR1 in HG002, and LSAU and SST1 in CHM13), the methylation level for G4s was lower than that for the satellites harboring them, suggesting G4 structure formation. For some satellites (e.g., HSat5 in HG002 and MSR1 for both cell types), we observed a bimodal distribution of methylated vs. unmethylated G4s, suggesting alternative structure formation (i.e., B vs. non-B DNA). Similarly, rRNA and G4 motifs in it were largely unmethylated in CHM13, suggesting G4 formation, and had a bimodal methylation score distribution in HG002. A more direct approach to infer G4 formation is G4-CUT&Tag, which marks G4s using a BG4 antibody before sequencing. Lifting over BG4 peak data from a previous study (71) onto CHM13 suggested G4 formation in ten out of the 12 G4 enriched repeats and satellites (Table S6). We noted that LSAU had the highest number of peaks per sequence length.

We then experimentally validated three G4 motifs found in the LSAU satellite—one of the satellites with the strongest G4 enrichment in our dataset—and the previously reported potentially G4-forming sequence in Walusat (69) (not annotated by Quadron, see Methods). CD spectroscopy, isothermal difference spectra, and thermal difference spectra consistently supported the formation of intramolecular parallel-stranded G4s for two of the three LSAU motifs tested. In contrast, the third LSAU motif formed a hairpin structure. The sequence from Walusat formed canonical B DNA in all assays (Fig. 6D, Fig. S14).

We also investigated the satellite SST1 in more detail, since it was recently suggested to act as the breakpoint in Robertsonian translocations of acrocentric chromosomes 13, 14, and 21 in humans (92, 93). The satellite itself was enriched in G4s (∼3.4-fold higher density than the genome-wide average), with some differences between the satellite subtypes (Table S7). In particular, subtypes sf1, present on acrocentric p-arms, and sf2, present on chromosomes 4, 17, and 19, were enriched in G4 motifs, whereas subtype sf3, present on chromosome Y, lacked such an enrichment. Lower methylation levels within G4s in SST1 compared to the full satellite suggests G4 formation in CHM13 (Fig. 6C). Strikingly, the sequence between the satellite monomers was found to be enriched in non-B motifs, especially in Z-DNA, for all SF subtypes. Notably, this enrichment was most prominent at subtype sf1 (spacer length of ∼135 bp), where it was 97×, 15×, 7×, and 5× for Z-DNA, mirror repeats, direct repeats, and STRs, respectively, compared to the genome-wide average (Fig. S15, Table S7). The presence of non-B DNA motifs at such high density could be involved in the destabilization of these regions (see Discussion).

### Enrichment of non-B DNA motifs at centromeres

We performed a more detailed analysis of the enrichment of non-B DNA at experimentally annotated, active centromeres available for the six great apes in our data set (Fig. 7, Fig. S16, and Fig. S17), as it was suggested that non-B DNA determines centromere formation (35).

**Figure 7.**
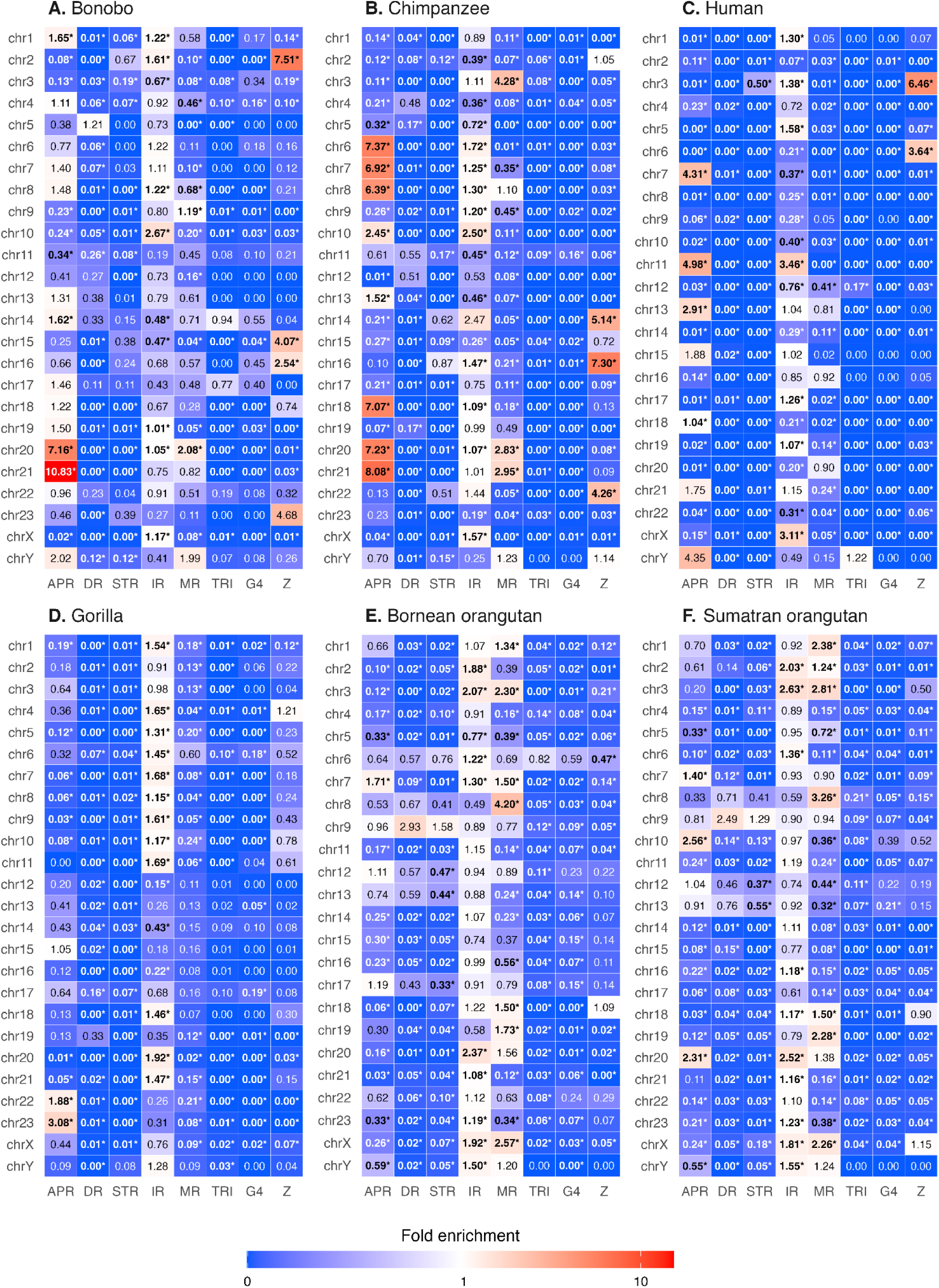
Fold-enrichment of non-B DNA densities in primary haplotype centromeres as compared to genome-wide average densities for (A) bonobo, (B) chimpanzee, (C) human, (D) gorilla, (E) Bornean orangutan, and (F) Sumatran orangutan. The underrepresentation of non-B DNA (values below 1) is shown in blue, while enrichment (values above 1) is shown in red. Densities with a significant underrepresentation or enrichment compared to the genome-wide average density are marked in bold with ‘*’ (two-sided randomization test*, P*<0.05). Abbreviations for non-B DNA are as in Fig. 2. Fold-enrichment for alternative haplotype centromeres can be found in Fig. S16.

Overall, most centromeres (158/263, or 60%) showed significant enrichment in at least one type of non-B DNA, and over a quarter of centromeres (74/263, or 28%) had >2-fold enrichment (Fig. 7, Fig. S16). However, G4s were always underrepresented at centromeres. The other types of non-B DNA displayed species- and chromosome-specific trends. Inverted repeats displayed a moderate enrichment at approximately half of all centromeres. A-phased repeats showed significant enrichment at centromeres of some chromosomes in most species, with a particularly high enrichment and many chromosomes affected in chimpanzee and human.

Z-DNA showed enrichment at centromeres of some chromosomes in bonobo, chimpanzee, and human. In contrast, mirror repeats showed enrichment at centromeres of some chromosomes in the two orangutan species.

Within species, some of these patterns could be explained by suprachromosomal families (SFs) (61, 78). Taken all species together, SF01 centromeres were enriched in Z-DNA, SF1 centromeres were usually enriched in inverted repeats, and SF4—in A-phased repeats (Fig. S18). Some patterns were species-specific. The centromeres in the two orangutan species almost exclusively belonged to SF5, and slightly more than half of them exhibited non-B DNA enrichment (SF1-3 dates after orangutan split from the other great apes (78)). In gorilla, chromosomes with centromeres annotated as SF1 were enriched in inverted repeats, while in chimpanzee and bonobo, SF1 chromosomes showed a mixed pattern of either Z-DNA enrichment, or A-phased, inverted, and/or mirror repeat enrichment (Table S8). The human SF1 and SF2 chromosomes generally lacked non-B DNA enrichment, with a notable exception of chromosomes 13 and 21, which were classified as SF2 and were enriched in A-phased repeats.

When comparing the non-B DNA patterns in centromere regions with detailed centromeric satellite repeat annotation from (61, 78), we observed that the regions of high non-B density often overlapped perfectly with the satellite annotations (Fig. S17), similar to what was observed in the short arms of acrocentric chromosomes and on the Y chromosome. For example, A-phased, direct, and short tandem repeats often occurred at HSat2 and HSat3 satellites, while HSat1A mostly overlapped with inverted and mirror repeats. Inactive α-satellites (αSat, a higher order repeat, or HOR, that does not interact with the kinetochore (78)), divergent αSat (older HORs that have started to degrade), and monomeric αSat (not organised into HORs) were in general not enriched for any type of non-B DNA motifs.

In all species, active centromeres annotated in the primary and alternative haplotypes showed similar non-B DNA enrichment and depletion patterns, with some striking exceptions. In bonobo, the centromere on the primary haplotype of chromosome 17 was significantly enriched for Z-DNA, while such enrichment was missing entirely in the centromere of the alternative haplotype of chromosome 17. We note that the active centromeres on these two haplotypes belong to different suprachromosomal families (SF4 and SF1 for the primary and alternative, respectively). The same pattern of enrichment and depletion of Z-DNA was observed between the centromeres on primary and alternative haplotypes of chromosome 15 in chimpanzee, however this discrepancy cannot be explained by different SF families, as both haplotypes were annotated as belonging to SF1.

Bornean orangutan chromosome 10 lacked an annotated active centromere for the primary haplotype (no evidence of CENP-A enrichment on the alpha satellite HOR array (61)). As there was an annotated centromere for the alternative haplotype, we sought to compare the non-B density in this region between the two haplotypes. However, the alignment of the region between the two haplotypes revealed that the centromere from the alternative haplotype, as well as the 1-Mb upstream flanking region, were entirely missing from the primary haplotype (Fig. S19). The alternative haplotype centromere was depleted of non-B DNA compared to the genome average, but the 1-Mb upstream flanking region showed enrichment in several types of non-B DNA motifs, especially in A-phased repeats, direct repeats, and STRs.

We also sought to compare the non-B density at centromeres across the ape species with and without theCENP-B binding motif, since it has been shown that the lack of CENP-B binding is correlated with increased non-B DNA formation (35). We found little difference in non-B DNA motif content between the 249 centromeres containing the CENP-B binding motif vs. the 23 centromeres that lack the motif (Fig. S20).

However, the 1-Mb flank on the p-arm showed significantly higher enrichment for A-phased, direct, and short tandem repeats in centromeres without the CENP-B binding motif (Fig. S20A). Additionally, the densities of inverted repeats, mirror repeats, and Z-DNA were significantly lower in the 1-Mb flanks of centromeres without vs. with the CENP-B binding motif. Note that for both groups of centromeres, the flanks were depleted in these non-B DNA motifs compared to their average frequency genome-wide. The q-arm flanks showed no significant differences between the two groups (Fig. S20C).

## Discussion

We conducted a detailed analysis of non-B DNA motifs in the T2T assemblies of human and non-human ape genomes, which have recently become available (58–61). Importantly, these genomes have been produced using long-read sequencing technologies, which are less error-prone at non-B DNA motifs than the Illumina short-read technology (56). Additionally, due to the use of long reads and novel assembly algorithms, these genomes have resolved highly repetitive genomic regions, such as long satellite arrays, including complete centromeres. We found an overrepresentation of most types of non-B DNA motifs in the newly added sequences of the human T2T genome, in agreement with previous studies of non-human T2T ape sex chromosomes (60) and autosomes (61). The ability to analyze previously inaccessible regions of the genomes, which are rich in non-B DNA motifs, allowed us to uncover the complete genome-wide repertoire of such motifs in humans, as well as non-human apes whose T2T genomes are available to date. Thus, our study complements earlier studies of non-B DNA motif enrichment in previously sequenced regions of the human genome (e.g., (68)).

### Non-B DNA at satellites and repetitive elements: General trends

We found that, on a large scale, non-B DNA motifs are unevenly distributed among and along ape genomes. In the human genome, many blocks of high-density non-B DNA motifs correspond to satellites, particularly centromeric satellites or satellites at the short arms of acrocentric chromosomes. Since satellites are repetitive structures based on a particular sequence, they could be highly enriched (for example, if a certain non-B motif occurs in the smallest repeat unit repeated hundreds of times in the satellite) or completely depleted (for example, if the smallest repeated unit is A-poor, A-phased repeats will likely not occur in the satellite) in non-B DNA motifs. Nevertheless, of the 38 satellite types shown in Figure 6B, 34 are enriched in at least one of the non-B motif types. Of the four that lack enrichment compared to genome-wide, one (HSat1) is completely interspersed with simple repeats. In the manually curated CenSat annotation (Figs. S7-S9), these simple repeats are incorporated into the HSat1-satellite array, and the combined array shows strong non-B DNA motif enrichment. This shows the importance of manual curation of complex sequence annotation. The remaining three are all centromeric repeats (CER, HSat4 and Alpha), of which the last includes full centromeres (regions in the >1-Mb scale).

We identified several instances of non-B DNA motif enrichment at particular satellites and transposable elements, consistent with previous analyses of non-T2T genomes showing that some types of non-B DNA motifs are present at repeats, and might be propagated through their spreading (51, 94–96). The ability of satellites to form alternative DNA structures, whether within or between monomers, might play an important role in their copy number dynamics and in determining their inter-unit similarity. Therefore, these non-B DNA features should be incorporated into future models of satellite evolution (97). When considered as groups, satellites displayed enrichment in most non-B DNA motif types, whereas transposable elements did not show such an enrichment.

For G4 motifs in particular, we could predict formation based on methylation status (methylation inhibits G4 formation (91)). The methylation data utilized in this study were extracted directly from Oxford Nanopore reads and have the advantage of being genome-wide. Such data have been shown to be highly accurate and correlate well with conventional bisulfite sequencing, albeit with less accuracy close to SNPs and at asymmetric methylated sites (98). However, these potential limitations should not affect our results because we average methylation scores over many bases. In many instances, G4s enriched at transposable elements (e.g., at SVAs) and satellites were methylated, and thus unlikely to form. In contrast to this pattern, G4s were shown to form at the SVA inserted in the *TAF1* gene and affect its expression in patients with X-linked dystonia parkinsonism (99). However, in some instances, G4s were less methylated than the overall satellites they are embedded into, or had a bimodal methylation density distribution. Such G4s should be investigated further as they may have functional significance for the satellites. Moreover, the methylomes of additional cell lines should be added to this analysis.

We acknowledge that using methylation levels as a proxy for G4 formation might be imprecise. Data show that methylation is almost non-existent at G4 forming regions (mean 1% vs 28.4% genome-wide) (91), but whether this implies that there are G4 structures in all regions of low methylation remains to be tested. More direct methods, such as CUT&Tag approaches using G4 binding antibodies (e.g., (71, 91)) or antibody independent methods such as G4Access (100) have been used to show G4 formation in several human cell lines, but have not been tested in CHM13 (the cell line from which the T2T human genome was sequenced). We used G4 formation from one of these studies to show that G4s form at satellites in two cancerous cell lines, but as G4 formation can vary significantly across cell lines (71), this cannot be used as evidence of formation in CHM13 or HG002. Furthermore, these studies have used short-read Illumina sequencing, which is problematic for investigating large satellite repeats. To fully explore G4 formation in complex repetitive regions genome-wide, targeted studies using long-read data will be necessary, potentially with the newly developed Nano-CUT&Tag (101) (though this has not yet been tested with BG4 antibodies) or PDAL-seq (102).

### Non-B DNA at satellites and repetitive elements: Examples

LSAU was one of the satellites with high G4 enrichment in our dataset, and we validated G4 formation in it experimentally. This satellite has previously been shown to have variable methylation levels in apes (103) and speculated to have an effect on gene expression (104). It is also part of the larger repeat complex D4Z4 whose copy number and methylation level are associated with fascioscapulohumeral muscular dystrophy (105). Our *in vitro* experiments confirmed the formation of two out of three tested LSAU G4 motifs showing low methylation in the CHM13 cell line. We note that, compared to the other two motifs tested, the LSAU motif that did not form a G4 *in vitro* showed higher methylation in HG002 and had a lower Quadron stability score (19.31, compared to >31 for the other two, and very close to the threshold of 19 suggested by the Quadron’s authors for discriminating stable and unstable G4s (62)) than the two others motifs tested. This motif also contained many cytosines that the guanines could pair within a hairpin, rather than forming a G4 (see Methods).

We found no enrichment of G4 motifs in Walusat, a satellite that previously has been reported as enriched in this motif (69). This discrepancy might result from the use of different prediction software programs. Quadron only predicts standard motifs with four G3 stems, while Hoyt and colleagues (69) based their predictions on G4Hunter (73), which additionally includes G4 motifs with bulges (i.e., the G3 stem can be interrupted by other nucleotides); this is the G4 type found in Walusat. We repeated the G4 prediction of the Walusat array on chromosome 14 from (69) and found that the G4Hunter scores of the four most common G4 motifs (each occurring >2,500-5,000 times at Walusat occurrences across the genome) were low, i.e., in the range of 1.20-1.32. This is very close to the default threshold (1.2) and below the more stringent threshold of 1.5 suggested to reduce the false discovery rate to below 10% (106). Our experimental validation of the most common Walusat motif resulted in B DNA formation. We conclude that G4 structures are unlikely to form at Walusat.

### Non-B DNA at the short arms of acrocentric chromosomes

We found that short arms of acrocentric chromosomes correspond to a patchwork of different combinations of non-B DNA motif types. For instance, the HSat1 satellite is rich in inverted and mirror repeats, and the HSat3 satellite is rich in A-phased repeats, direct repeats, and STRs. We performed an in-depth investigation of the satellite SST1, which is present in large arrays on the short arms of acrocentric chromosomes and was suggested to be the breakpoint of Robertsonian translocations in humans (92, 93). We discovered that non-B motifs are enriched not only at the annotated SST1 satellites themselves (where G4s have low methylation levels), but also at the sequence between its satellite monomers. The SST1 subtype sf1, present on the p-arms of chromosomes 13, 14, and 21, has a binding site for PRMD9, a recombinogenic protein. It was suggested that the resulting increase in recombination is one of the prerequisites for Robertsonian translocations (93). Here, we show that the spacers between the SST1 satellite monomers are highly enriched in Z-DNA, which is another known inducer for double-strand breaks (107), and that this enrichment is by far highest on the aforementioned acrocentric chromosomes. We hypothesize that this enrichment could play an important role in this type of translocations.

### Non-B DNA at the centromeres

Our examination of active, experimentally defined centromeres in great apes indicated that more than half of them are enriched in at least one type of non-B DNA motifs, particularly A-phased and direct repeats. This extends an earlier study of non-B DNA enrichment at active centromeres of human, African monkey, and mouse (35) to complete chromosome sequences of multiple species of great apes and suggests an important role of non-B DNA structures in defining centromeres. In fact, Patchigolla and Mellone (38) studied fruit fly chromosomes and suggested that satellite repeats occur at centromeres at least in part because they can form non-B structures. Enrichment in non-B DNA motifs and in R-loop formation was also found at oat centromeres (39), arguing that the involvement of non-B DNA in centromere definition and/or function might be conserved across eukaryotes.

Whereas we observed a pattern of non-B motif enrichment at the centromeres, we could not clearly detect a particular non-B DNA type as the dominant feature of centromeres. Instead, many centromeres were annotated as harboring several non-B DNA types. This is consistent with a recent analysis of human centromeres suggesting that alternative non-B structures can form at them, as evident from high ensemble diversity values (108). Centromeres belonging to the same SF often (but not always) shared common patterns of non-B DNA enrichment. We note that the annotation of SF into subtypes was developed for the human genome, and hypothesize that a more detailed annotation of the non-human apes will generate more subtypes of suprachromosomal families and further increase the correlation between non-B DNA and SFs.

We saw no difference in non-B DNA enrichment between centromeres containing CENP-B binding motifs compared to centromeres lacking these motifs. In contrast, the p-arm 1-Mb flanks of centromeres without CENP-B showed enrichment for several non-B motif types. The CENP-B binding motif is highly conserved over many taxa and has been shown to be essential for *de novo* centromere formation on synthetic chromosomes (109). However, it is absent from some centromeres, and Kasinathan and Henikoff (35) suggested that non-B DNA can substitute CENP-B binding motif in them. Perhaps, to define the centromere, non-B DNA does not have to form within the active α-satellite itself, but can instead occur in its close proximity. In fact, a recent study investigating the minimum free energy (MFE) and thermodynamic ensemble diversity as a proxy for secondary structure and stability in human centromeres found highest MFE (indicating low stability and non-canonical structure formation) in both the active centromere itself and the divergent HORs adjacent to it (108). This is consistent with the importance of the pericentromeric regions for keeping the sister chromatids together at meiosis (110), a process possibly mediated by alternative DNA structures.

### Non-B DNA at functional genomic elements

We showed that several functional elements, including enhancers and promoters, are enriched in G4s, consistent with findings in a previous, non-T2T version of the human genome (68) and in other taxa (reviewed in (111)). These regions are often GC-rich, and GC content correlates with G4 motif abundance (see for example (112)). Nevertheless, it is not resolved whether G4 motifs are often found in these regions because they are GC-rich, or whether these regions are GC-rich due to their high G4 density. We showed that the G4 enrichment for most functional elements also remained after a linear GC correction we applied as in (68). However, as the GC-G4 correlation might be non-linear (67), a more sophisticated correction might be required. For instance, Mohanty and colleagues (67) found that coding regions are no longer significantly enriched in G4s after a quadratic correction for GC content. Because the other non-B motif types show relationships with GC content that are neither linear nor quadratic, we had difficulty finding the same GC correction model suitable for all eight non-B motif types investigated in this study.

### Caveats, future studies, and conclusions

One caveat of our study is that the software we used, gfa (7), only predicts non-B DNA motifs with identical arms for direct, inverted, and mirror repeats. On the one hand, this explains why not all satellites are annotated as direct repeats, even though most have monomers shorter than the maximum arm length considered by gfa (300 bp). On the other hand, mismatches in the arms sequence should destabilize the potential formation of non-B DNA. Different types of non-B DNA motif annotations for the same sequences, however, can lead to more stable non-canonical structures. For instance, slipped-strand structures in long sequences of STRs, which in turn contain inverted motifs (e.g., CTGCAG_n_), are known to be stabilized by forming hairpins in the loops (113). Here we included all mirror repeats in our analyses, consistent with several prior studies (2, 60, 108, 114, 115). We note that only a subset of mirror repeats is predicted to form triplex DNA, which we added here as a separate motif type. We did not include i-motifs (116) in our analysis—an additional type of non-B DNA that forms a tetrameric structure similar to that of a G4 but with cytosines instead of guanines—as its own motif type, due to the fact that i-motifs are predicted at the same positions as G4 motifs, but on the complementary strand. Hence the distribution of i-motifs is similar to that of G4 motifs, though we note that these two structures are unlikely to form simultaneously (117).

In the future, more direct experimental studies should be performed to investigate the formation of non-B DNA structures in ape cells and tissues. Such experiments should also elucidate the precise structures these motifs form, particularly when the same sequence is annotated as multiple non-B DNA motif types. Distinguishing these structures can be important, as, for instance, the promoter of the human *c-MYC* oncogene can form either a G4 or H-DNA, which might have different effects on genomic instability in this genomic region (22). Similarly, knowing what particular non-B DNA structures form at satellites can inform their expansion mode.

In conclusion, our new annotations of non-B DNA motifs in complete ape genomes have shown that there is strong but uneven potential for non-B formation along these genomes and among species. This potential is particularly high in the genomic sequences added to the T2T assemblies. We predict the formation of several alternative secondary structures at many genomic locations. Further studies and experimental validation will determine which structures form in any given species and tissue, as well as their effects on cellular processes.

## Data Availability

All code used for running our analyses and all in-house scripts generated for this paper are available on github: https://github.com/makovalab-psu/T2T_primate_nonB, and archived on Zenodo https://doi.org/10.5281/zenodo.14933484. Non-B DNA annotations are available at the UCSC Genome Browser hub for the ape T2T genomes (https://github.com/marbl/T2T-Browser).

## Acknowledgments

We are grateful to Glennis Logsdon, Karen Miga, Jennifer Gerton, Saswat Mohanty, Edmundo Torres-Gonzalez, Karol Pál, and Jacob Sieg for discussions of the results and useful suggestions. Saswat Mohanty provided the annotations of functional regions, based on a script written by Karol Pál. Robert Harris provided code for running winnowmap. Eddie Yong Hwee Loh provided useful explanations for methylation analysis. Glennis Logsdon and Karen Miga provided information on centromere annotations. Matthias Weissensteiner helped with generating Figure 1. This research was supported by the grant R35GM151945 and by the Willaman Chair Endowment Fund from the Eberly College of Science to KDM. Computations were performed at the Penn State Institute of Computational Data Sciences. This work was also supported by the grant 21–00580S from the Czech Science Foundation awarded to EK.

