## Supplemental Information for "Non-canonical DNA in human and other ape telomere-to-telomere genomes"

**Supplementary figures** ... **2**
Figure S1. Spacer length distribution ... 2
Figure S2. Methylation at satellites and G4s ... 3
Figure S3. GC content of the centromeres ... 4
Figure S4. Overlap between non-B motif types ... 5
Figure S5. Circos plots of non-B DNA motif density for siamang haplotypes ... 11
Figure S6. Circos plots of non-B DNA motif density for alternative haplotypes ... 12
Figure S7. Density plot of non-B DNA motif density for acrocentric short arms ... 13
Figure S8. Non-B DNA density plot of Y chromosomes ... 19
Figure S9. Enrichment of non-B DNA at centromeric and acrocentric satellites ... 20
Figure S10. Enrichment of non-B DNA at functional regions ... 21
Figure S11. GC-corrected enrichment of non-B DNA at functional regions ... 22
Figure S12. Enrichment of non-B DNA at manually curated human repeats ... 23
Figure S13. Enrichment of non-B DNA at repeats in non-human apes ... 24
Figure S14. Experimental validation of G4s ... 25
Figure S15. Non-B DNA motifs at SST1 ... 27
Figure S16. Enrichment of non-B DNA at centromeres for alternative haplotypes ... 28
Figure S17. Density of non-B DNA around centromeres ... 29
Figure S18. Enrichment of non-B DNA at suprachromosomal families ... 35
Figure S19. Non-B DNA density at the Bornean orangutan chr10 ... 36
Figure S20. Non-B DNA fold enrichment around centromeres with or without CENP-B binding motif ... 37
**Supplementary tables** ... **38**
Table S1. Non-B DNA annotations in ape genomes ... 38
Table S2. Non-B DNA motif properties ... 39
Table S3. Enrichment of non-B DNA motifs at repetitive sequences ... 40
Table S4. Non-B DNA motif enrichment at repeat groups ... 41
Table S5. Methylation at repeats and satellites enriched in G4 motifs ... 42
Table S6. G4 formation at repeats and satellites ... 43
Table S7. Non-B DNA enrichment at SST1 ... 44
Table S8. Non-B DNA enrichment at the centromeres, suprachromosomal families, and depending on CENP-B box presence/absence ... 45

### Supplementary figures

**Figure S1. Spacer length distribution**
Spacer length distribution for (A) direct repeats (B) inverted repeats, and (C) mirror repeats. The vast majority of spacers are shorter than 15 bp, which has previously been suggested as a limit for the formation of these structures (see main text for references). The cumulative fraction of spacers equal or shorter than the x-axis length is shown as a red dashed line in each figure. For DR, IR, and MR, the percentage of spacers with length 0 is 31%, 17%, and 13%, respectively.

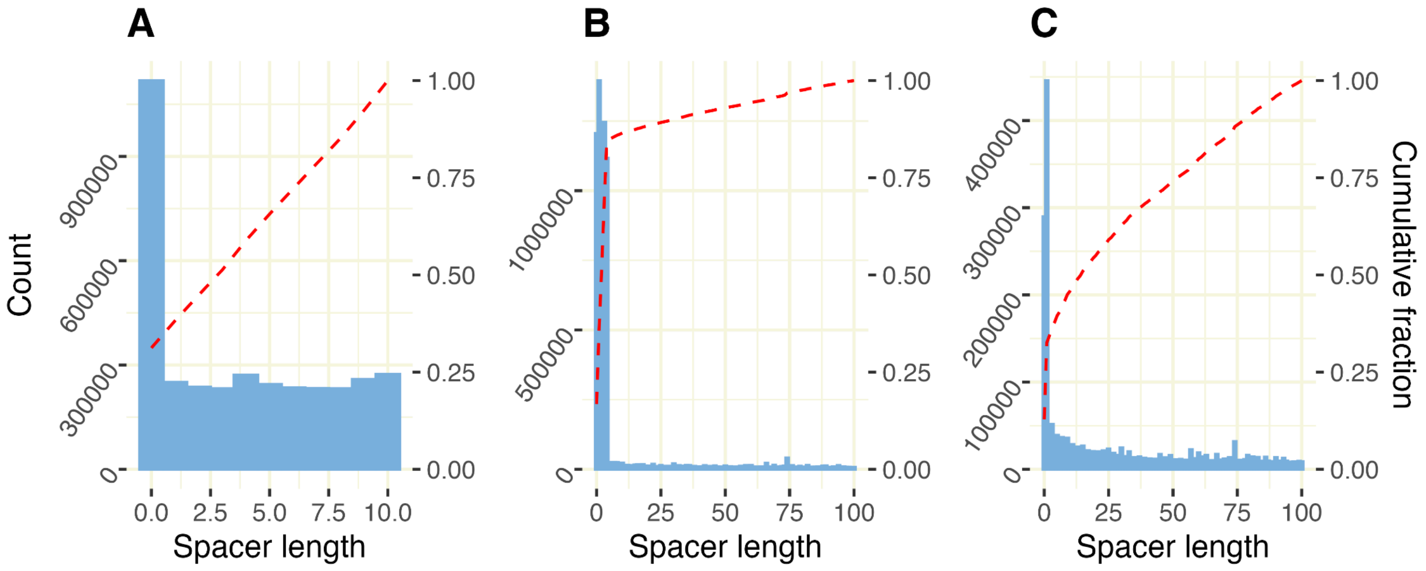

**Figure S2. Methylation at satellites and G4s**

**(A)** Methylation score and **(B)** Quadron score distributions for sequence clusters of G4 motifs in the LSAU satellite. Strand, chromosome, and cluster number are given above each plot. Chromosomes 4 and 10 were run separately as they contained long and uniform stretches of LSAU, and the remaining chromosomes were clustered together (labeled excl4+10). Clusters selected for experimental validation are marked LS1-3, note that two clusters shared identical motifs (marked with LS2).

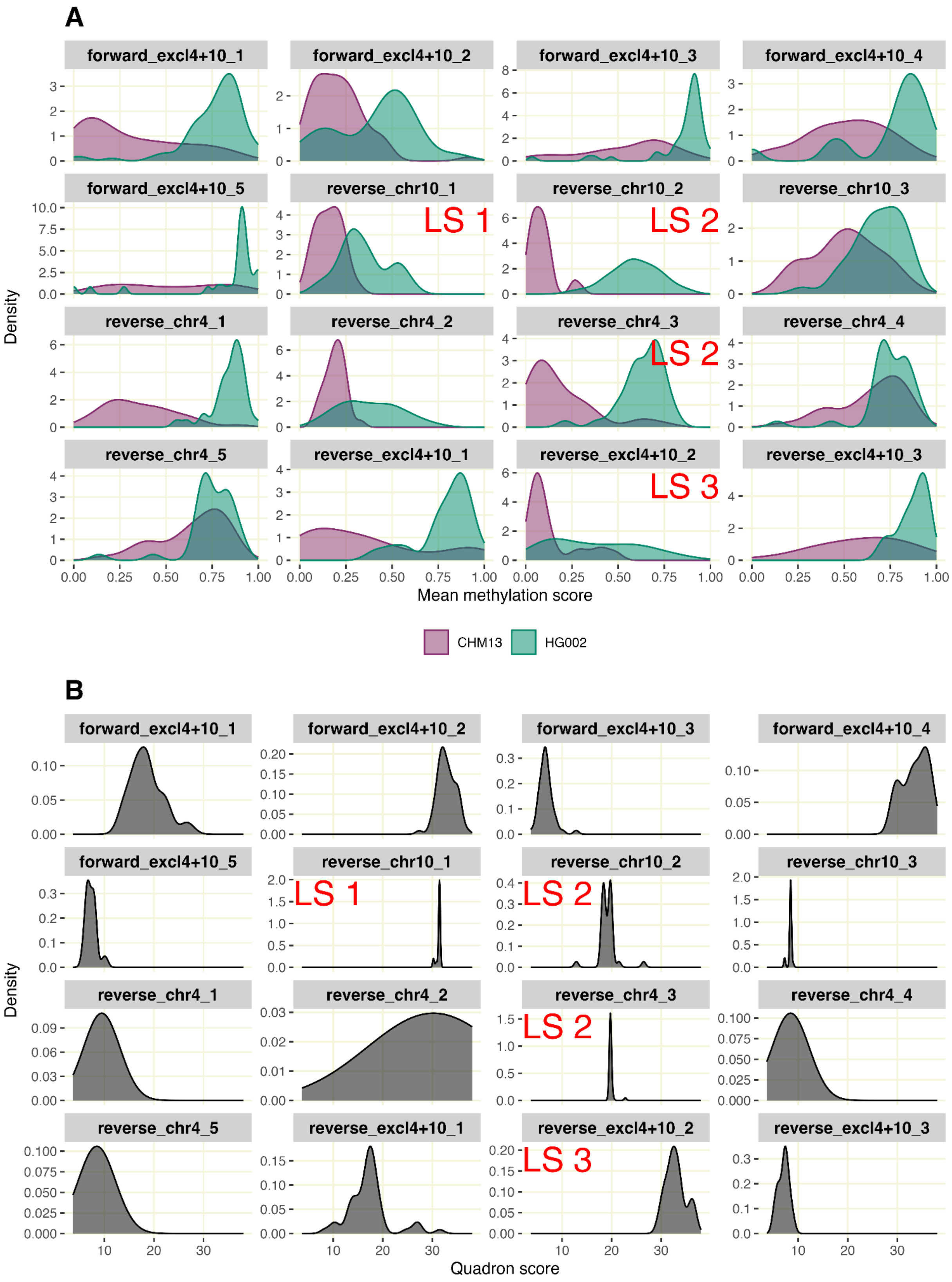

**Figure S3. GC content of the centromeres**
GC content in centromere and background (non-centromere) windows from the primary haplotype assemblies of each species: **(A)** bonobo, **(B)** chimpanzee, **(C)** human, **(D)** gorilla**, (E)** Bornean orangutan, and **(F)** Sumatran orangutan. Window size is determined for each chromosome by the centromere size of that chromosome.

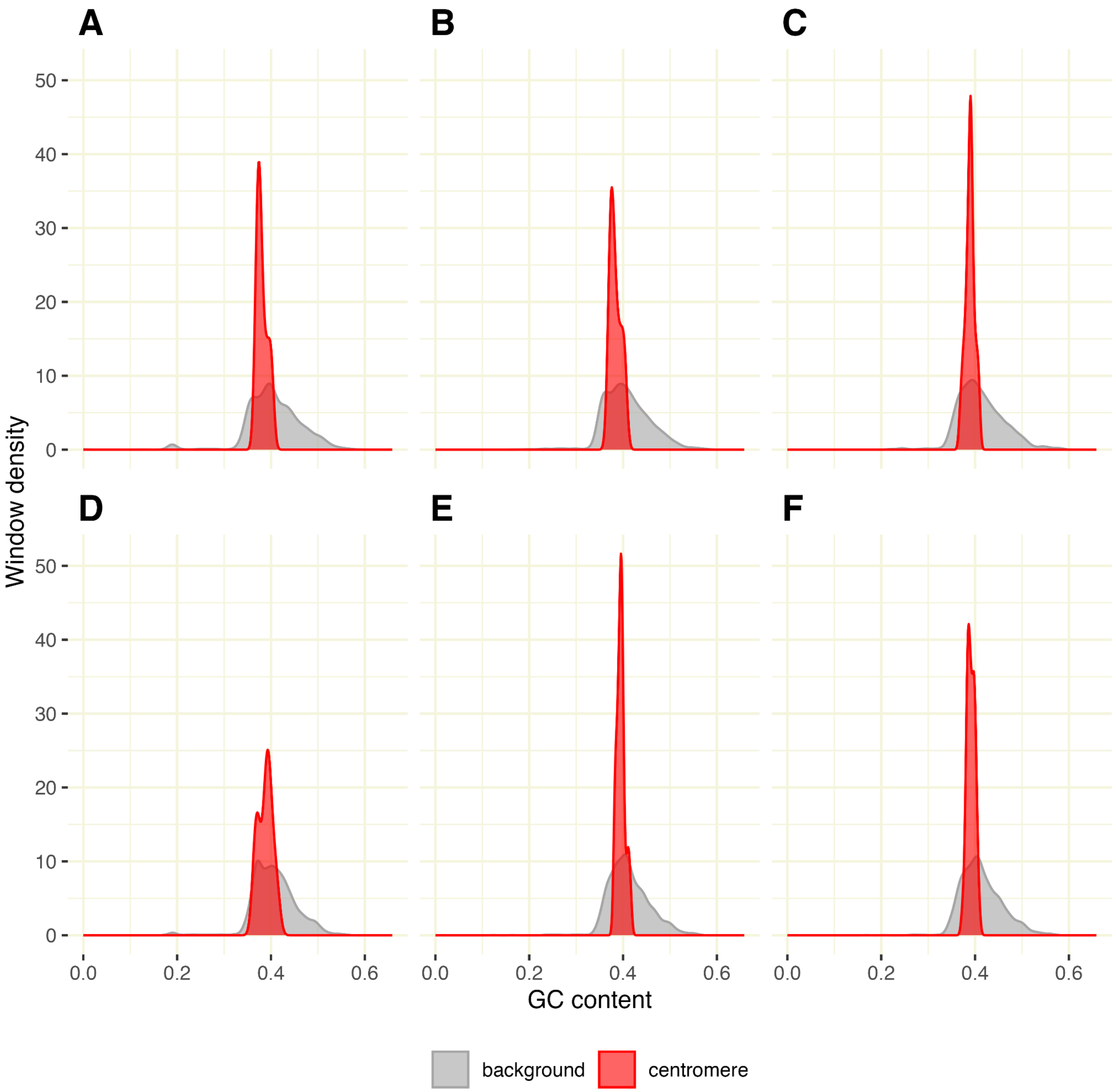

**Figure S4. Overlap between non-B motif types**

Overlap (i.e., the same bases annotated) across different non-B DNA motif types in the non-human T2T genomes on autosomes, chromosome X, and chromosome Y for **(A-B)** bonobo, **(C-D)** chimpanzee, **(E-F)** gorilla, **(G-H)** Bornean orangutan, **(I-J)** Sumatran orangutan, and **(K-L)** siamang. Upset plots with all combinations that have a total overlap of >10 kb are shown in (A, C, E, G, I & K); pairwise overlaps given as the percentage of the left type that overlaps with the bottom type are shown in (B, D, F, H, J & L). The results for human are shown in Fig. 3. Abbreviations: APR: A-phased repeats; DR: direct repeats; STR: short tandem repeats; IR: inverted repeats; MR: mirror repeats; TRI: triplex motifs; G4: G-quadruplexes; Z: Z-DNA.

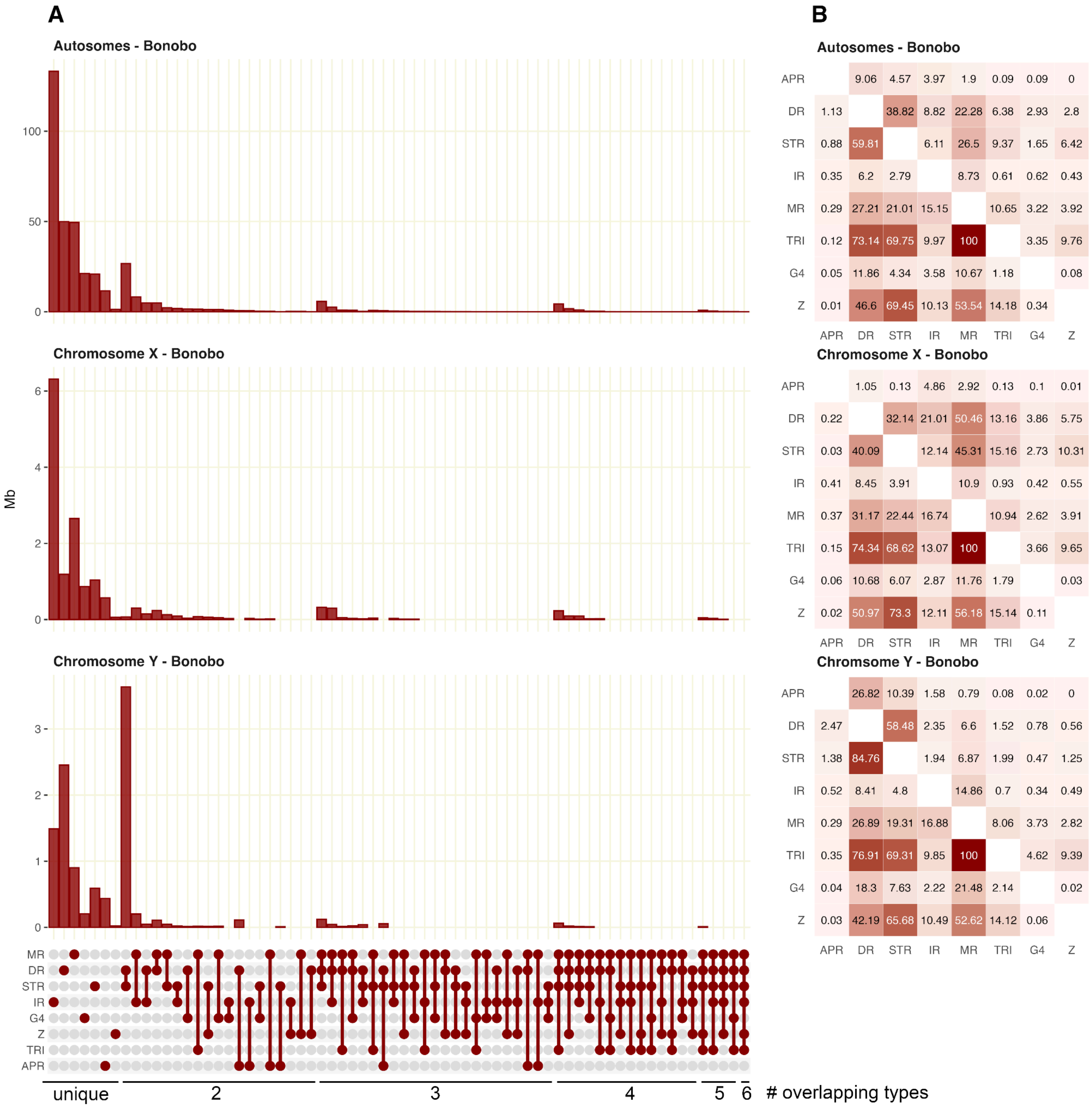

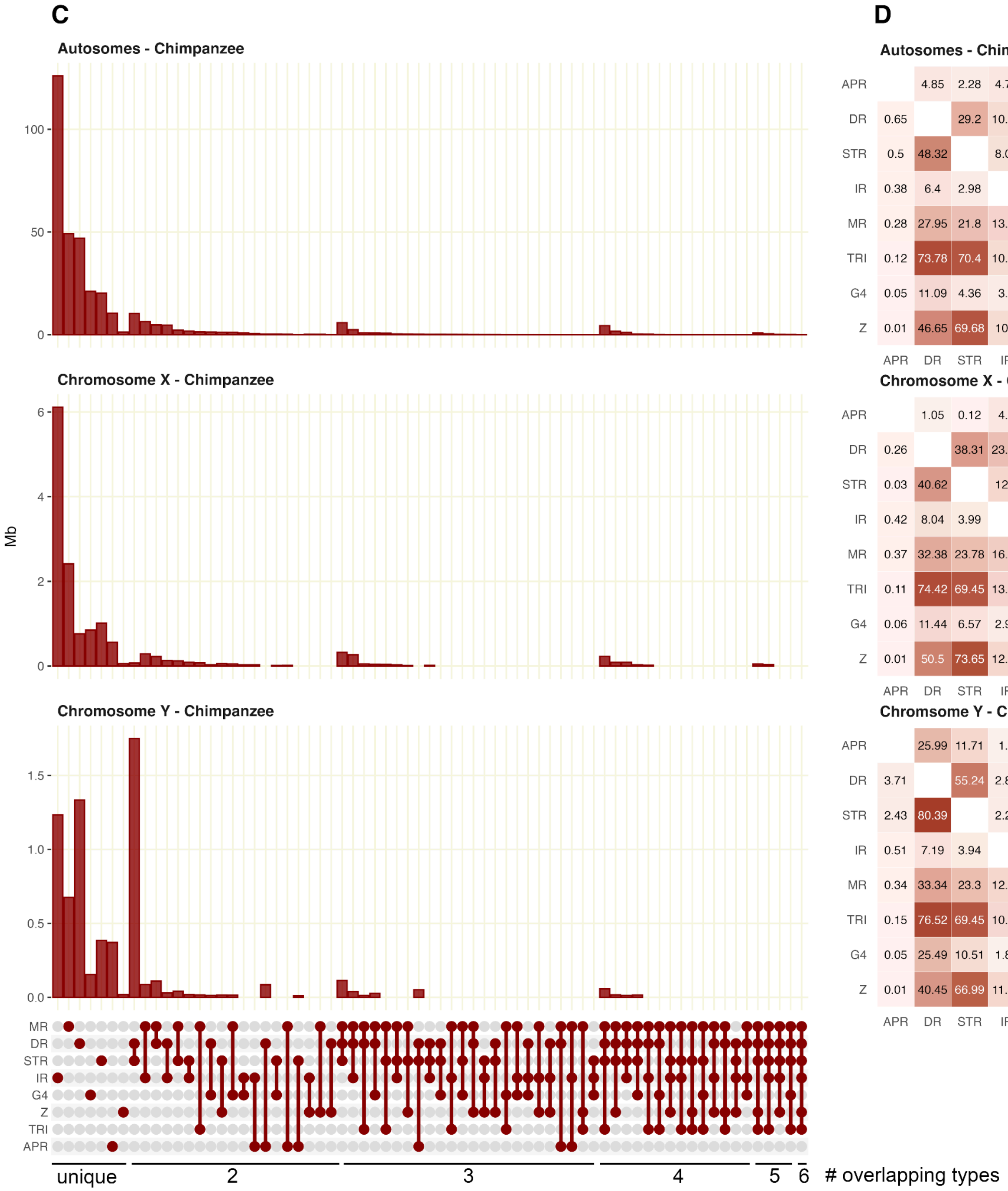


**D**

**Autosomes - Chimpanzee**

| | APR | DR | STR | IR | MR | TRI | G4 | Z |
|---|---|---|---|---|---|---|---|---|
| APR | | 4.85 | 2.28 | 4.71 | 2.04 | 0.09 | 0.11 | 0 |
| DR | 0.65 | | 29.2 | 10.51 | 27.44 | 7.98 | 3.33 | 3.46 |
| STR | 0.5 | 48.32 | | 8.09 | 35.41 | 12.6 | 2.17 | 8.54 |
| IR | 0.38 | 6.4 | 2.98 | | 7.96 | 0.66 | 0.66 | 0.46 |
| MR | 0.28 | 27.95 | 21.8 | 13.31 | | 11.01 | 3.25 | 4.06 |
| TRI | 0.12 | 73.78 | 70.4 | 10.02 | 100 | | 3.41 | 9.92 |
| G4 | 0.05 | 11.09 | 4.36 | 3.6 | 10.63 | 1.23 | | 0.07 |
| Z | 0.01 | 46.65 | 69.68 | 10.3 | 53.78 | 14.48 | 0.3 | |

**Chromosome X - Chimpanzee**

| | APR | DR | STR | IR | MR | TRI | G4 | Z |
|---|---|---|---|---|---|---|---|---|
| APR | | 1.05 | 0.12 | 4.9 | 2.76 | 0.1 | 0.11 | 0.01 |
| DR | 0.26 | | 38.31 | 23.02 | 58.89 | 15.82 | 4.91 | 6.52 |
| STR | 0.03 | 40.62 | | 12.1 | 45.86 | 15.65 | 2.99 | 10.08 |
| IR | 0.42 | 8.04 | 3.99 | | 10.65 | 0.98 | 0.45 | 0.55 |
| MR | 0.37 | 32.38 | 23.78 | 16.77 | | 11.69 | 3.22 | 3.99 |
| TRI | 0.11 | 74.42 | 69.45 | 13.16 | 100 | | 3.8 | 10.07 |
| G4 | 0.06 | 11.44 | 6.57 | 2.98 | 13.65 | 1.88 | | 0.04 |
| Z | 0.01 | 50.5 | 73.65 | 12.16 | 56.29 | 16.6 | 0.12 | |

**Chromsome Y - Chimpanzee**

| | APR | DR | STR | IR | MR | TRI | G4 | Z |
|---|---|---|---|---|---|---|---|---|
| APR | | 25.99 | 11.71 | 1.4 | 0.84 | 0.03 | 0.02 | 0 |
| DR | 3.71 | | 55.24 | 2.85 | 11.7 | 2.53 | 1.65 | 0.88 |
| STR | 2.43 | 80.39 | | 2.27 | 11.9 | 3.34 | 0.99 | 2.13 |
| IR | 0.51 | 7.19 | 3.94 | | 11.02 | 0.9 | 0.3 | 0.65 |
| MR | 0.34 | 33.34 | 23.3 | 12.43 | | 9.4 | 5.06 | 3.18 |
| TRI | 0.15 | 76.52 | 69.45 | 10.79 | 100 | | 4.25 | 8.32 |
| G4 | 0.05 | 25.49 | 10.51 | 1.84 | 27.37 | 2.16 | | 0.03 |
| Z | 0.01 | 40.45 | 66.99 | 11.83 | 51.03 | 12.56 | 0.08 | |

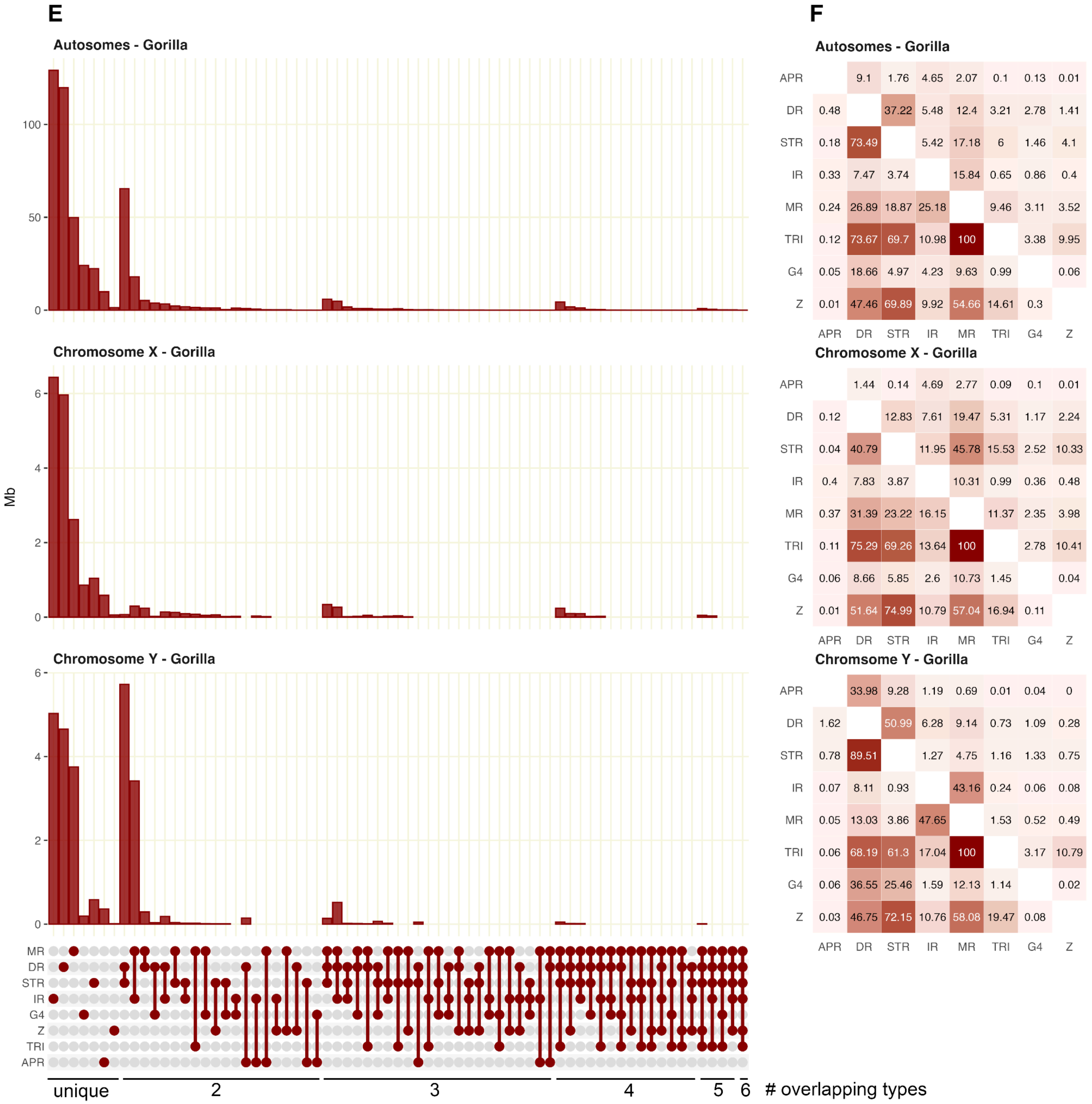
E
Autosomes - Gorilla
100
50
0
Chromosome X - Gorilla
6
4
2
0
Mb
Chromosome Y - Gorilla
6
4
2
0
MR
DR
STR
IR
G4
Z
TRI
APR
unique
2
3
4
5
6
### overlapping types
F
Autosomes - Gorilla
APR 9.1 1.76 4.65 2.07 0.1 0.13 0.01
DR 0.48 37.22 5.48 12.4 3.21 2.78 1.41
STR 0.18 73.49 5.42 17.18 6 1.46 4.1
IR 0.33 7.47 3.74 15.84 0.65 0.86 0.4
MR 0.24 26.89 18.87 25.18 9.46 3.11 3.52
TRI 0.12 73.67 69.7 10.98 100 3.38 9.95
G4 0.05 18.66 4.97 4.23 9.63 0.99 0.06
Z 0.01 47.46 69.89 9.92 54.66 14.61 0.3
APR DR STR IR MR TRI G4 Z
Chromosome X - Gorilla
APR 1.44 0.14 4.69 2.77 0.09 0.1 0.01
DR 0.12 12.83 7.61 19.47 5.31 1.17 2.24
STR 0.04 40.79 11.95 45.78 15.53 2.52 10.33
IR 0.4 7.83 3.87 10.31 0.99 0.36 0.48
MR 0.37 31.39 23.22 16.15 11.37 2.35 3.98
TRI 0.11 75.29 69.26 13.64 100 2.78 10.41
G4 0.06 8.66 5.85 2.6 10.73 1.45 0.04
Z 0.01 51.64 74.99 10.79 57.04 16.94 0.11
APR DR STR IR MR TRI G4 Z
Chromsome Y - Gorilla
APR 33.98 9.28 1.19 0.69 0.01 0.04 0
DR 1.62 50.99 6.28 9.14 0.73 1.09 0.28
STR 0.78 89.51 1.27 4.75 1.16 1.33 0.75
IR 0.07 8.11 0.93 43.16 0.24 0.06 0.08
MR 0.05 13.03 3.86 47.65 1.53 0.52 0.49
TRI 0.06 68.19 61.3 17.04 100 3.17 10.79
G4 0.06 36.55 25.46 1.59 12.13 1.14 0.02
Z 0.03 46.75 72.15 10.76 58.08 19.47 0.08
APR DR STR IR MR TRI G4 Z

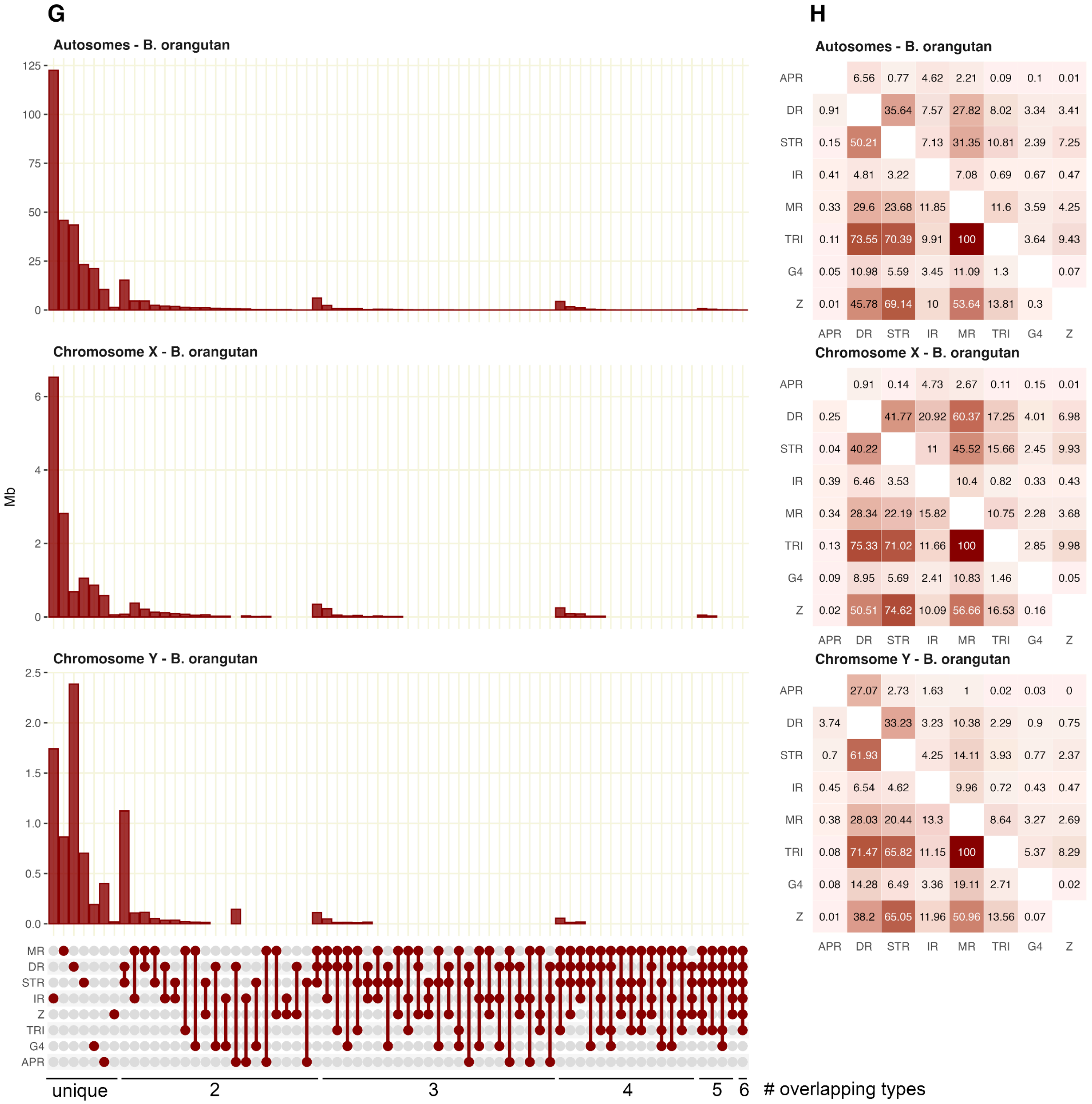
G
Autosomes - B. orangutan
Chromosome X - B. orangutan
Chromosome Y - B. orangutan
Mb
125 100 75 50 25 0
6 4 2 0
2.5 2.0 1.5 1.0 0.5 0.0
MR DR STR IR Z TRI G4 APR
unique 2 3 4 5 6 # overlapping types
H
Autosomes - B. orangutan
APR 6.56 0.77 4.62 2.21 0.09 0.1 0.01
DR 0.91 35.64 7.57 27.82 8.02 3.34 3.41
STR 0.15 50.21 7.13 31.35 10.81 2.39 7.25
IR 0.41 4.81 3.22 7.08 0.69 0.67 0.47
MR 0.33 29.6 23.68 11.85 11.6 3.59 4.25
TRI 0.11 73.55 70.39 9.91 100 3.64 9.43
G4 0.05 10.98 5.59 3.45 11.09 1.3 0.07
Z 0.01 45.78 69.14 10 53.64 13.81 0.3
APR DR STR IR MR TRI G4 Z
Chromosome X - B. orangutan
APR 0.91 0.14 4.73 2.67 0.11 0.15 0.01
DR 0.25 41.77 20.92 60.37 17.25 4.01 6.98
STR 0.04 40.22 11 45.52 15.66 2.45 9.93
IR 0.39 6.46 3.53 10.4 0.82 0.33 0.43
MR 0.34 28.34 22.19 15.82 10.75 2.28 3.68
TRI 0.13 75.33 71.02 11.66 100 2.85 9.98
G4 0.09 8.95 5.69 2.41 10.83 1.46 0.05
Z 0.02 50.51 74.62 10.09 56.66 16.53 0.16
APR DR STR IR MR TRI G4 Z
Chromsome Y - B. orangutan
APR 27.07 2.73 1.63 1 0.02 0.03 0
DR 3.74 33.23 3.23 10.38 2.29 0.9 0.75
STR 0.7 61.93 4.25 14.11 3.93 0.77 2.37
IR 0.45 6.54 4.62 9.96 0.72 0.43 0.47
MR 0.38 28.03 20.44 13.3 8.64 3.27 2.69
TRI 0.08 71.47 65.82 11.15 100 5.37 8.29
G4 0.08 14.28 6.49 3.36 19.11 2.71 0.02
Z 0.01 38.2 65.05 11.96 50.96 13.56 0.07
APR DR STR IR MR TRI G4 Z

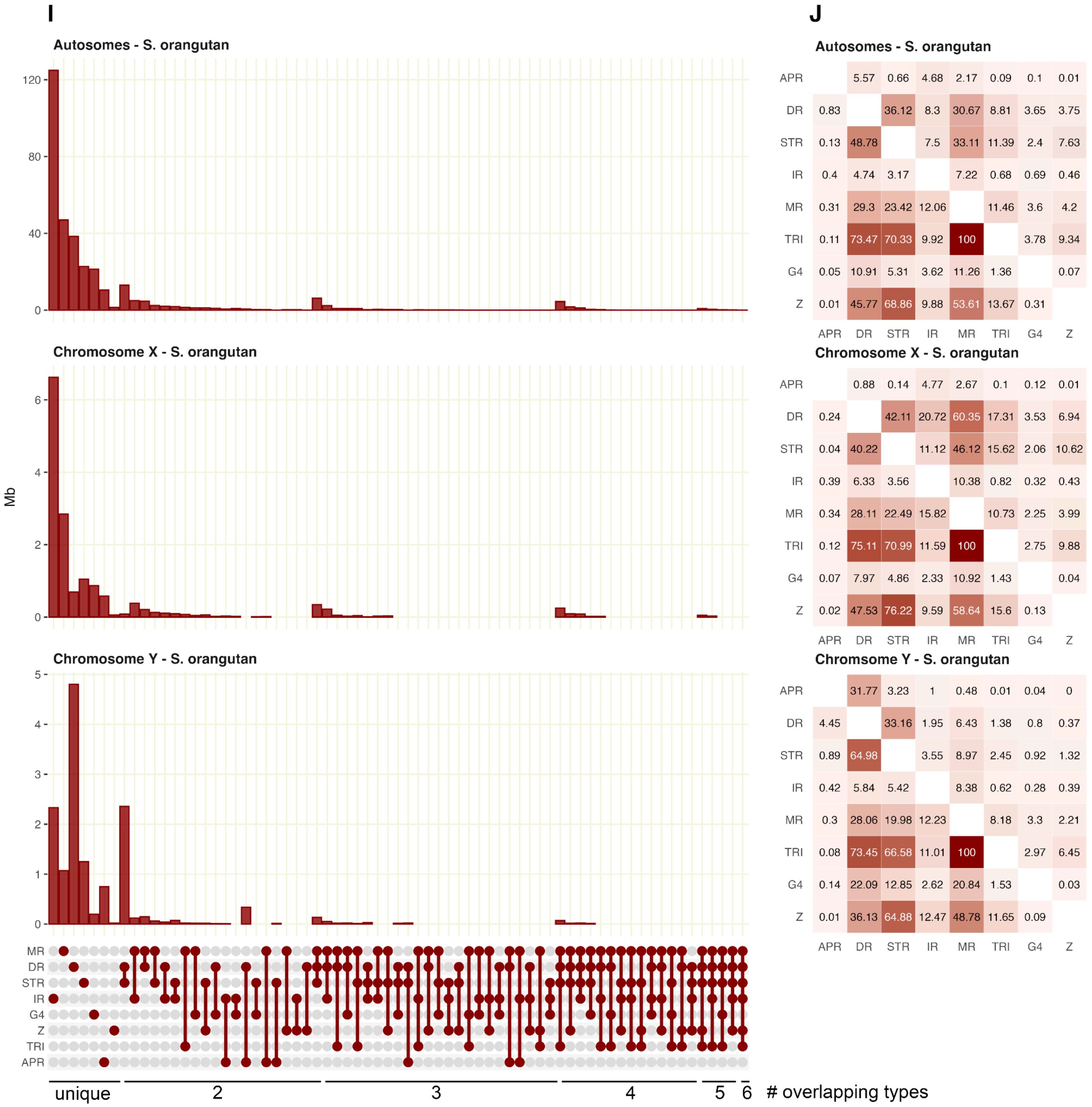
I
Autosomes - S. orangutan
120
80
40
0
Chromosome X - S. orangutan
Mb
6
4
2
0
Chromosome Y - S. orangutan
5
4
3
2
1
0
MR
DR
STR
IR
G4
Z
TRI
APR
unique
2
3
4
5
6
### overlapping types
J
Autosomes - S. orangutan
APR 5.57 0.66 4.68 2.17 0.09 0.1 0.01
DR 0.83 36.12 8.3 30.67 8.81 3.65 3.75
STR 0.13 48.78 7.5 33.11 11.39 2.4 7.63
IR 0.4 4.74 3.17 7.22 0.68 0.69 0.46
MR 0.31 29.3 23.42 12.06 11.46 3.6 4.2
TRI 0.11 73.47 70.33 9.92 100 3.78 9.34
G4 0.05 10.91 5.31 3.62 11.26 1.36 0.07
Z 0.01 45.77 68.86 9.88 53.61 13.67 0.31
APR DR STR IR MR TRI G4 Z
Chromosome X - S. orangutan
APR 0.88 0.14 4.77 2.67 0.1 0.12 0.01
DR 0.24 42.11 20.72 60.35 17.31 3.53 6.94
STR 0.04 40.22 11.12 46.12 15.62 2.06 10.62
IR 0.39 6.33 3.56 10.38 0.82 0.32 0.43
MR 0.34 28.11 22.49 15.82 10.73 2.25 3.99
TRI 0.12 75.11 70.99 11.59 100 2.75 9.88
G4 0.07 7.97 4.86 2.33 10.92 1.43 0.04
Z 0.02 47.53 76.22 9.59 58.64 15.6 0.13
APR DR STR IR MR TRI G4 Z
Chromsome Y - S. orangutan
APR 31.77 3.23 1 0.48 0.01 0.04 0
DR 4.45 33.16 1.95 6.43 1.38 0.8 0.37
STR 0.89 64.98 3.55 8.97 2.45 0.92 1.32
IR 0.42 5.84 5.42 8.38 0.62 0.28 0.39
MR 0.3 28.06 19.98 12.23 8.18 3.3 2.21
TRI 0.08 73.45 66.58 11.01 100 2.97 6.45
G4 0.14 22.09 12.85 2.62 20.84 1.53 0.03
Z 0.01 36.13 64.88 12.47 48.78 11.65 0.09
APR DR STR IR MR TRI G4 Z

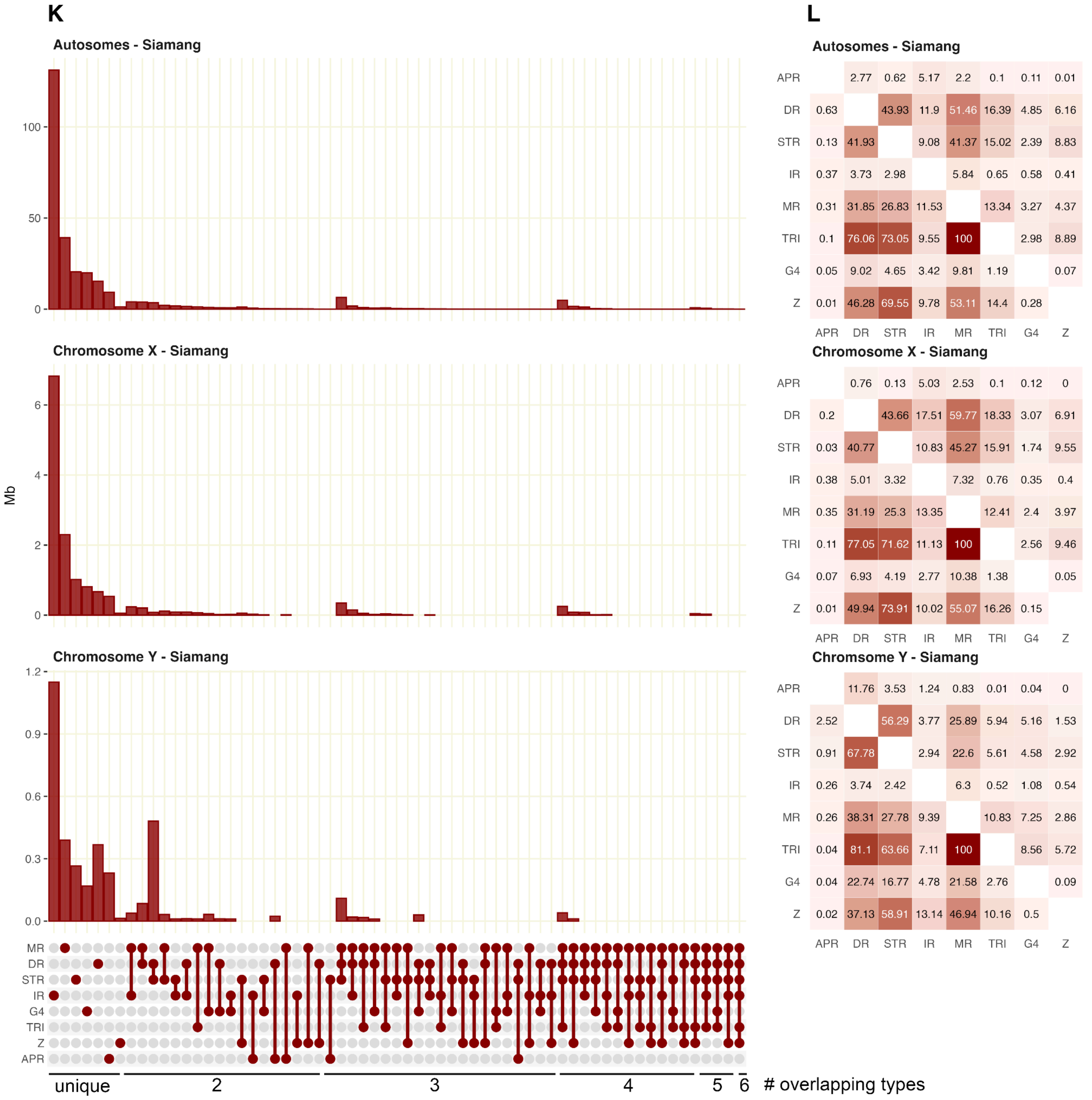
K
Autosomes - Siamang
100
50
0
Chromosome X - Siamang
6
4
2
0
Mb
Chromosome Y - Siamang
1.2
0.9
0.6
0.3
0.0
MR
DR
STR
IR
G4
TRI
Z
APR
unique
2
3
4
5
6
### overlapping types
L
Autosomes - Siamang
APR 2.77 0.62 5.17 2.2 0.1 0.11 0.01
DR 0.63 43.93 11.9 51.46 16.39 4.85 6.16
STR 0.13 41.93 9.08 41.37 15.02 2.39 8.83
IR 0.37 3.73 2.98 5.84 0.65 0.58 0.41
MR 0.31 31.85 26.83 11.53 13.34 3.27 4.37
TRI 0.1 76.06 73.05 9.55 100 2.98 8.89
G4 0.05 9.02 4.65 3.42 9.81 1.19 0.07
Z 0.01 46.28 69.55 9.78 53.11 14.4 0.28
APR DR STR IR MR TRI G4 Z
Chromosome X - Siamang
APR 0.76 0.13 5.03 2.53 0.1 0.12 0
DR 0.2 43.66 17.51 59.77 18.33 3.07 6.91
STR 0.03 40.77 10.83 45.27 15.91 1.74 9.55
IR 0.38 5.01 3.32 7.32 0.76 0.35 0.4
MR 0.35 31.19 25.3 13.35 12.41 2.4 3.97
TRI 0.11 77.05 71.62 11.13 100 2.56 9.46
G4 0.07 6.93 4.19 2.77 10.38 1.38 0.05
Z 0.01 49.94 73.91 10.02 55.07 16.26 0.15
APR DR STR IR MR TRI G4 Z
Chromsome Y - Siamang
APR 11.76 3.53 1.24 0.83 0.01 0.04 0
DR 2.52 56.29 3.77 25.89 5.94 5.16 1.53
STR 0.91 67.78 2.94 22.6 5.61 4.58 2.92
IR 0.26 3.74 2.42 6.3 0.52 1.08 0.54
MR 0.26 38.31 27.78 9.39 10.83 7.25 2.86
TRI 0.04 81.1 63.66 7.11 100 8.56 5.72
G4 0.04 22.74 16.77 4.78 21.58 2.76 0.09
Z 0.02 37.13 58.91 13.14 46.94 10.16 0.5
APR DR STR IR MR TRI G4 Z

**Figure S5. Circos plots of non-B DNA motif density for siamang haplotypes**
Circos plots of non-B DNA motif density along chromosomes for **(A)** primary haplotype and **(B)** alternative haplotype, of siamang. The density of each motif type is scaled separately by the window with the highest density. Abbreviations: APR: A-phased repeats; DR: direct repeats; STR: short tandem repeats; IR: inverted repeats; MR: mirror repeats; TRI: triplex motifs; G4: G-quadruplexes; Z: Z-DNA. Animal silhouette is from [https://www.phylopic.org](https://www.phylopic.org).

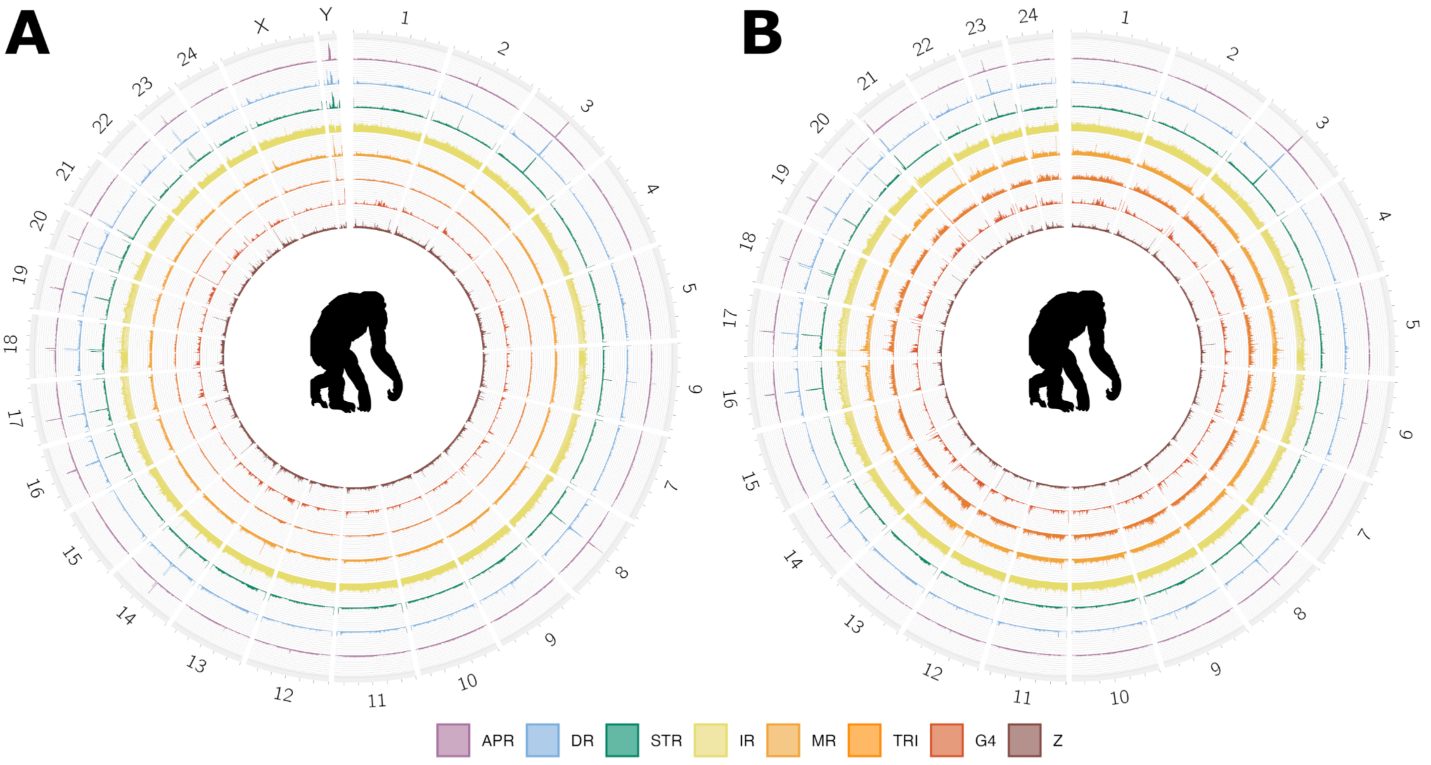

**Figure S6. Circos plots of  non-B DNA motif density for alternative haplotypes**
Circos plots of non-B DNA motif density along chromosomes for alternative haplotypes of **(A)** bonobo, **(B)** chimpanzee, **(C)** gorilla, **(D)** Bornean orangutan, and **(E)** Sumatran orangutan. Human CHM13 has no alternative haplotype because this cell line is homozygous. Abbreviations are as in Figure S5. Animal silhouettes are from https://www.phylopic.org.

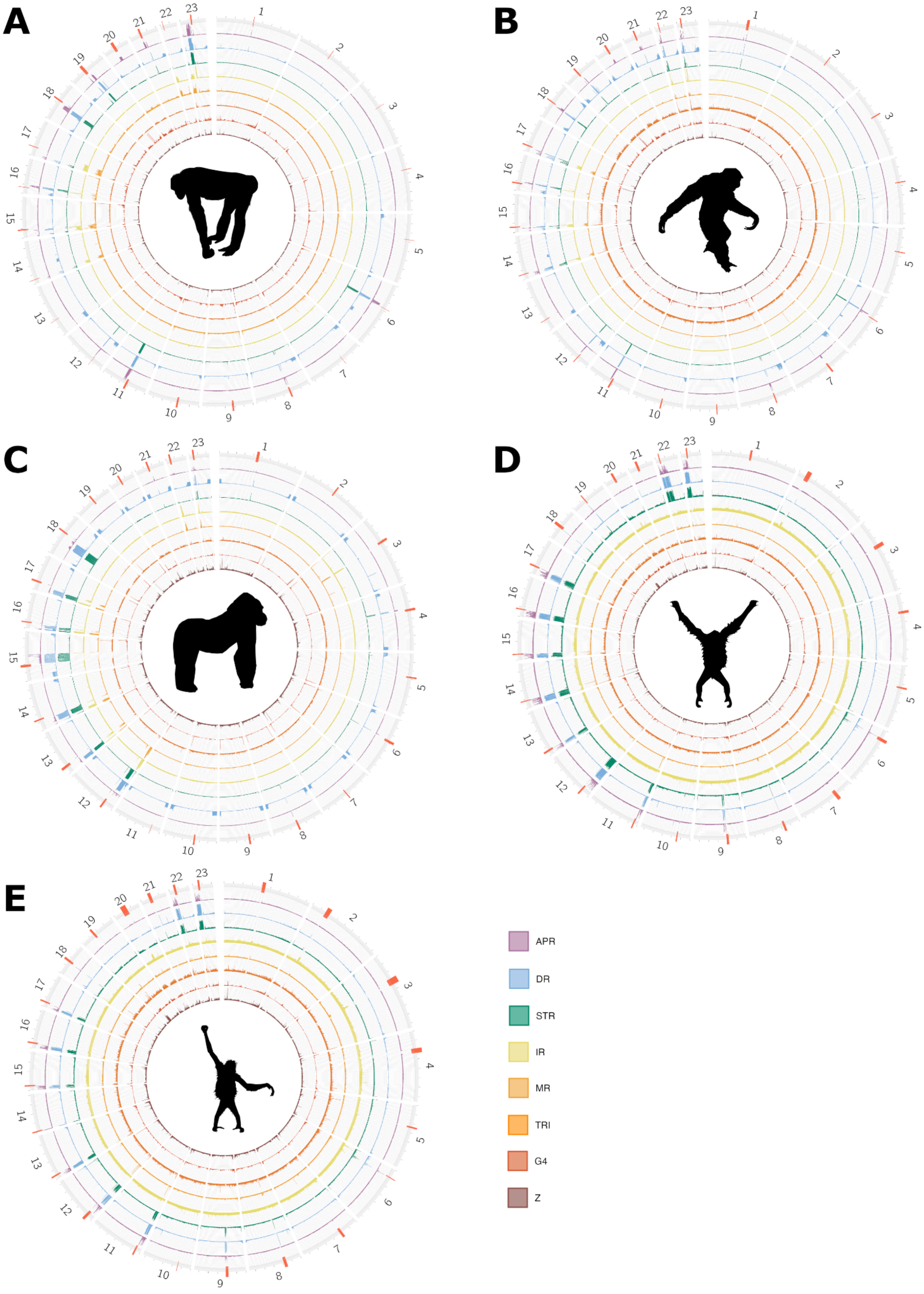

**Figure S7. Density plot of non-B DNA motif density for acrocentric short arms**
Detailed density plot for short arms of the acrocentric chromosomes in **(A)** bonobo, **(B)** chimpanzee, **(C)** human, **(D)** gorilla, **(E)** Bornean orangutan, and **(F)** Sumatran orangutan, based on annotations from Yoo et al. (2024). Normalized non-B density is shown for 100-kb windows, where darker colors represent higher non-B content. Centromeric satellite repeats are shown in a separate track using the color scheme from the UCSC Genome Browser. Black arrows represent rDNA.

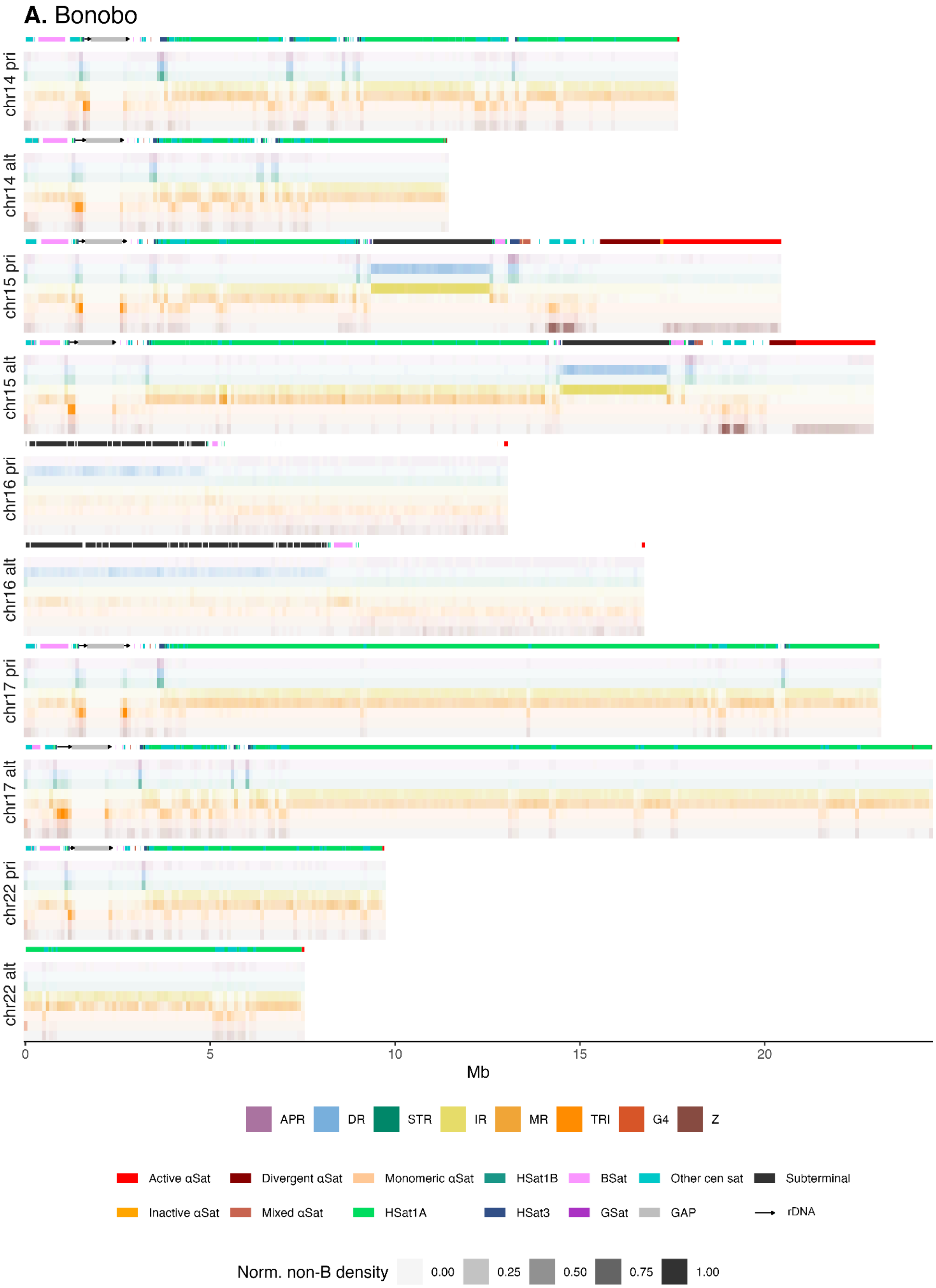

**B.** Chimpanzee

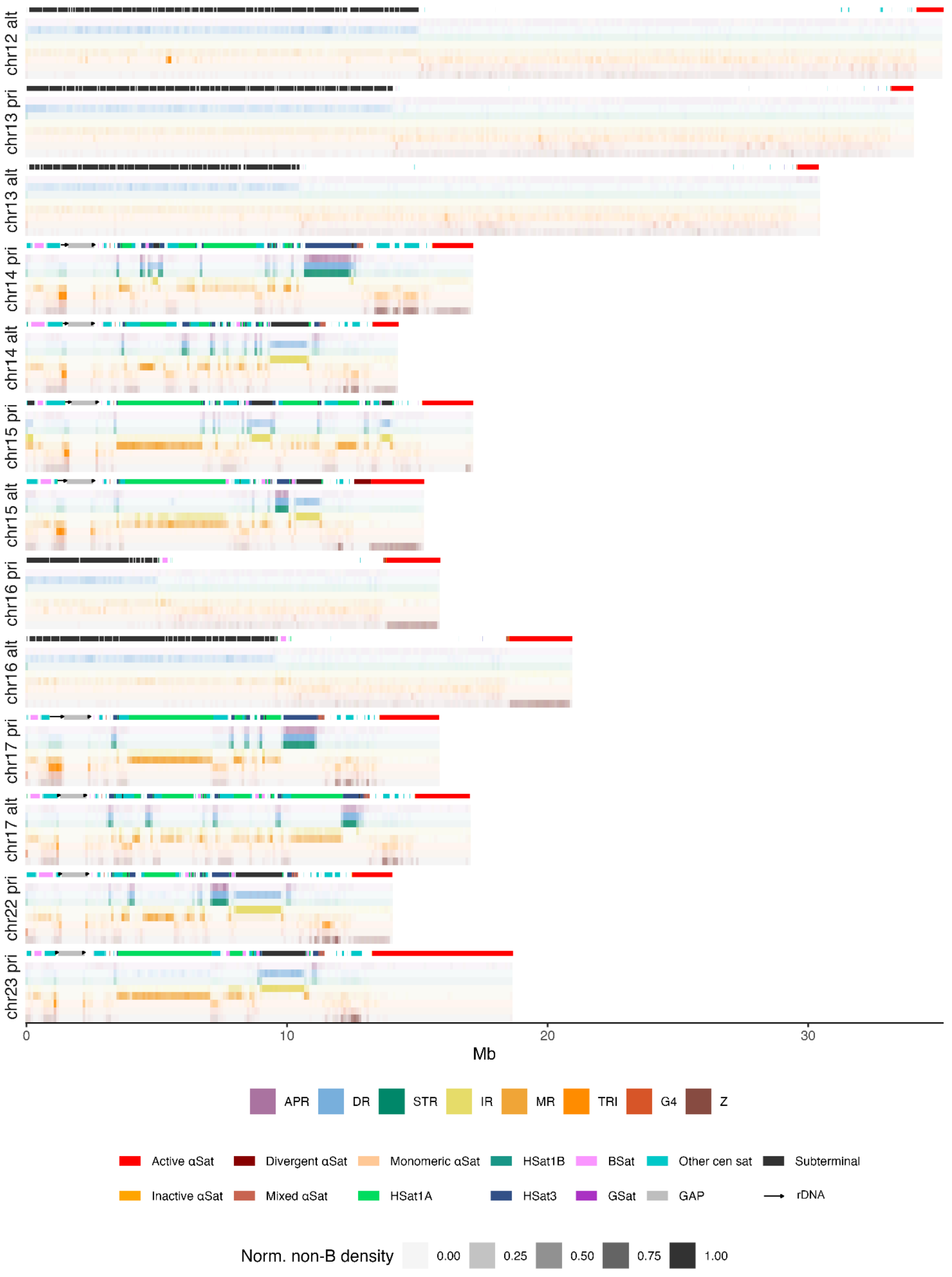

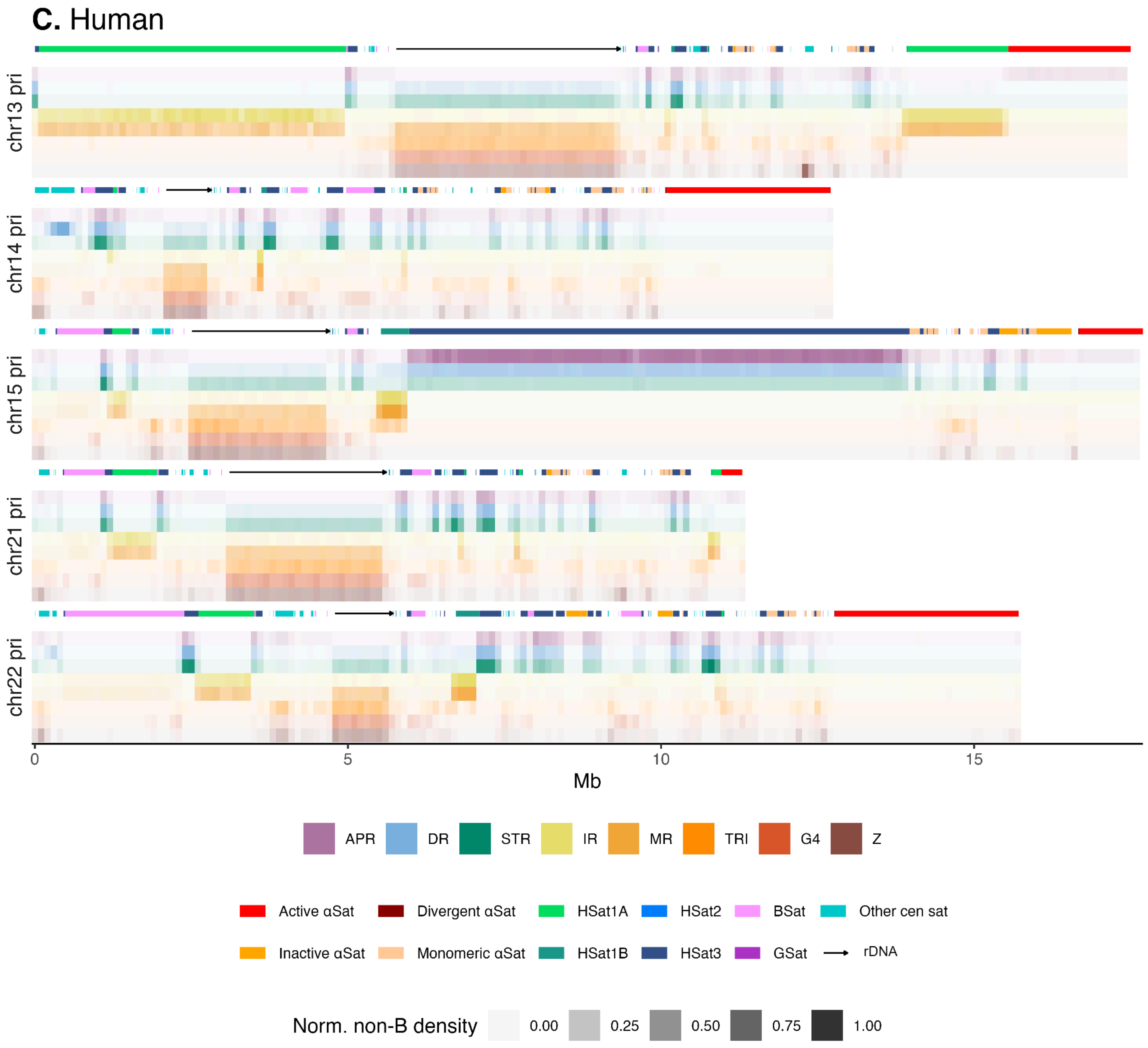
C. Human
chr13 pri
chr14 pri
chr15 pri
chr21 pri
chr22 pri
0
5
10
15
Mb
APR
DR
STR
IR
MR
TRI
G4
Z
Active αSat
Divergent αSat
HSat1A
HSat2
BSat
Other cen sat
Inactive αSat
Monomeric αSat
HSat1B
HSat3
GSat
rDNA
Norm. non-B density
0.00
0.25
0.50
0.75
1.00

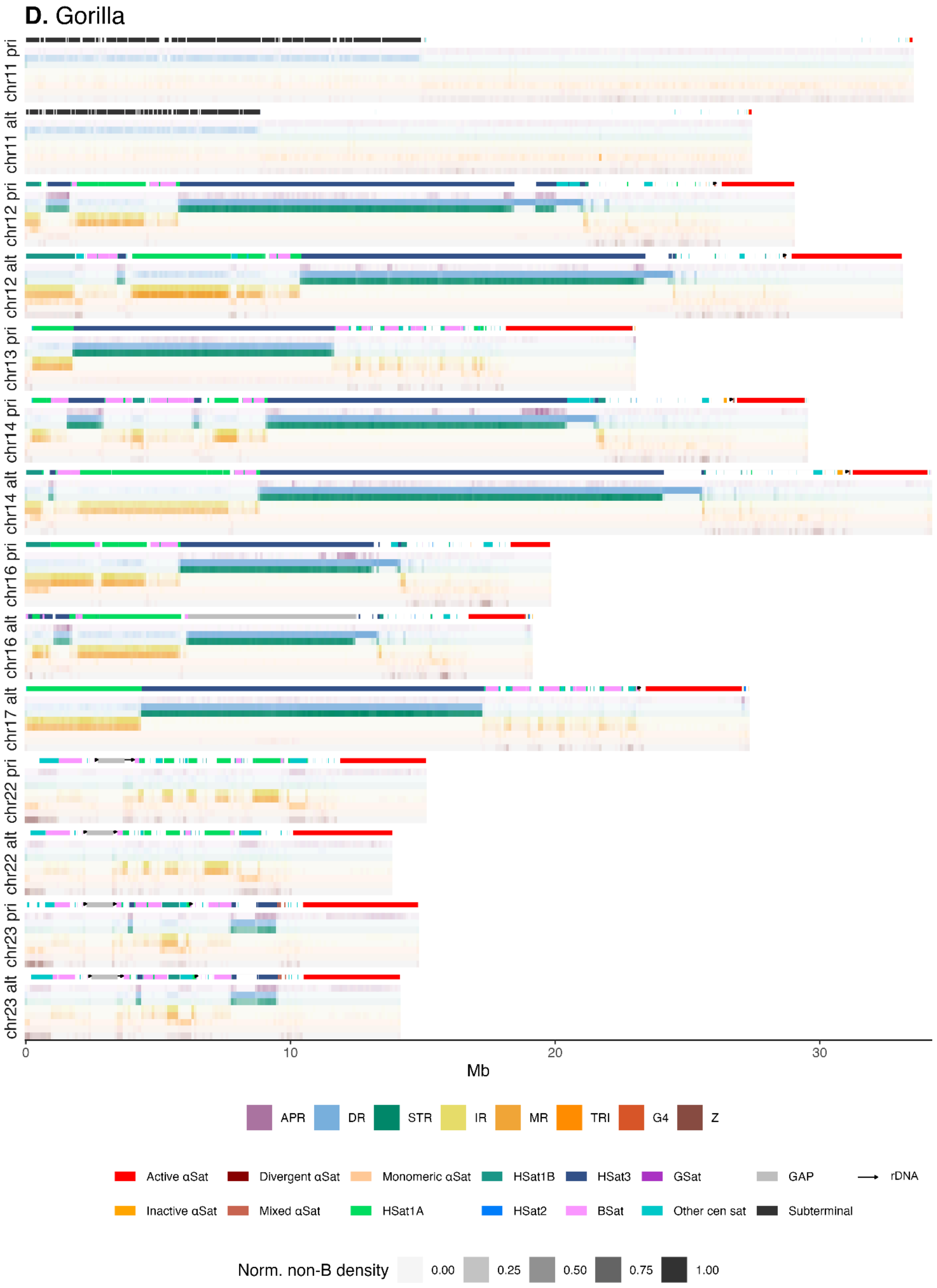

D. Gorilla
chr11 pri
chr11 alt
chr12 pri
chr12 alt
chr13 pri
chr14 pri
chr14 alt
chr16 pri
chr16 alt
chr17 alt
chr22 pri
chr22 alt
chr23 pri
chr23 alt
0
10
20
30
Mb
APR
DR
STR
IR
MR
TRI
G4
Z
Active αSat
Divergent αSat
Monomeric αSat
HSat1B
HSat3
GSat
GAP
rDNA
Inactive αSat
Mixed αSat
HSat1A
HSat2
BSat
Other cen sat
Subterminal
Norm. non-B density
0.00
0.25
0.50
0.75
1.00

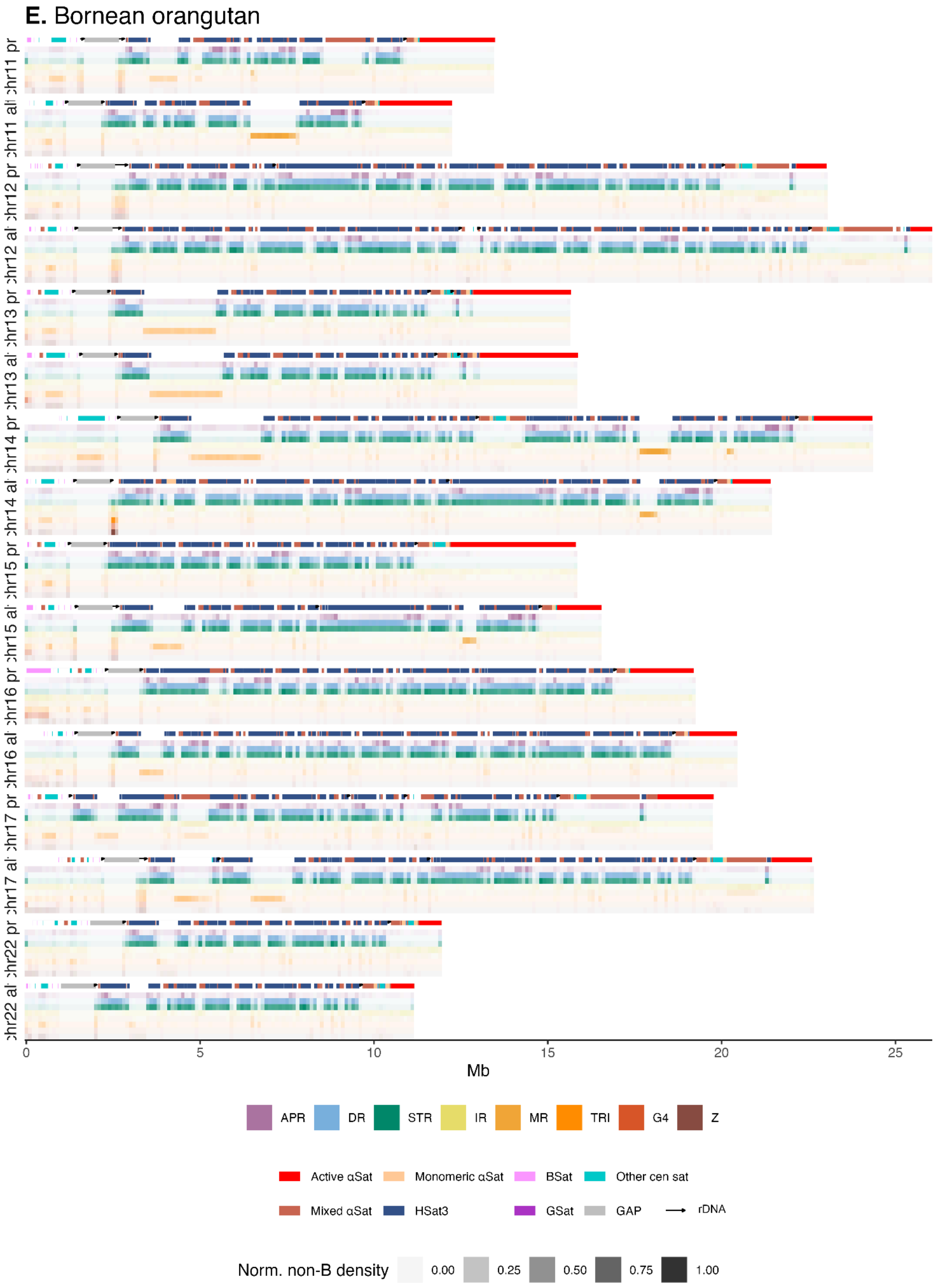

E. Bornean orangutan
chr11 pr
chr11 al
chr12 pr
chr12 al
chr13 pr
chr13 al
chr14 pr
chr14 al
chr15 pr
chr15 al
chr16 pr
chr16 al
chr17 pr
chr17 al
chr22 pr
chr22 al
0
5
10
15
20
25
Mb
APR
DR
STR
IR
MR
TRI
G4
Z
Active αSat
Monomeric αSat
BSat
Other cen sat
Mixed αSat
HSat3
GSat
GAP
rDNA
Norm. non-B density
0.00
0.25
0.50
0.75
1.00

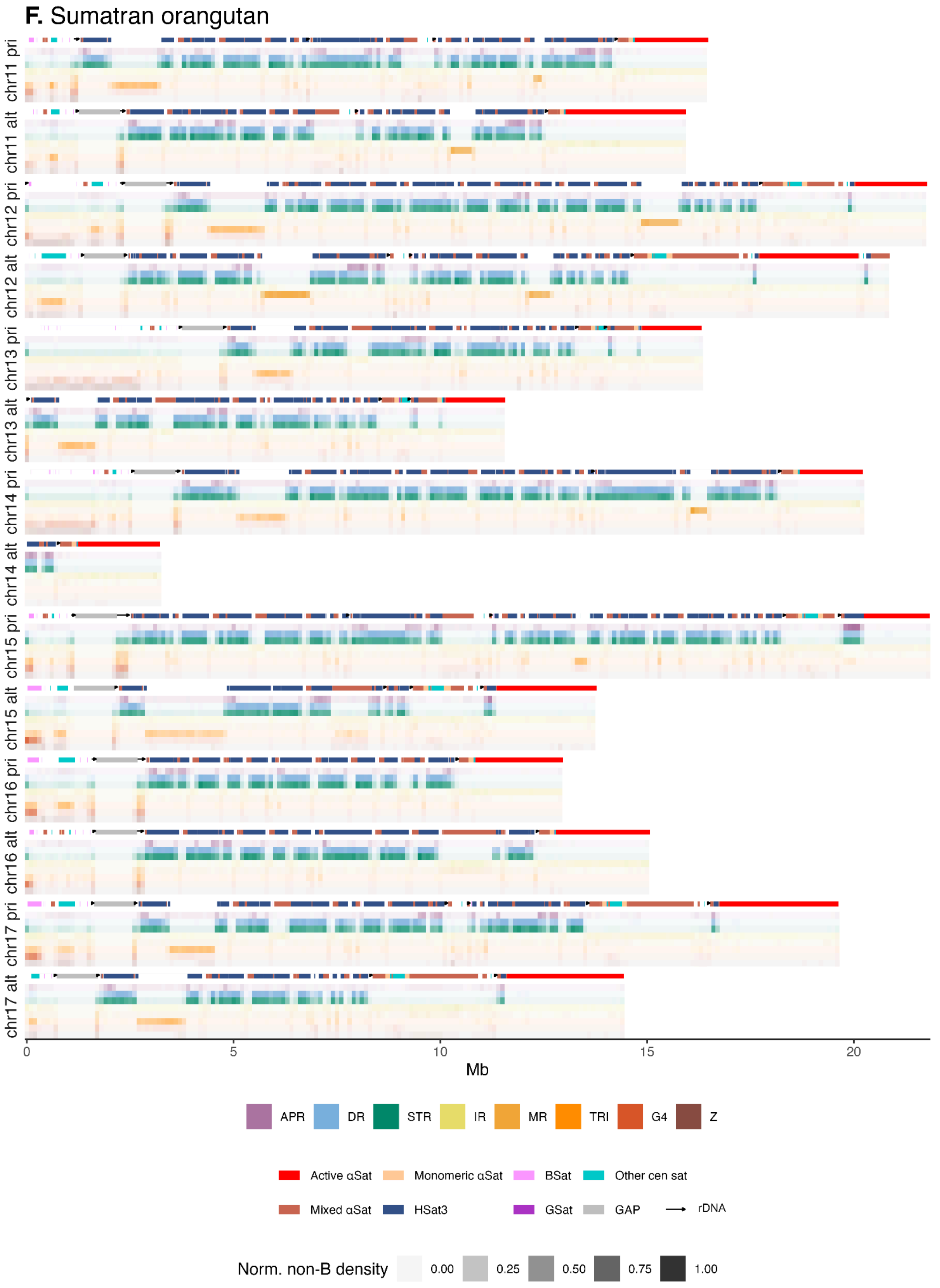

F. Sumatran orangutan
chr11 pri
chr11 alt
chr12 pri
chr12 alt
chr13 pri
chr13 alt
chr14 pri
chr14 alt
chr15 pri
chr15 alt
chr16 pri
chr16 alt
chr17 pri
chr17 alt
0
5
10
15
20
Mb
APR
DR
STR
IR
MR
TRI
G4
Z
Active αSat
Monomeric αSat
BSat
Other cen sat
Mixed αSat
HSat3
GSat
GAP
rDNA
Norm. non-B density
0.00
0.25
0.50
0.75
1.00

**Figure S8. Non-B DNA density plot of Y chromosomes**
Detailed density plot for the Y chromosome in bonobo, chimpanzee, human, gorilla, Bornean orangutan, and Sumatran orangutan, based on annotations from Makova et al. (2024). Normalized non-B density is shown for 100-kb windows, where darker colors represent higher non-B content. Centromeric satellite repeats are shown in a separate track using the color scheme from the UCSC Genome Browser. Black arrows represent rDNA.

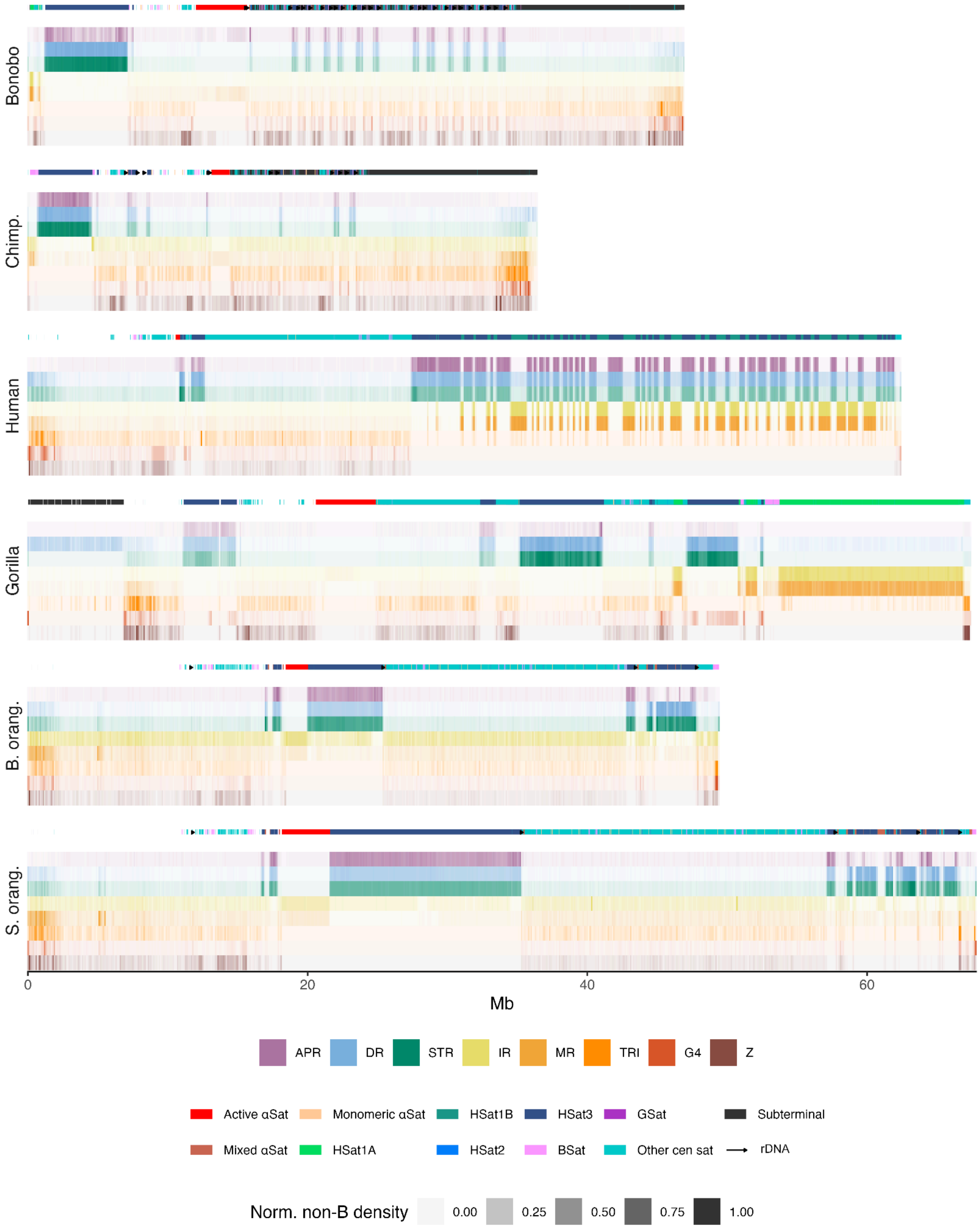

**Figure S9. Enrichment of non-B DNA at centromeric and acrocentric satellites**

Fold enrichment of non-B DNA at the centromeric and acrocentric satellites from Fig. S7 for **(A)** bonobo, **(B)** chimpanzee, (**C)** human, **(D)** gorilla, **(E)** Bornean orangutan, **(F)** Sumatran orangutan, and **(G)** siamang. Underrepresentation of non-B DNA (values below 1) is shown in blue, while enrichment (values above 1) is shown in red. Fold enrichment is calculated as motif density in the region divided by the genome-wide density. Active αSat represents the active centromere. If non-B DNA motifs of a certain type are missing for a repeat, the tile is colored in dark blue. Note that not all satellite classes are present in orangutans and siamang.

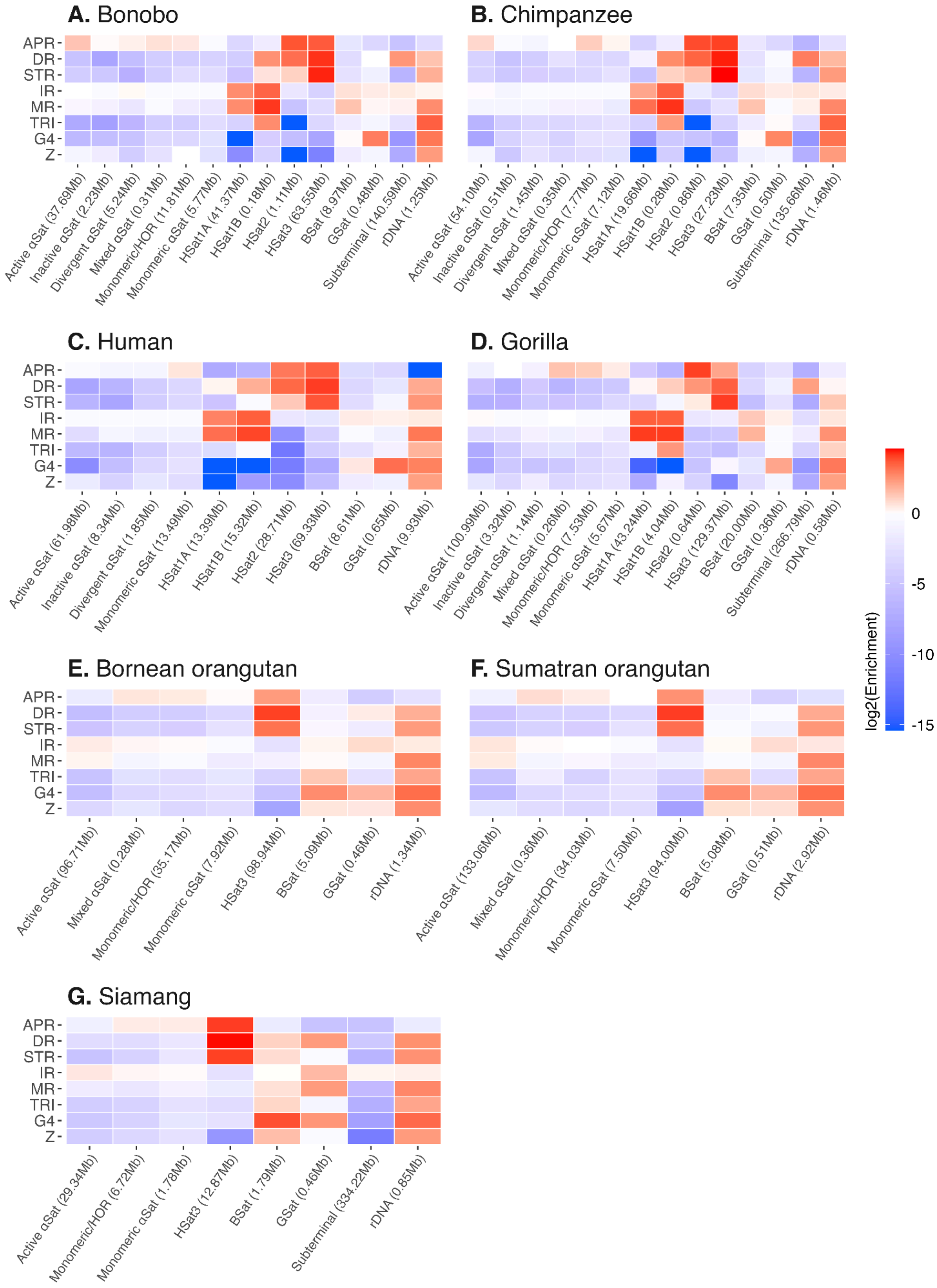

**Figure S10. Enrichment of non-B DNA at functional regions**
Fold enrichment of non-B DNA at functional regions (same as Fig. 5) with confidence intervals based on **(A)** resampling 10% of the data 20 times, or **(B)** resampling 50% of the data 20 times. The red dashed line (fold enrichment=1) represents the genome-wide average. Bars with confidence intervals overlapping the red line are not considered to be significantly different from the genome-wide average.

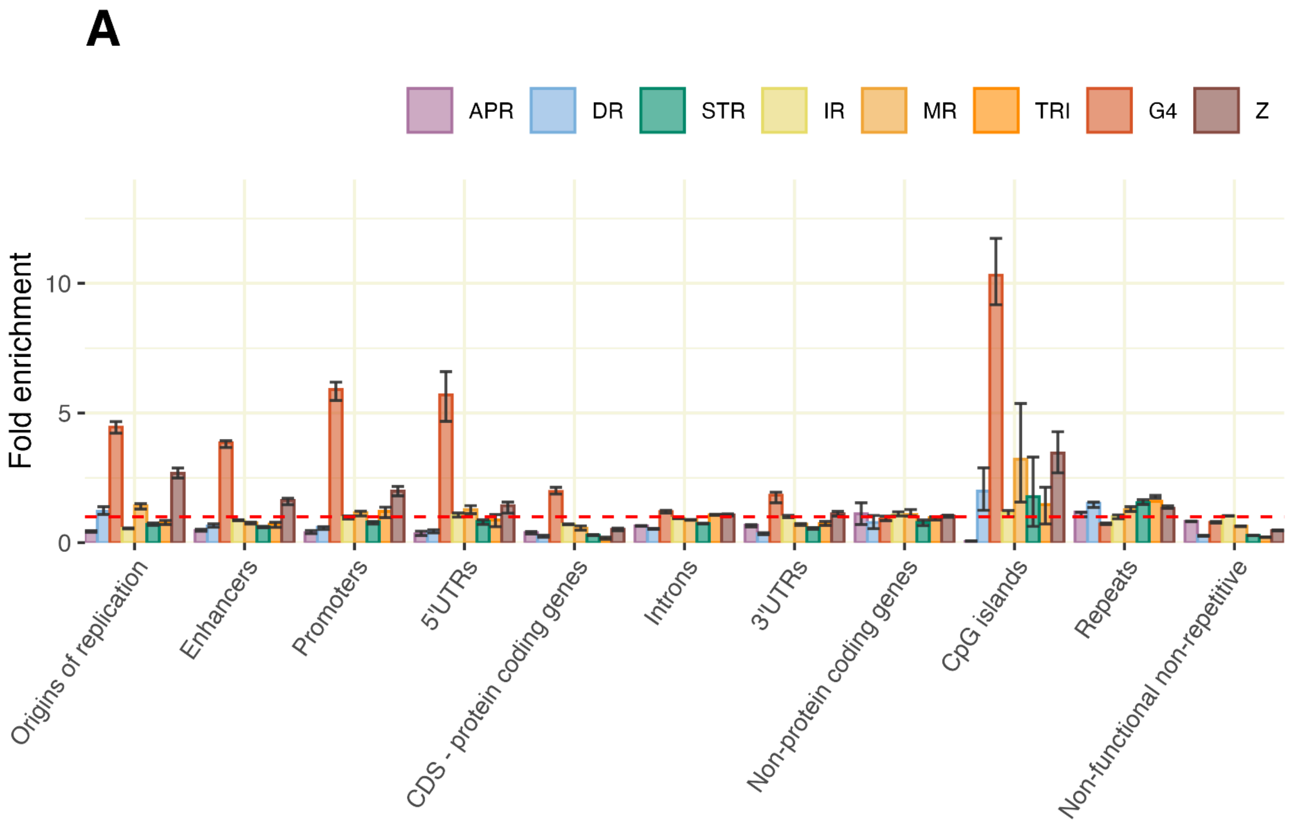


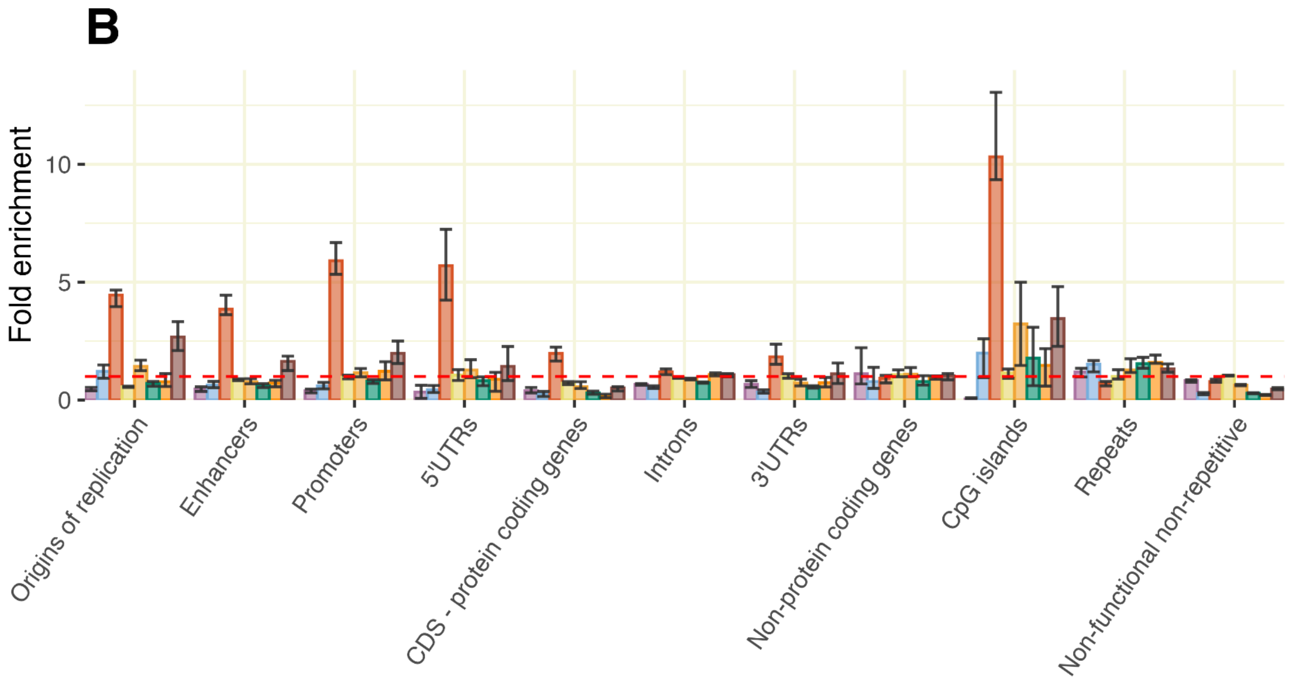

**Figure S11. GC-corrected enrichment of non-B DNA at functional regions**
Fold enrichment of non-B DNA at functional regions (same as Fig. 5) with the G4s corrected for GC content. Corrected G4 bars (using a simple conversion factor, see Methods) are outlined in black, while the original fold enrichment is seen behind in lighter vermilion. The red dashed line (fold enrichment=1) represents the genome-wide average.

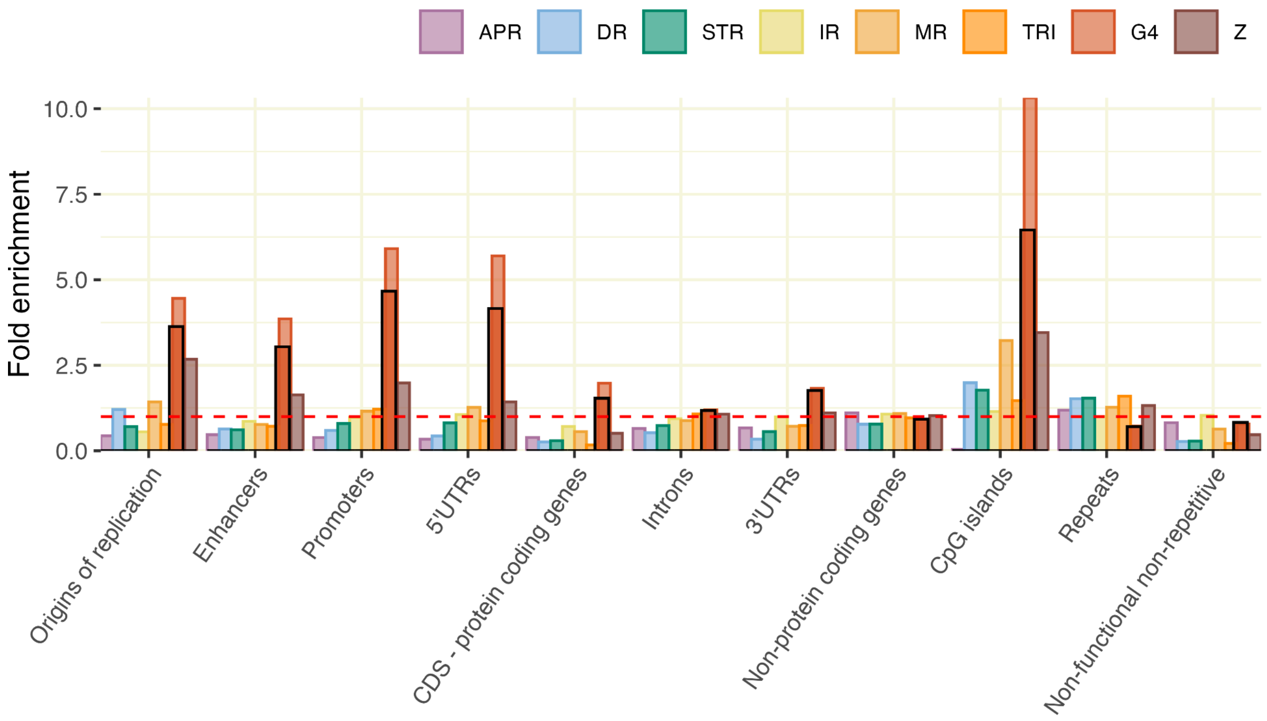

**Figure S12. Enrichment of non-B DNA at manually curated human repeats**
Enrichment of non-B DNA at manually curated repeat annotations for human, given as log-fold densities compared to genome-wide densities. (**A**) Enrichment at new satellites and (**B**) enrichment at composite repeats. If non-B DNA motifs of a certain type are missing for a repeat, the tile is colored in dark blue. Total repeat lengths are given after the names. Repeats with a total length shorter than 5 kb are not shown.

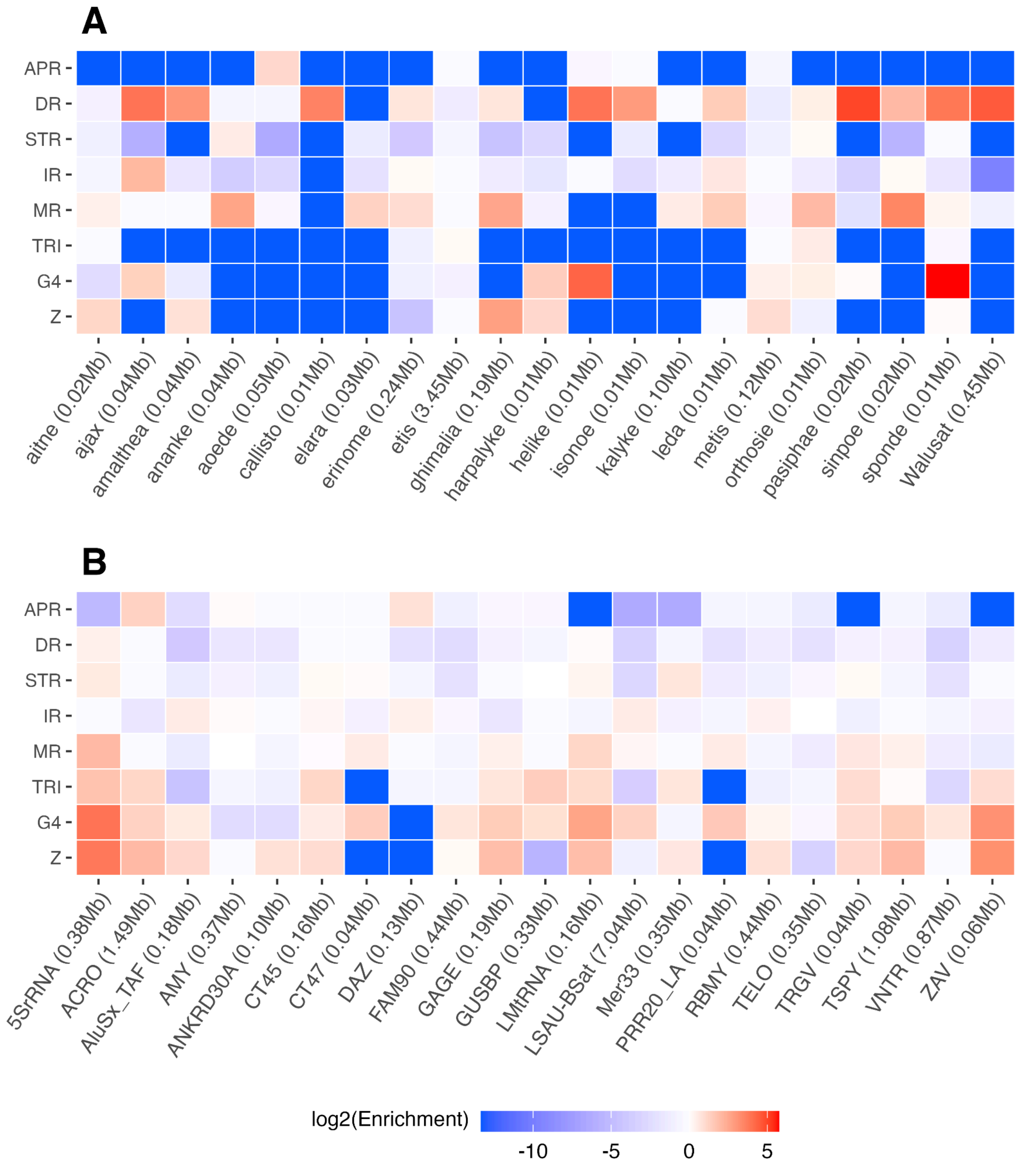

**Figure S13. Enrichment of non-B DNA at repeats in non-human apes**

Enrichment of non-B DNA at repeats, given as log-fold densities compared to genome-wide densities for (**A**) bonobo, **(B)** chimpanzee, **(C)** gorilla, **(D)** Bornean orangutan, **(E)** Sumatran orangutan, and **(F)** siamang. Underrepresentation of non-B DNA (values below 1) is shown in blue, while enrichment (values above 1) is shown in red. If non-B DNA motifs of a certain type are missing for a repeat, the tile is colored in dark blue. Long repeat names have been shortened for visualization purposes (marked with *). Total repeat lengths are given after the names. Repeats with a total length shorter than 50 kb are not shown.

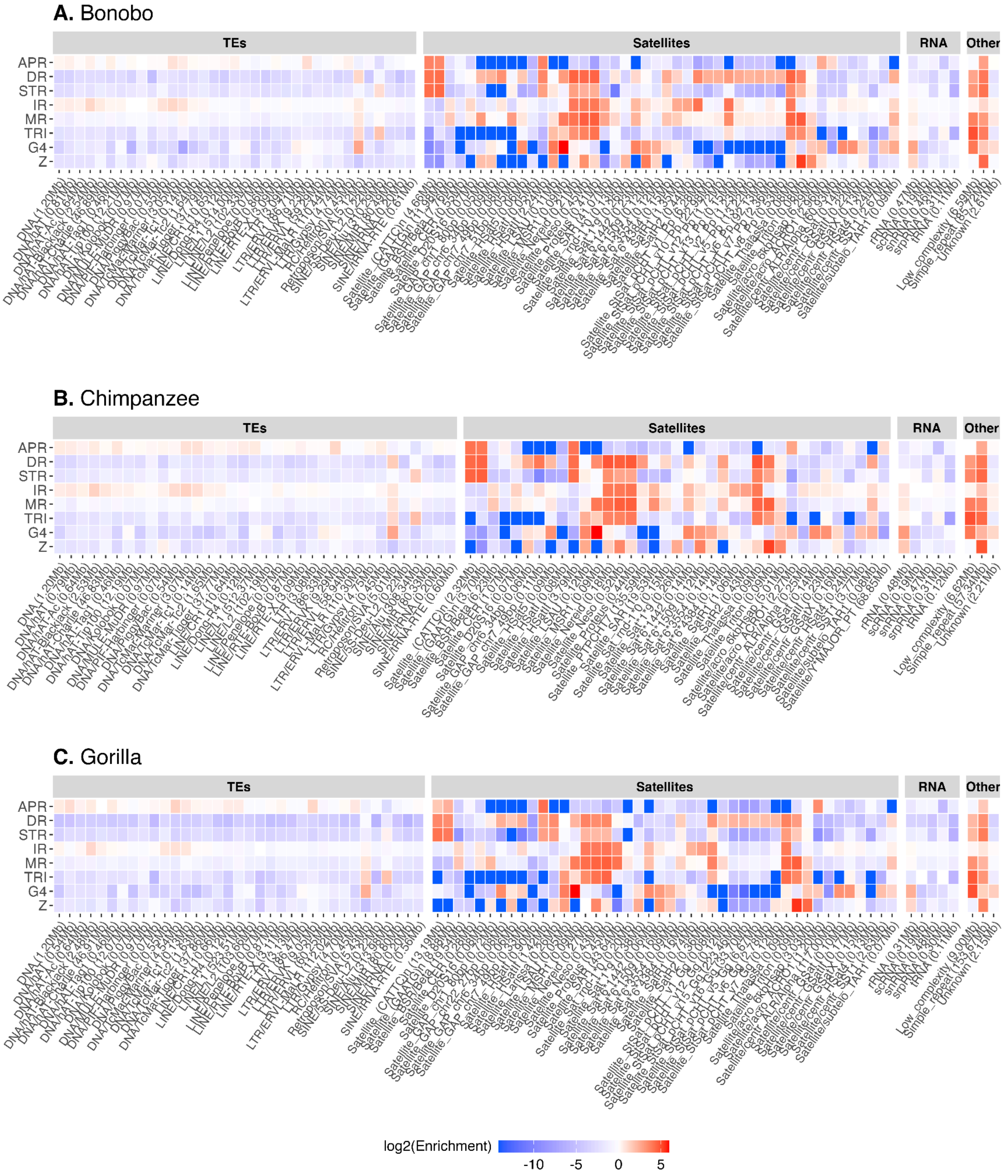

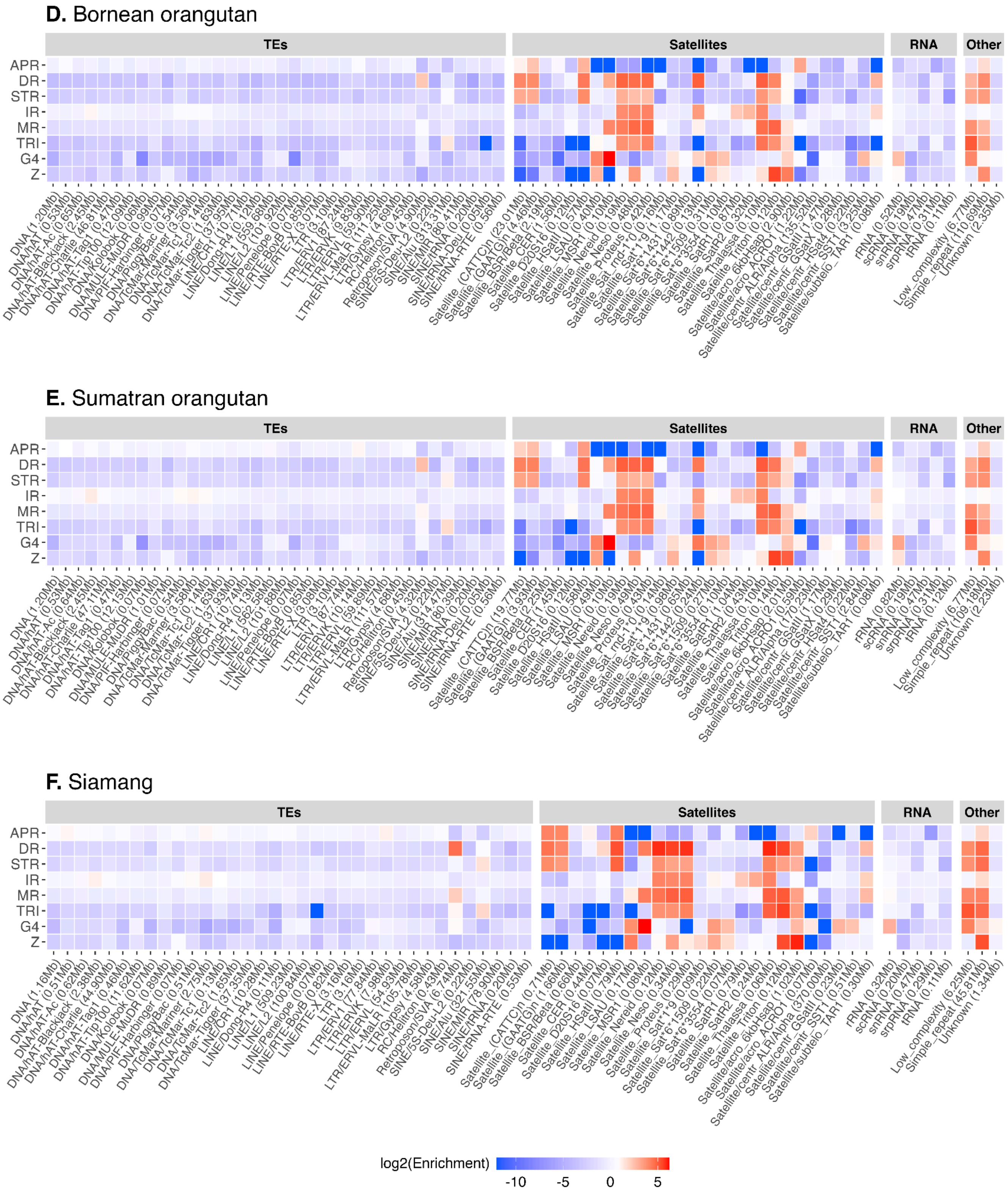
D. Bornean orangutan
E. Sumatran orangutan
F. Siamang
TEs
Satellites
RNA
Other
APR
DR
STR
IR
MR
TRI
G4
Z
log2(Enrichment)
-10
-5
0
5

**Figure S14. Experimental validation of G4s**

Experimental validation of three G4s in the satellite LSAU (LS1-LS3) and one G4 in Walusat (LS4). LS1 and LS3 formed parallel G4s (see Methods for actual sequences). (**A**) Isothermal difference spectra (IDS), (**B**) Thermal difference spectra (TDS). G4 samples are characterized by a minimum of around 297 nm (LS1 and LS3). (**C**) Polyacrylamide gel electrophoresis (PAGE) of the four motifs. Samples were prepared either immediately (loaded on the gel after adding K+) or 24 hours before loading onto the gel. Samples LS1-LS3 migrated as intramolecular species, while LS4 was bimolecular.

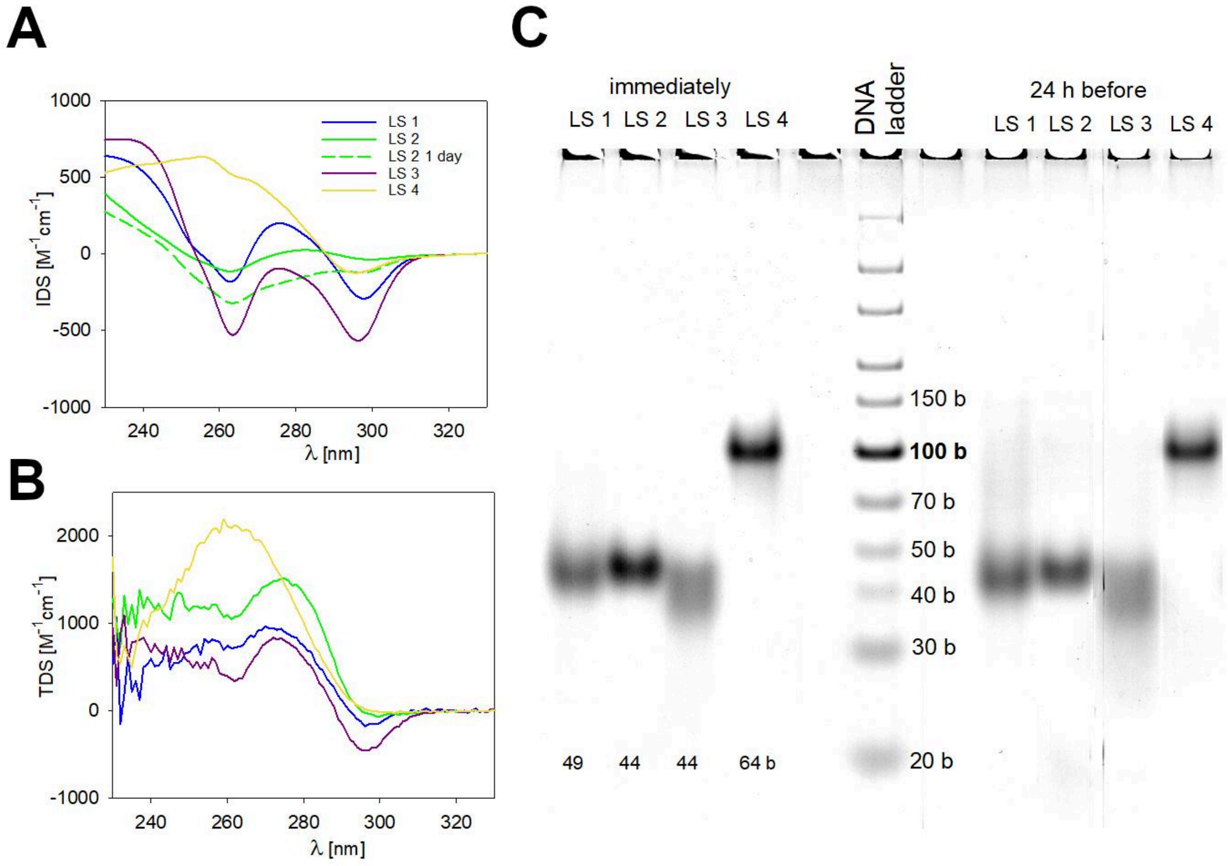

**Figure S15. Non-B DNA motifs at SST1**
Example of non-B DNA motifs in and between monomers of the satellite SST1, subtype sf1, in human. Six monomers (out of ~100 on chromosome 13) are shown as boxes below the annotated non-B motifs. Z-DNA motifs occur in all spacers between sf1 monomers.

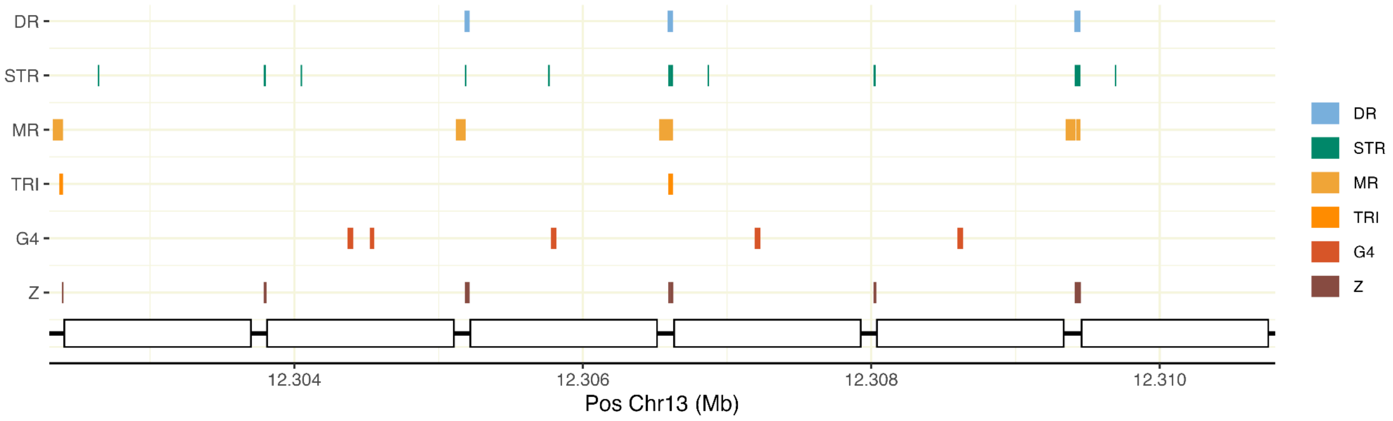

**Figure S16. Enrichment of non-B DNA at centromeres for alternative haplotypes**

Fold enrichment of non-B DNA densities at alternative haplotype centromeres compared to genome-wide densities. **(A)** chimpanzee, **(B)** bonobo, **(C)** gorilla, **(D)** Bornean orangutan, and **(E)** Sumatran orangutan. Underrepresentation of non-B DNA (values below 1) is shown in blue, while enrichment (values above 1) is shown in red. Centromeres that show a significant underrepresentation or enrichment compared to the genome-wide average density are marked in bold with '*' (two-sided randomization test, *p*<0.05). Abbreviations for non-B DNA are as in Fig. S4.

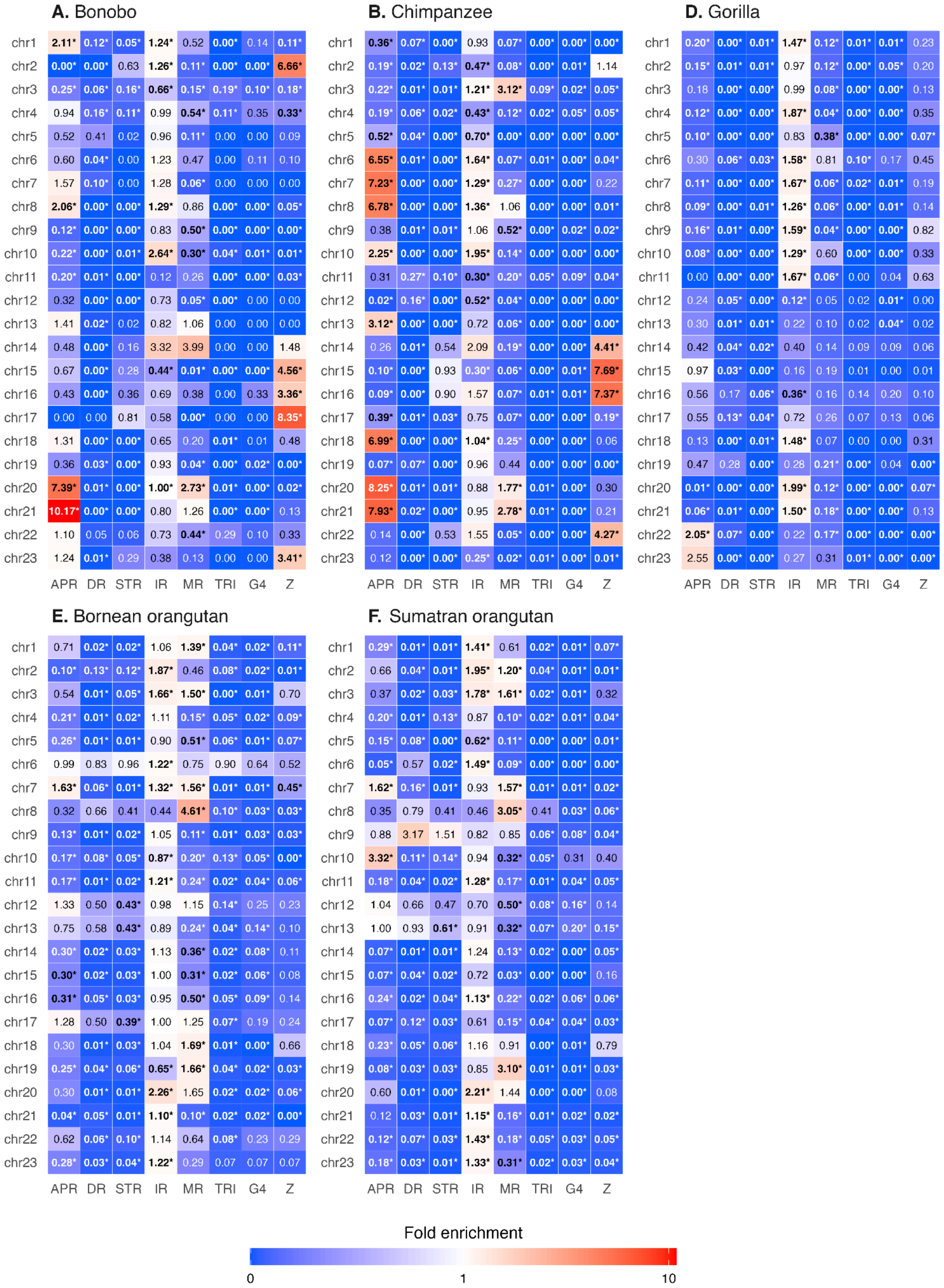

**Figure S17. Density of non-B DNA around centromeres**

Non-B DNA motif density of centromeres (showing 10 Mb around the midpoint of each centromere) normalized across the genome to a scale from 0-1 for each non-B type for **(A)** bonobo, **(B)** chimpanzee, **(C)** human, **(D)** gorilla, **(E)** Bornean orangutan, and **(F)** Sumatran orangutan. Both primary and alternative haplotypes are shown for non-human apes. Above the density blocks is a track with satellite repeat annotation using the color scheme from the UCSC Genome Browser. Satellite “Active αsat” defines the active centromere, and ”Mixed αsat” represents the annotations “MixedSuperFamily” and “mon/hor”.

**A.** Bonobo

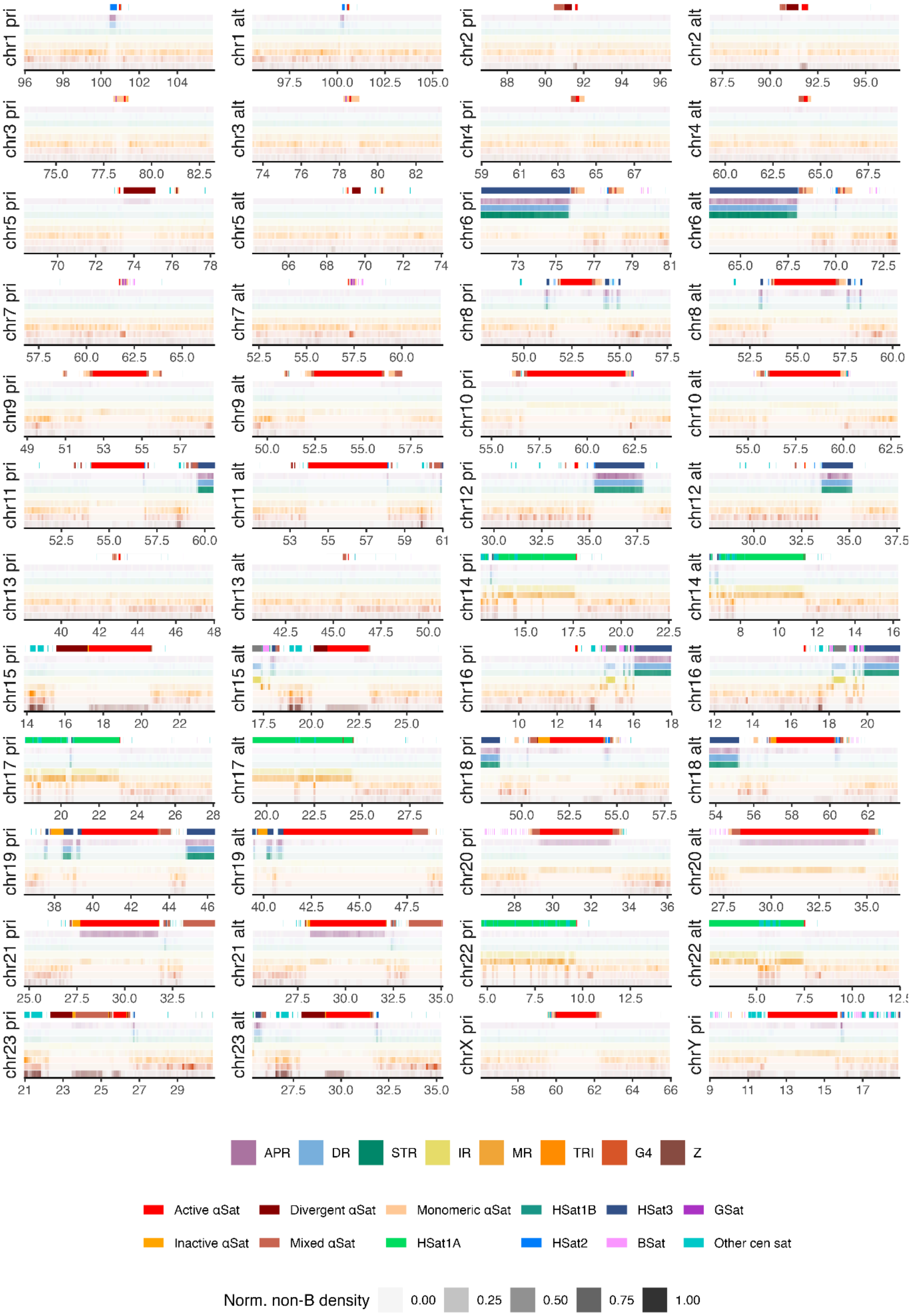

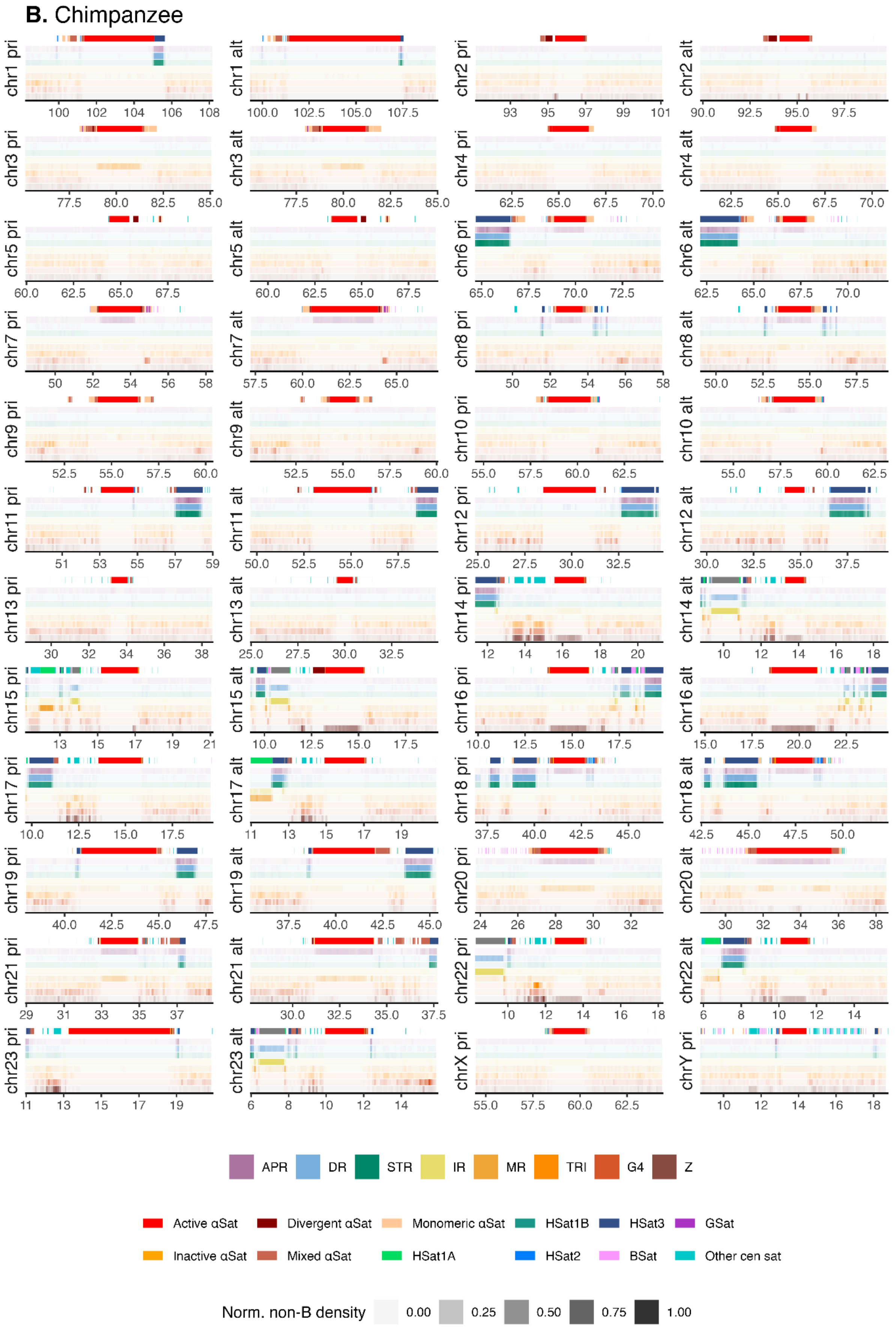
B. Chimpanzee
chr1 pri
100 102 104 106 108
chr1 alt
100.0 102.5 105.0 107.5
chr2 pri
93 95 97 99 101
chr2 alt
90.0 92.5 95.0 97.5
chr3 pri
77.5 80.0 82.5 85.0
chr3 alt
77.5 80.0 82.5 85.0
chr4 pri
62.5 65.0 67.5 70.0
chr4 alt
62.5 65.0 67.5 70.0
chr5 pri
60.0 62.5 65.0 67.5
chr5 alt
60.0 62.5 65.0 67.5
chr6 pri
65.0 67.5 70.0 72.5
chr6 alt
62.5 65.0 67.5 70.0
chr7 pri
50 52 54 56 58
chr7 alt
57.5 60.0 62.5 65.0
chr8 pri
50 52 54 56 58
chr8 alt
50.0 52.5 55.0 57.5
chr9 pri
52.5 55.0 57.5 60.0
chr9 alt
52.5 55.0 57.5 60.0
chr10 pri
55.0 57.5 60.0 62.5
chr10 alt
55.0 57.5 60.0 62.5
chr11 pri
51 53 55 57 59
chr11 alt
50.0 52.5 55.0 57.5
chr12 pri
25.0 27.5 30.0 32.5
chr12 alt
30.0 32.5 35.0 37.5
chr13 pri
30 32 34 36 38
chr13 alt
25.0 27.5 30.0 32.5
chr14 pri
12 14 16 18 20
chr14 alt
10 12 14 16 18
chr15 pri
13 15 17 19 21
chr15 alt
10.0 12.5 15.0 17.5
chr16 pri
10.0 12.5 15.0 17.5
chr16 alt
15.0 17.5 20.0 22.5
chr17 pri
10.0 12.5 15.0 17.5
chr17 alt
11 13 15 17 19
chr18 pri
37.5 40.0 42.5 45.0
chr18 alt
42.5 45.0 47.5 50.0
chr19 pri
40.0 42.5 45.0 47.5
chr19 alt
37.5 40.0 42.5 45.0
chr20 pri
24 26 28 30 32
chr20 alt
30 32 34 36 38
chr21 pri
29 31 33 35 37
chr21 alt
30.0 32.5 35.0 37.5
chr22 pri
10 12 14 16 18
chr22 alt
6 8 10 12 14
chr23 pri
11 13 15 17 19
chr23 alt
6 8 10 12 14
chrX pri
55.0 57.5 60.0 62.5
chrY pri
10 12 14 16 18
APR
DR
STR
IR
MR
TRI
G4
Z
Active αSat
Divergent αSat
Monomeric αSat
HSat1B
HSat3
GSat
Inactive αSat
Mixed αSat
HSat1A
HSat2
BSat
Other cen sat
Norm. non-B density
0.00
0.25
0.50
0.75
1.00

**C.** Human

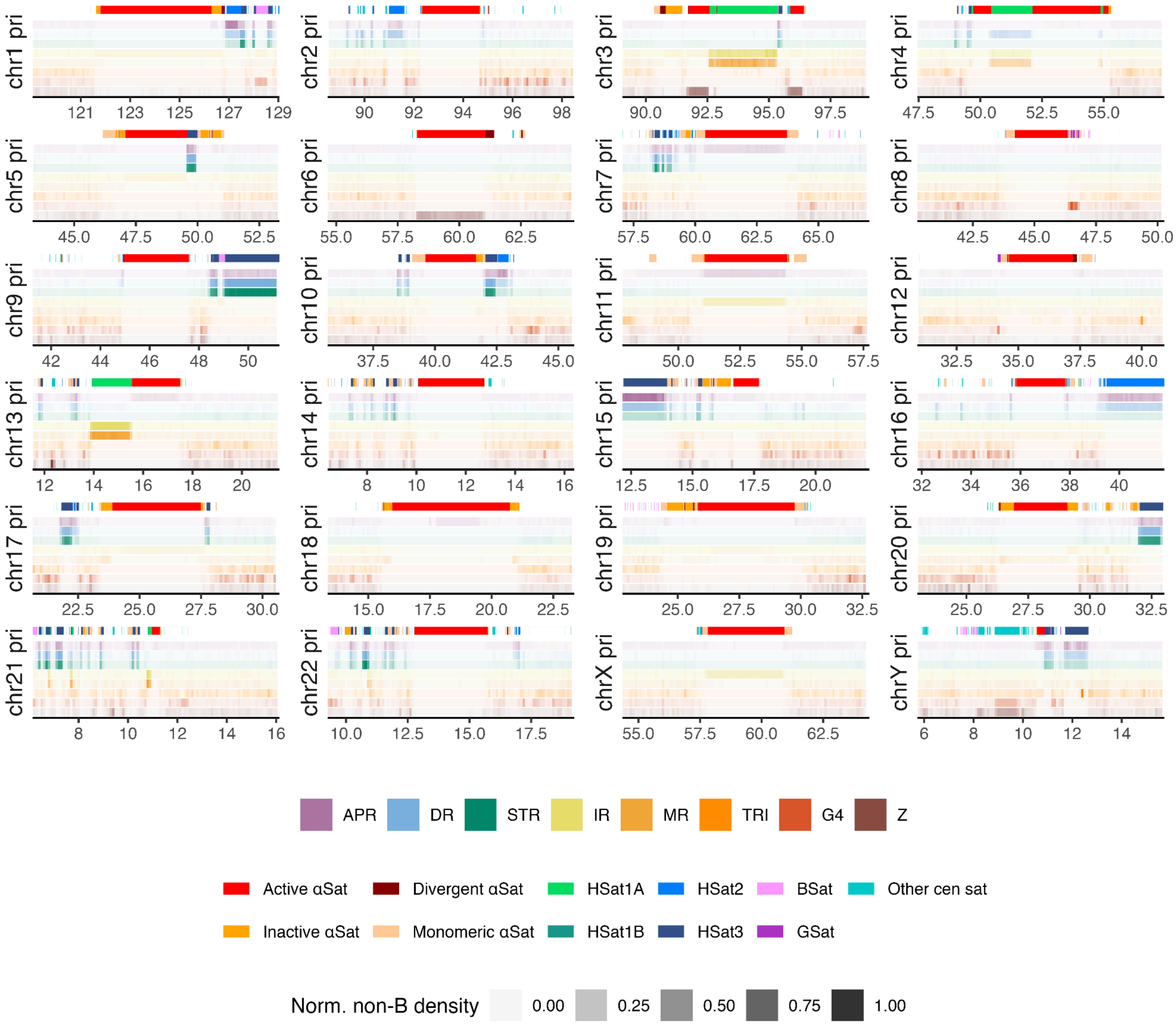
chr1 pri
chr2 pri
chr3 pri
chr4 pri
chr5 pri
chr6 pri
chr7 pri
chr8 pri
chr9 pri
chr10 pri
chr11 pri
chr12 pri
chr13 pri
chr14 pri
chr15 pri
chr16 pri
chr17 pri
chr18 pri
chr19 pri
chr20 pri
chr21 pri
chr22 pri
chrX pri
chrY pri
APR
DR
STR
IR
MR
TRI
G4
Z
Active αSat
Divergent αSat
HSat1A
HSat2
BSat
Other cen sat
Inactive αSat
Monomeric αSat
HSat1B
HSat3
GSat
Norm. non-B density
0.00
0.25
0.50
0.75
1.00

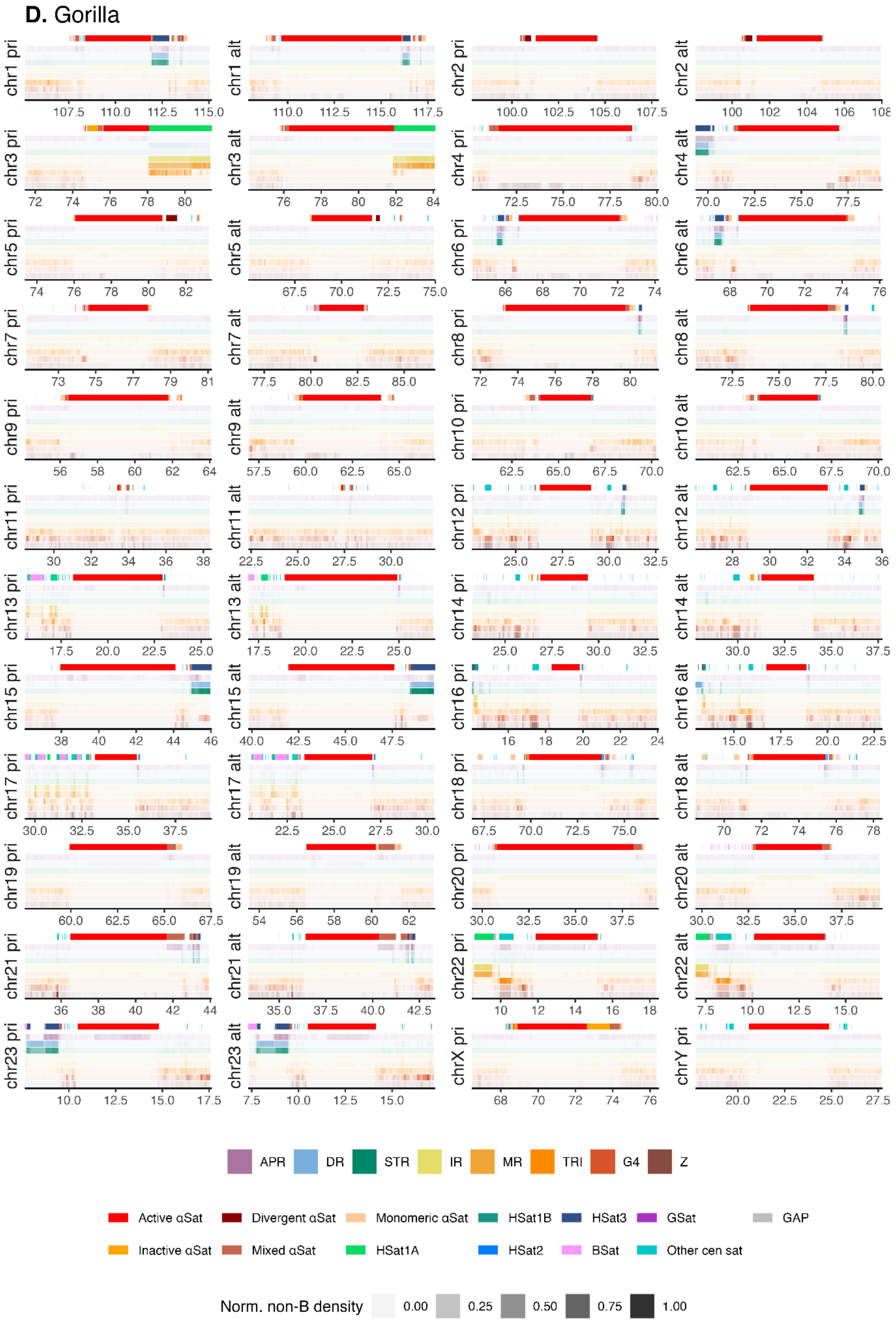
D. Gorilla
chr1 pri
chr1 alt
chr2 pri
chr2 alt
chr3 pri
chr3 alt
chr4 pri
chr4 alt
chr5 pri
chr5 alt
chr6 pri
chr6 alt
chr7 pri
chr7 alt
chr8 pri
chr8 alt
chr9 pri
chr9 alt
chr10 pri
chr10 alt
chr11 pri
chr11 alt
chr12 pri
chr12 alt
chr13 pri
chr13 alt
chr14 pri
chr14 alt
chr15 pri
chr15 alt
chr16 pri
chr16 alt
chr17 pri
chr17 alt
chr18 pri
chr18 alt
chr19 pri
chr19 alt
chr20 pri
chr20 alt
chr21 pri
chr21 alt
chr22 pri
chr22 alt
chr23 pri
chr23 alt
chrX pri
chrY pri
APR
DR
STR
IR
MR
TRI
G4
Z
Active αSat
Divergent αSat
Monomeric αSat
HSat1B
HSat3
GSat
GAP
Inactive αSat
Mixed αSat
HSat1A
HSat2
BSat
Other cen sat
Norm. non-B density
0.00
0.25
0.50
0.75
1.00

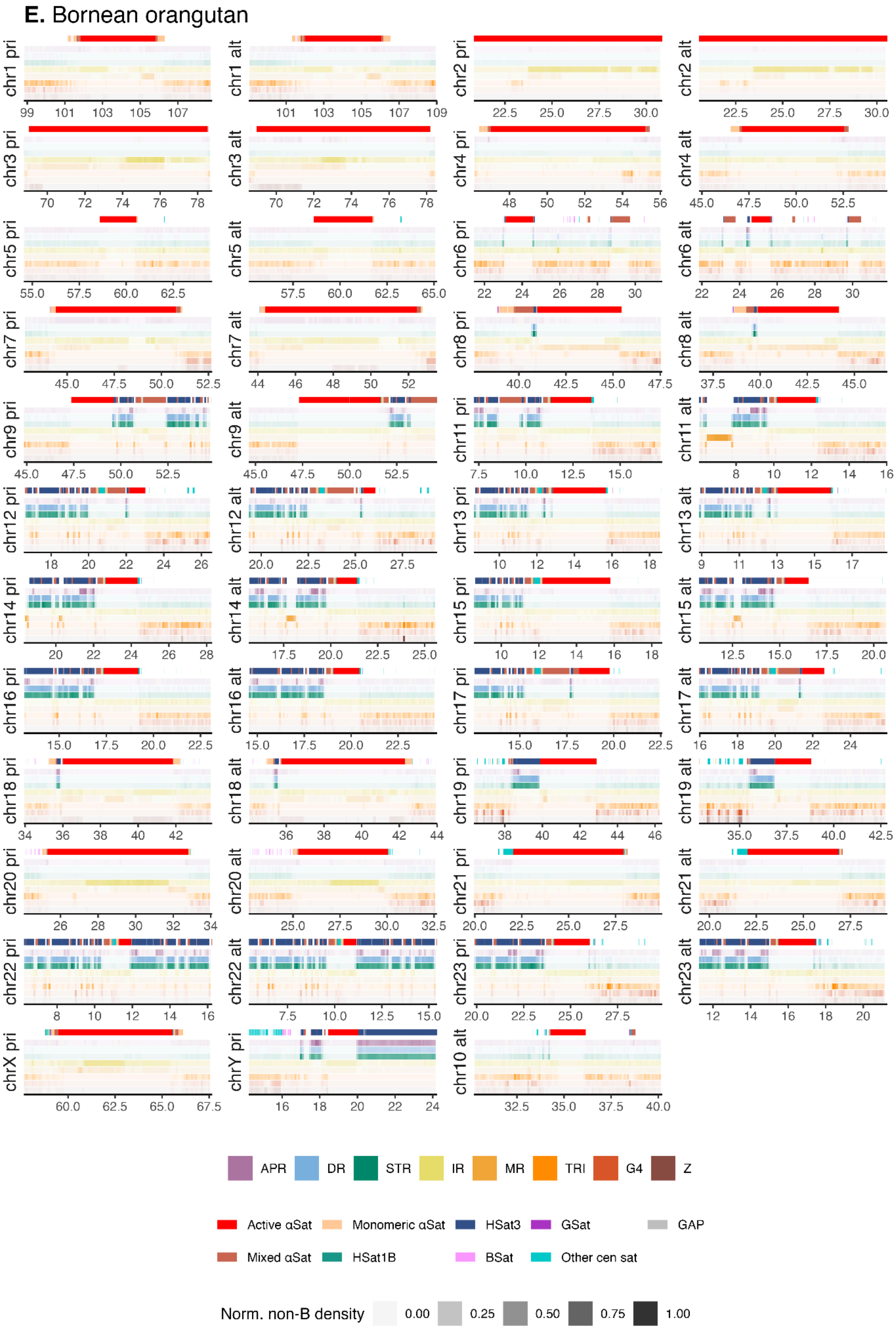

E. Bornean orangutan
chr1 pri
99 101 103 105 107
chr1 alt
101 103 105 107 109
chr2 pri
22.5 25.0 27.5 30.0
chr2 alt
22.5 25.0 27.5 30.0
chr3 pri
70 72 74 76 78
chr3 alt
70 72 74 76 78
chr4 pri
48 50 52 54 56
chr4 alt
45.0 47.5 50.0 52.5
chr5 pri
55.0 57.5 60.0 62.5
chr5 alt
57.5 60.0 62.5 65.0
chr6 pri
22 24 26 28 30
chr6 alt
22 24 26 28 30
chr7 pri
45.0 47.5 50.0 52.5
chr7 alt
44 46 48 50 52
chr8 pri
40.0 42.5 45.0 47.5
chr8 alt
37.5 40.0 42.5 45.0
chr9 pri
45.0 47.5 50.0 52.5
chr9 alt
45.0 47.5 50.0 52.5
chr11 pri
7.5 10.0 12.5 15.0
chr11 alt
8 10 12 14 16
chr12 pri
18 20 22 24 26
chr12 alt
20.0 22.5 25.0 27.5
chr13 pri
10 12 14 16 18
chr13 alt
9 11 13 15 17
chr14 pri
20 22 24 26 28
chr14 alt
17.5 20.0 22.5 25.0
chr15 pri
10 12 14 16 18
chr15 alt
12.5 15.0 17.5 20.0
chr16 pri
15.0 17.5 20.0 22.5
chr16 alt
15.0 17.5 20.0 22.5
chr17 pri
15.0 17.5 20.0 22.5
chr17 alt
16 18 20 22 24
chr18 pri
34 36 38 40 42
chr18 alt
36 38 40 42 44
chr19 pri
38 40 42 44 46
chr19 alt
35.0 37.5 40.0 42.5
chr20 pri
26 28 30 32 34
chr20 alt
25.0 27.5 30.0 32.5
chr21 pri
20.0 22.5 25.0 27.5
chr21 alt
20.0 22.5 25.0 27.5
chr22 pri
8 10 12 14 16
chr22 alt
7.5 10.0 12.5 15.0
chr23 pri
20.0 22.5 25.0 27.5
chr23 alt
12 14 16 18 20
chrX pri
60.0 62.5 65.0 67.5
chrY pri
16 18 20 22 24
chr10 alt
32.5 35.0 37.5 40.0
APR
DR
STR
IR
MR
TRI
G4
Z
Active αSat
Monomeric αSat
HSat3
GSat
GAP
Mixed αSat
HSat1B
BSat
Other cen sat
Norm. non-B density
0.00
0.25
0.50
0.75
1.00

**F.** Sumatran orangutan

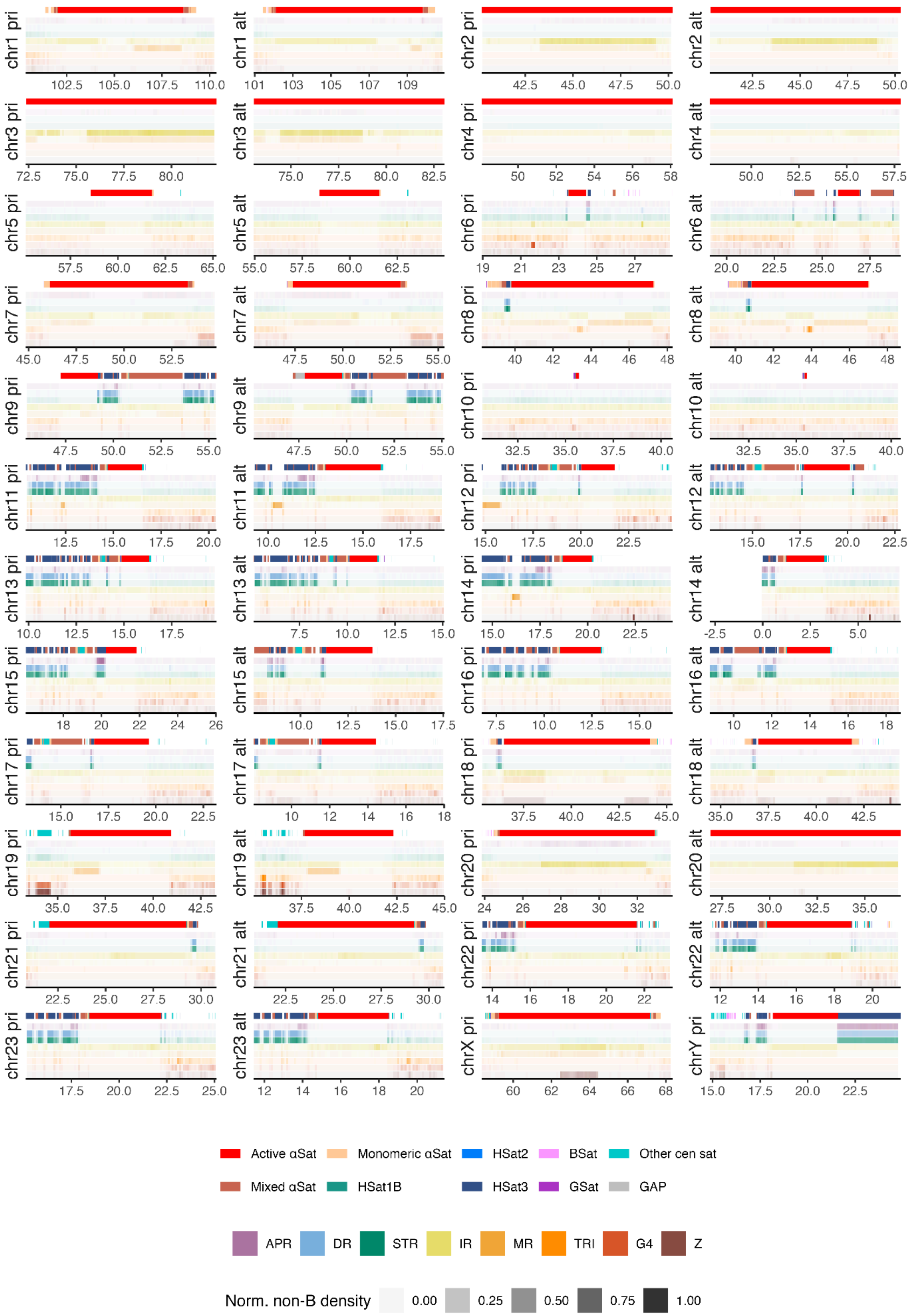

**Figure S18. Enrichment of non-B DNA at suprachromosomal families**
Enrichment of non-B DNA at centromeres belonging to different suprachromosomal families. Fold enrichment is from Fig. 7 and Fig. S16; all annotated centromeres from both primary and alternative haplotypes are used. The number of centromeres for each suprachromosomal family is given in the figure. Red dashed line at y=1 denotes the genome-wide average. Three bonobo centromeres with ambiguous SF annotation (primary chromosome 6 and both primary and alternative chr13) were removed for this analysis. Abbreviations for non-B DNA are as in Fig. S4.

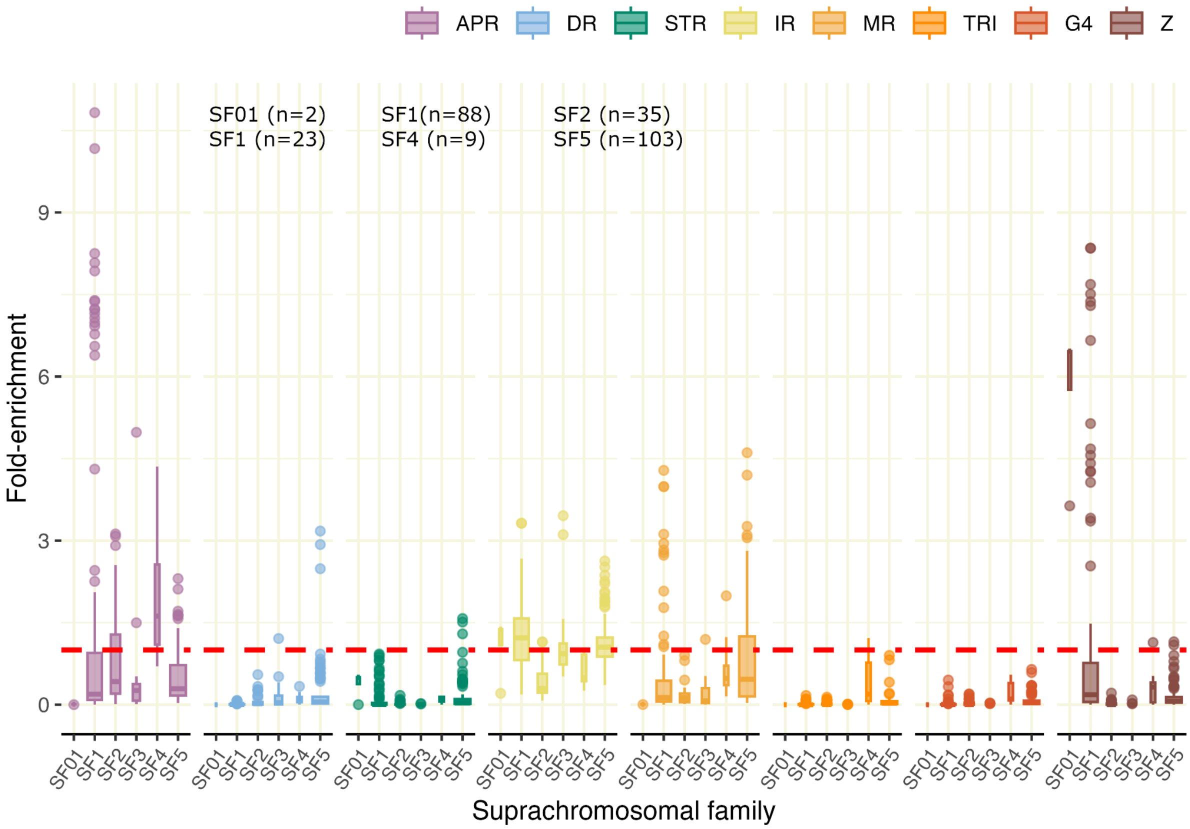

**Figure S19. Non-B DNA density at the Bornean orangutan chr10**
The centromeric region on chromosome 10 for Bornean orangutan. Alignment between the primary and alternative haplotype for the active centromere region. Only the alternative haplotype has an active centromere (marked with red bars). Abbreviations for non-B DNA are as in Fig. S4.

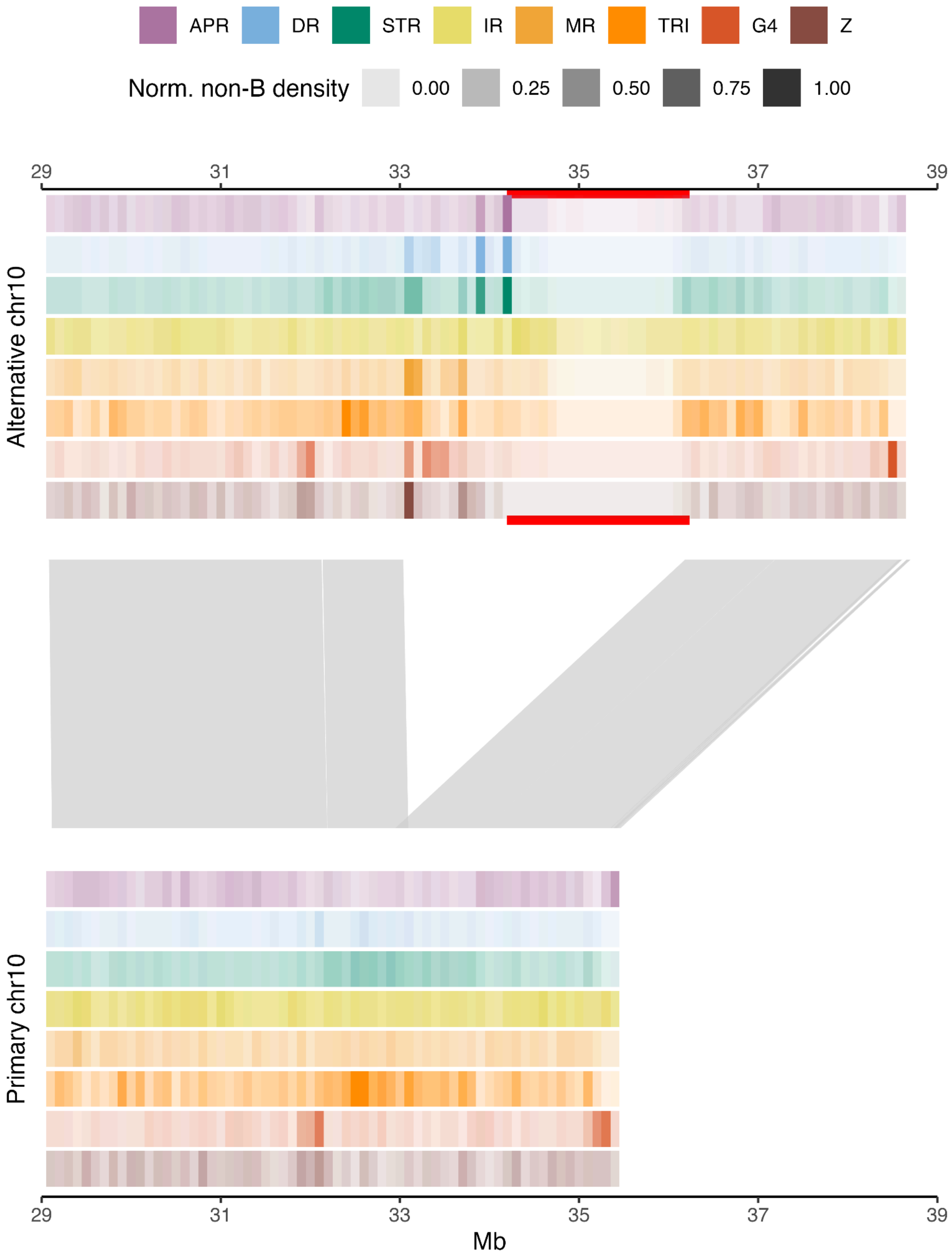

**Figure S20. Non-B DNA fold enrichment around centromeres with or without CENP-B binding motif.**
Centromeres with CENP-B motif are marked with "Yes", and centromeres without are marked with "No", on the x-axis. Enrichment at **(A)** 1-Mb flanks on the p-arms, (**B**) active centromeres, and (**C**) 1-Mb flanks on the q-arms. Red dashed line at y=1 denotes the genome-wide average.

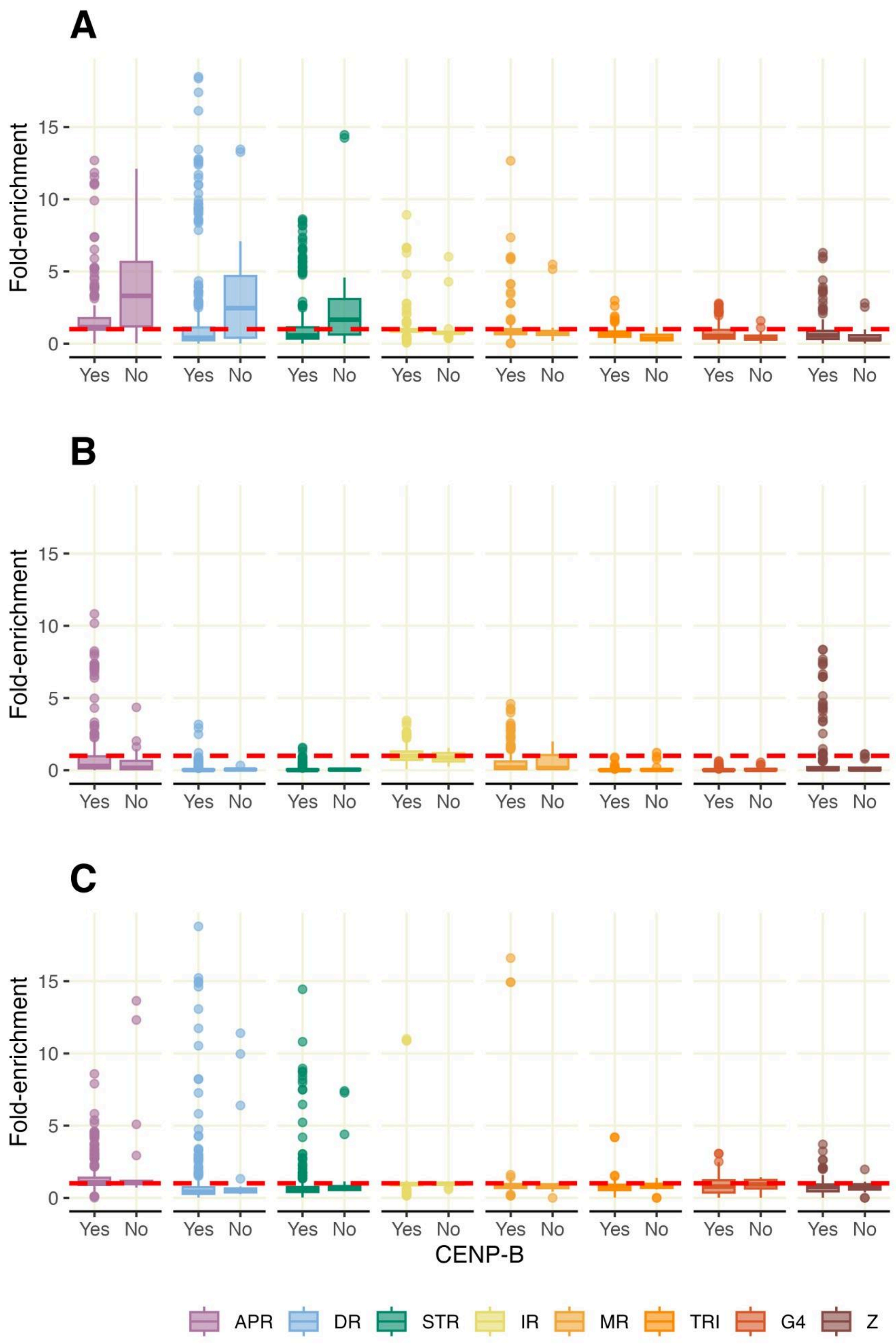

### Supplementary tables

##### Table S1. Non-B DNA annotations in ape genomes

Non-B DNA motif annotations (in Mb and percentage of total length) shown separately for each non-B DNA type, and for autosomes and sex chromosomes. APR: A-phased repeats; DR: direct repeats; STR: short tandem repeats; IR: inverted repeats; MR: mirror repeats; TRI: triplex motifs (a subset of mirror repeats); G4: G-quadruplexes; Z: Z-DNA.

| | | APR | | DR | | STR | | IR | | MR | | TRI | | G4 | | Z-DNA | | ALL | |
|---|---|---|---|---|---|---|---|---|---|---|---|---|---|---|---|---|---|---|---|
| **Chromosomes** | **Species** | **Mb** | **%** | **Mb** | **%** | **Mb** | **%** | **Mb** | **%** | **Mb** | **%** | **Mb** | **%** | **Mb** | **%** | **Mb** | **%** | **Mb** | **%** |
| Autosomes | Bonobo | 13.7 | 0.5 | 109.3 | 3.6 | 70.9 | 2.3 | 155.4 | 5.1 | 89.5 | 2.9 | 9.5 | 0.3 | 27.0 | 0.9 | 6.6 | 0.2 | 367.7 | 12.1 |
| | Chimpanzee | 11.8 | 0.4 | 88.9 | 3.0 | 53.7 | 1.8 | 146.0 | 4.9 | 87.3 | 2.9 | 9.6 | 0.3 | 26.7 | 0.9 | 6.6 | 0.2 | 335.9 | 11.2 |
| | Human | 13.8 | 0.5 | 81.5 | 2.8 | 65.1 | 2.2 | 138.2 | 4.8 | 82.4 | 2.8 | 9.8 | 0.3 | 26.2 | 0.9 | 6.6 | 0.2 | 320.2 | 11.0 |
| | Gorilla | 11.8 | 0.4 | 223.8 | 6.8 | 113.3 | 3.4 | 164.1 | 5.0 | 103.2 | 3.1 | 9.8 | 0.3 | 33.3 | 1.0 | 6.7 | 0.2 | 491.4 | 14.9 |
| | B. orangutan | 12.2 | 0.4 | 87.9 | 2.9 | 62.4 | 2.1 | 138.1 | 4.6 | 82.6 | 2.7 | 9.6 | 0.3 | 26.7 | 0.9 | 6.5 | 0.2 | 329.9 | 11.0 |
| | S. orangutan | 12.0 | 0.4 | 80.5 | 2.7 | 59.6 | 2.0 | 140.9 | 4.7 | 84.3 | 2.8 | 9.7 | 0.3 | 25.7 | 0.8 | 6.6 | 0.2 | 325.1 | 10.7 |
| | Siamang | 10.4 | 0.3 | 45.3 | 1.5 | 47.5 | 1.5 | 144.7 | 4.7 | 73.2 | 2.4 | 9.8 | 0.3 | 24.4 | 0.8 | 6.0 | 0.2 | 282.2 | 9.2 |
| Chr X | Bonobo | 0.6 | 0.4 | 3.0 | 1.9 | 2.4 | 1.5 | 7.5 | 4.7 | 4.9 | 3.0 | 0.5 | 0.3 | 1.1 | 0.7 | 0.3 | 0.2 | 15.4 | 9.6 |
| | Chimpanzee | 0.6 | 0.4 | 2.5 | 1.6 | 2.4 | 1.5 | 7.2 | 4.7 | 4.6 | 3.0 | 0.5 | 0.3 | 1.1 | 0.7 | 0.3 | 0.2 | 14.4 | 9.4 |
| | Human | 0.6 | 0.4 | 2.3 | 1.5 | 2.4 | 1.6 | 7.5 | 4.8 | 4.5 | 2.9 | 0.6 | 0.4 | 1.0 | 0.7 | 0.3 | 0.2 | 14.5 | 9.4 |
| | Gorilla | 0.6 | 0.4 | 7.7 | 4.4 | 2.4 | 1.4 | 7.5 | 4.2 | 4.8 | 2.7 | 0.5 | 0.3 | 1.1 | 0.6 | 0.3 | 0.2 | 20.3 | 11.4 |
| | B. orangutan | 0.6 | 0.4 | 2.3 | 1.5 | 2.4 | 1.5 | 7.6 | 4.7 | 5.0 | 3.1 | 0.5 | 0.3 | 1.1 | 0.7 | 0.3 | 0.2 | 15.2 | 9.5 |
| | S. orangutan | 0.6 | 0.4 | 2.4 | 1.4 | 2.5 | 1.5 | 7.7 | 4.7 | 5.1 | 3.1 | 0.5 | 0.3 | 1.0 | 0.6 | 0.3 | 0.2 | 15.4 | 9.5 |
| | Siamang | 0.6 | 0.4 | 2.2 | 1.3 | 2.3 | 1.4 | 7.6 | 4.6 | 4.2 | 2.5 | 0.5 | 0.3 | 1.0 | 0.6 | 0.3 | 0.2 | 14.5 | 8.8 |
| Chr Y | Bonobo | 0.6 | 1.3 | 6.8 | 14.5 | 4.7 | 10.0 | 1.9 | 4.0 | 1.7 | 3.6 | 0.1 | 0.3 | 0.3 | 0.6 | 0.1 | 0.2 | 10.8 | 23.1 |
| | Chimpanzee | 0.5 | 1.5 | 3.7 | 10.2 | 2.5 | 7.0 | 1.5 | 4.0 | 1.3 | 3.6 | 0.1 | 0.3 | 0.2 | 0.7 | 0.1 | 0.2 | 6.8 | 18.7 |
| | Human | 1.9 | 3.0 | 8.0 | 12.8 | 2.9 | 4.6 | 9.5 | 15.3 | 7.6 | 12.2 | 0.2 | 0.3 | 0.2 | 0.3 | 0.1 | 0.1 | 20.2 | 32.4 |
| | Gorilla | 0.6 | 0.8 | 12.0 | 17.8 | 6.8 | 10.1 | 9.3 | 13.7 | 8.4 | 12.5 | 0.1 | 0.2 | 0.4 | 0.5 | 0.1 | 0.1 | 25.5 | 37.9 |
| | B. orangutan | 0.6 | 1.2 | 4.2 | 8.4 | 2.2 | 4.5 | 2.1 | 4.2 | 1.5 | 3.1 | 0.1 | 0.3 | 0.3 | 0.5 | 0.1 | 0.2 | 8.4 | 17.0 |
| | S. orangutan | 1.1 | 1.7 | 8.1 | 12.0 | 4.2 | 6.1 | 2.7 | 4.0 | 1.9 | 2.8 | 0.2 | 0.2 | 0.3 | 0.4 | 0.1 | 0.1 | 14.2 | 20.9 |
| | Siamang | 0.3 | 0.9 | 1.3 | 4.2 | 1.0 | 3.5 | 1.3 | 4.3 | 0.9 | 2.8 | 0.1 | 0.3 | 0.3 | 1.0 | 0.1 | 0.2 | 3.7 | 12.2 |

**Table S2. Non-B DNA motif properties**
Total number of motifs, median length, and largest motif size for each species and motif type (primary haplotype only). Abbreviations for non-B DNA are as in Table S1.

| | Species | APR | DR | G4 | IR | MR | TRI | STR | Z |
|---|---|---|---|---|---|---|---|---|---|
| Number of motifs (genome-wide) | Chimpanzee | 502,595 | 2,974,304 | 752,601 | 7,162,803 | 2,156,713 | 425,119 | 3,609,972 | 448,683 |
| | Bonobo | 584,827 | 4,059,137 | 759,648 | 7,482,232 | 2,193,329 | 421,533 | 4,376,891 | 447,399 |
| | Human | 640,147 | 3,580,232 | 740,625 | 7,186,673 | 2,257,480 | 425,639 | 4,163,613 | 437,368 |
| | Gorilla | 500,671 | 7,223,474 | 852,151 | 8,056,679 | 2,539,384 | 426,325 | 5,626,242 | 446,513 |
| | B. orangutan | 517,920 | 3,780,971 | 738,899 | 7,060,589 | 2,113,530 | 430,576 | 4,207,235 | 447,042 |
| | S. orangutan | 531,382 | 3,609,826 | 745,395 | 7,246,561 | 2,145,577 | 434,344 | 4,171,360 | 454,282 |
| | Siamang | 431,987 | 1,713,830 | 695,659 | 7,637,314 | 1,923,960 | 424,303 | 3,235,785 | 411,074 |
| Median length (bp) | Chimpanzee | 25 | 32 | 33 | 15 | 41 | 25 | 14 | 12 |
| | Bonobo | 25 | 32 | 33 | 15 | 43 | 25 | 15 | 12 |
| | Human | 24 | 29 | 33 | 15 | 43 | 25 | 15 | 12 |
| | Gorilla | 25 | 39 | 34 | 15 | 50 | 25 | 18 | 12 |
| | B. orangutan | 25 | 28 | 34 | 15 | 40 | 25 | 15 | 12 |
| | S. orangutan | 25 | 28 | 34 | 15 | 40 | 25 | 15 | 12 |
| | Siamang | 25 | 28 | 33 | 15 | 38 | 25 | 13 | 12 |
| Maximum length (bp) | Chimpanzee | 193 | 23,346 | 13,652 | 1,691 | 559 | 353 | 10,768 | 184 |
| | Bonobo | 154 | 28,262 | 10,934 | 1,231 | 902 | 902 | 9,464 | 706 |
| | Human | 313 | 3,229 | 13,939 | 5,333 | 797 | 627 | 3,229 | 327 |
| | Gorilla | 113 | 8,425 | 14,159 | 590 | 1,428 | 737 | 8,294 | 207 |
| | B. orangutan | 1,652 | 30,334 | 21,788 | 4,108 | 1,615 | 1,051 | 9,043 | 458 |
| | S. orangutan | 293 | 9,686 | 13,763 | 13,100 | 2,397 | 2,397 | 9,176 | 660 |
| | Siamang | 131 | 11,471 | 12,945 | 1,082 | 2,395 | 1,891 | 11,471 | 315 |

**Table S3. Enrichment of non-B DNA motifs at repetitive sequences**

This is the same table as Table 2, but using two different subsample sizes for statistical testing. Cells for which 20 random subsamples (using either 10% or 50% of the data) all showed either enrichment or depletion are marked with "*". Abbreviations for non-B DNA are as in Table S1.

| Subsampling strategy | Species | APR | DR | STR | IR | MR | TRI | G4 | Z | all |
|---|---|---|---|---|---|---|---|---|---|---|
| 20 subsamples, each with 10% of the total regions | Bonobo | 1.31 | 5.90* | 5.52* | 1.04 | 1.90* | 6.42* | 0.55* | 2.15* | 1.53* |
| | Chimpanzee | 1.08 | 4.73* | 4.10* | 0.97 | 1.88* | 6.68* | 0.55* | 2.22* | 1.36* |
| | Human | 1.58 | 4.47* | 5.15* | 1.00 | 1.92* | 7.09* | 0.56* | 2.28* | 1.39* |
| | Gorilla | 0.91 | 11.32* | 7.86* | 1.05 | 2.12* | 5.60* | 0.66* | 1.92* | 2.04* |
| | Bornean orangutan | 1.10 | 4.93* | 4.65* | 0.86* | 1.76* | 6.59* | 0.58* | 2.15* | 1.33* |
| | Sumatran orangutan | 1.13 | 4.56* | 4.47* | 0.89* | 1.76* | 6.49* | 0.57* | 2.13* | 1.31* |
| | Siamang | 0.78* | 2.61* | 3.37* | 0.92* | 1.49* | 6.65* | 0.51* | 1.94* | 1.03 |
| 20 subsamples, each with 50% of the total regions | Bonobo | 1.31 | 5.90* | 5.52* | 1.04 | 1.90* | 6.42* | 0.55* | 2.15* | 1.53* |
| | Chimpanzee | 1.08 | 4.73* | 4.10* | 0.97 | 1.88* | 6.68* | 0.55* | 2.22* | 1.36* |
| | Human | 1.58* | 4.47* | 5.15* | 1.00 | 1.92* | 7.09* | 0.56* | 2.28* | 1.39* |
| | Gorilla | 0.91 | 11.32* | 7.86* | 1.05 | 2.12* | 5.60* | 0.66* | 1.92* | 2.04* |
| | Bornean orangutan | 1.10 | 4.93* | 4.65* | 0.86* | 1.76* | 6.59* | 0.58* | 2.15* | 1.33* |
| | Sumatran orangutan | 1.13 | 4.56* | 4.47* | 0.89* | 1.76* | 6.49* | 0.57* | 2.13* | 1.31* |
| | Siamang | 0.78* | 2.61* | 3.37* | 0.92* | 1.49* | 6.65* | 0.51* | 1.94* | 1.03* |

**Table S4. Non-B DNA motif enrichment at repeat groups**
Fold enrichment for different repeat groups in each species. Enrichment is calculated as non-B motif density divided by the genome-wide density. TEs: transposable elements, RNA: RNA repeats, Other: Simple repeats, Low complexity repeats, and Unknown repeats.

| Repeat Group | Species | APR | DR | STR | IR | MR | TRI | G4 | Z |
|---|---|---|---|---|---|---|---|---|---|
| TEs | Bonobo | 0.75 | 0.25 | 0.60 | 0.77 | 0.68 | 0.96 | 0.57 | 0.30 |
| | Chimpanzee | 0.85 | 0.31 | 0.79 | 0.80 | 0.69 | 0.96 | 0.57 | 0.29 |
| | Human | 0.66 | 0.32 | 0.66 | 0.78 | 0.68 | 0.93 | 0.56 | 0.29 |
| | Gorilla | 0.94 | 0.13 | 0.42 | 0.76 | 0.61 | 1.02 | 0.51 | 0.32 |
| | B. orangutan | 0.85 | 0.31 | 0.69 | 0.84 | 0.75 | 0.92 | 0.57 | 0.28 |
| | S. orangutan | 0.84 | 0.32 | 0.70 | 0.83 | 0.73 | 0.92 | 0.57 | 0.28 |
| | Siamang | 0.97 | 0.71 | 0.98 | 0.80 | 0.85 | 1.04 | 0.61 | 0.31 |
| Satellites | Bonobo | 2.33 | 4.76 | 3.20 | 1.53 | 1.57 | 0.52 | 0.62 | 0.67 |
| | Chimpanzee | 1.78 | 4.58 | 2.02 | 1.31 | 1.56 | 0.54 | 0.64 | 0.80 |
| | Human | 2.79 | 3.78 | 2.65 | 1.35 | 1.52 | 0.64 | 0.73 | 0.65 |
| | Gorilla | 1.02 | 4.10 | 3.16 | 1.35 | 1.67 | 0.32 | 0.76 | 0.42 |
| | B. orangutan | 1.89 | 4.76 | 2.81 | 0.94 | 1.20 | 0.58 | 0.70 | 0.77 |
| | S. orangutan | 1.88 | 4.17 | 2.48 | 1.03 | 1.23 | 0.60 | 0.71 | 0.78 |
| | Siamang | 0.71 | 1.03 | 0.69 | 1.17 | 0.37 | 0.29 | 0.37 | 0.37 |
| RNA | Bonobo | 0.29 | 0.19 | 0.46 | 0.86 | 0.82 | 0.63 | 2.15 | 0.89 |
| | Chimpanzee | 0.33 | 0.23 | 0.61 | 0.97 | 0.84 | 0.64 | 2.51 | 1.16 |
| | Human | 0.13 | 0.41 | 0.97 | 1.58 | 1.58 | 0.62 | 7.34 | 3.12 |
| | Gorilla | 0.38 | 0.09 | 0.28 | 0.88 | 0.62 | 0.43 | 1.27 | 0.78 |
| | B. orangutan | 0.26 | 0.40 | 0.61 | 1.10 | 0.75 | 0.53 | 2.40 | 0.94 |
| | S. orangutan | 0.22 | 0.37 | 0.76 | 1.22 | 0.84 | 0.50 | 4.04 | 1.44 |
| | Siamang | 0.33 | 0.48 | 0.70 | 0.97 | 0.96 | 0.67 | 1.89 | 0.70 |
| Other | Bonobo | 6.15 | 14.48 | 17.11 | 1.91 | 8.18 | 15.50 | 3.76 | 19.25 |
| | Chimpanzee | 4.66 | 14.99 | 19.69 | 2.77 | 11.78 | 21.62 | 5.44 | 27.05 |
| | Human | 4.55 | 14.35 | 13.36 | 3.28 | 9.59 | 13.25 | 3.06 | 16.70 |
| | Gorilla | 2.85 | 9.38 | 14.03 | 1.65 | 4.87 | 10.02 | 4.22 | 12.46 |
| | B. orangutan | 3.54 | 15.02 | 12.34 | 1.68 | 7.50 | 12.58 | 3.27 | 15.15 |
| | S. orangutan | 3.90 | 15.53 | 12.70 | 1.71 | 7.56 | 12.93 | 3.30 | 15.42 |
| | Siamang | 3.35 | 25.87 | 22.80 | 3.14 | 16.72 | 27.31 | 6.86 | 34.25 |

**Table S5. Methylation at repeats and satellites enriched in G4 motifs**
Repeats and satellites enriched in G4 motifs, and their average methylation scores within G4s and for the full annotated repeats/satellites, in two different human cell lines: HG002 (lymphoblastoid) and CHM13 (hydatidiform mole).

| | HG002 | | | | CHM13 | | | |
|---|---|---|---|---|---|---|---|---|
| **Repeat** | **Average methylation in G4 motifs** | **# CpG sites in G4s** | **Average methylation in full repeats** | **# CpG sites in full repeats** | **Average methylation in G4 motifs** | **# CpG sites in G4s** | **Average methylation in full repeats** | **# CpG sites in full repeats** |
| **Retroposon/SVA** | 0.89 | 7,401 | 0.91 | 234,912 | 0.75 | 10,839 | 0.78 | 239,192 |
| **rRNA** | 0.52 | 8,856 | 0.57 | 73,832 | 0.03 | 9,661 | 0.10 | 72,377 |
| **Satellite/acro_ACRO1** | 0.86 | 299 | 0.59 | 10,241 | 0.36 | 304 | 0.20 | 10,528 |
| **Satellite/centr_GSat** | 0.57 | 616 | 0.63 | 5,898 | 0.14 | 704 | 0.23 | 5,901 |
| **Satellite/centr_GSatII** | 0.59 | 1,169 | 0.63 | 10,920 | 0.16 | 1,191 | 0.27 | 10,983 |
| **Satellite/centr_GSatX** | 0.80 | 64 | 0.76 | 3,610 | 0.30 | 55 | 0.39 | 2,871 |
| **Satellite/centr_SST1** | 0.72 | 1,283 | 0.78 | 44,309 | 0.43 | 1,332 | 0.60 | 42,789 |
| **Satellite/subtelo_TAR1** | 0.60 | 431 | 0.75 | 13,887 | 0.30 | 468 | 0.39 | 7,737 |
| **Satellite_COMP-subunit_LSAU-BSat_rnd-1_family-1** | 0.77 | 2,931 | 0.78 | 46,344 | 0.48 | 3,236 | 0.48 | 47,492 |
| **Satellite_COMP-subunit_LSAU-BSat_rnd-1_family-4** | 0.64 | 1,506 | 0.73 | 13,009 | 0.23 | 1,570 | 0.36 | 13,344 |
| **Satellite_COMP-subunit_LSAU-BSat_rnd-6_family-5403** | 0.63 | 1,156 | 0.69 | 6,603 | 0.30 | 1,167 | 0.38 | 6,583 |
| **Satellite_HSat5** | 0.52 | 47 | 0.51 | 710 | 0.22 | 48 | 0.21 | 718 |
| **Satellite_LSAU** | 0.65 | 2,424 | 0.76 | 20,211 | 0.33 | 2,693 | 0.44 | 20,787 |
| **Satellite_MSR1** | 0.47 | 449 | 0.44 | 821 | 0.38 | 462 | 0.38 | 831 |
| **Satellite_Sat-VAR_rnd-6_family-3554** | 0.52 | 53 | 0.54 | 1,266 | 0.15 | 65 | 0.19 | 1,262 |
| **Genome-wide average** | 0.44 | 723,187 | 0.67 | 33,328,344 | 0.31 | 794,497 | 0.44 | 32,498,737 |

**Table S6. G4 formation at repeats and satellites**
Experimental data based on single-nuclei CUT&TAG from Hui et al. (2021) with BG4 peaks that indicate G4 formation in two different cancerous cell lines (K562 and U20S), lifted over from the previous human reference genome hg38 onto CHM13 (the T2T genome reference). Most satellites and repeats were not fully assembled in hg38, and hence the number of bases and G4s differ between the assembly versions. The third set of columns shows the number of BG4 peaks found in Hui et al. (2021) that could be lifted over to our satellites and repeats of interest. In the fourth column set, these numbers have been divided by the length of the lift-over sequence (higher numbers indicate more G4 formation per Mb of sequence). The highest density (marked in bold) was found in LSAU.

| | **Length (bp)** | | **N of G4s** | | **N of BG4 peak regions (lift-over to CHM13)** | | **N of BG4 peaks per Mb of sequence after lift-over** | |
|---|---|---|---|---|---|---|---|---|
| **Repeat** | CHM13 | hg38 (unique bp after liftOver) | CHM13 | hg38 (unique G4s after liftOver) | Cell line K562 | Cell line U20S | Cell line K562 | Cell line U20S |
| **Retroposon/SVA** | 4,409,546 | 3,663,447 | 9,390 | 5,749 | 37 | 49 | 10.1 | 13.4 |
| **rRNA** | 1,716,544 | 204,169 | 3,998 | 138 | 27 | 24 | 132.2 | 117.5 |
| **Satellite/acro_ACRO1** | 483,303 | 28,587 | 404 | 35 | 15 | 3 | 524.7 | 104.9 |
| **Satellite/centr_GSat** | 239,376 | 239,380 | 663 | 662 | 0 | 0 | 0.0 | 0.0 |
| **Satellite/centr_GSatII** | 275,973 | 179,121 | 633 | 369 | 5 | 7 | 27.9 | 39.1 |
| **Satellite/centr_GSatX** | 102,494 | 102,495 | 48 | 47 | 0 | 0 | 0.0 | 0.0 |
| **Satellite_HSat5** | 79,612 | 45,751 | 132 | 94 | 10 | 1 | 218.6 | 21.9 |
| **Satellite_LSAU** | 320,922 | 22,955 | 1,027 | 252 | 15 | 14 | **653.5** | **609.9** |
| **Satellite_MSR1** | 92,309 | 82,583 | 609 | 538 | 51 | 41 | 617.6 | 496.5 |
| **Satellite_Sat-VAR_rnd-6_family-3554** | 138,470 | 111,697 | 185 | 156 | 3 | 1 | 26.9 | 9.0 |
| **Satellite/centr_SST1** | 927,721 | 513,374 | 668 | 351 | 13 | 24 | 25.3 | 46.7 |
| **Satellite/subtelo_TAR1** | 135,875 | 56,429 | 131 | 56 | 11 | 22 | 194.9 | 389.9 |

**Table S7. Non-B DNA enrichment at SST1**

Fold enrichment of non-B motifs at the satellite SST1, as compared to the genome-wide average. The SST1 subfamily annotations are taken from de Lima et al. 2024. Fold enrichment is calculated separately for annotated repeat units and for the intermediate sequence (“spacer”) between annotated satellites (for repeat units less than 1 kb apart). Average spacer lengths are 135 bp for sf1, 815 bp for sf2, and 484 bp for sf3. The extreme fold enrichment in Z-DNA of >97× for spacers on sf1 corresponds to a density of 0.22 (i.e., 22% of bases annotated as potentially Z-DNA forming), compared to a genome-wide density of 0.0023 genome-wide).

| Region | SST1 subfamily | Chromosomes | Fold enrichment compared to genome-wide | | | | | | | |
|---|---|---|---|---|---|---|---|---|---|---|
| | | | APR | DR | STR | IR | MR | TRI | G4 | Z-DNA |
| **Annotated satellite units** | SST1 SF1 | chr13, chr14, chr21 | 0.00 | 0.01 | 0.34 | 0.69 | 0.28 | 0.00 | 3.13 | 0.27 |
| | SST1 SF2 | chr4, chr17, chr19 | 0.04 | 0.31 | 0.40 | 0.46 | 0.52 | 0.07 | 5.01 | 0.66 |
| | SST1 SF3 | chrY | 1.05 | 0.06 | 0.10 | 0.69 | 0.31 | 0.03 | 0.27 | 0.12 |
| **Non-satellite sequence within satellite arrays (monomeres <1kb apart)** | SST1 SF1 | chr13, chr14, chr21 | 0.00 | 6.86 | 5.36 | 0.00 | 14.69 | 9.46 | 0.41 | 97.14 |
| | SST1 SF2 | chr4, chr17, chr19 | 0.10 | 2.33 | 3.53 | 0.37 | 3.68 | 4.48 | 0.14 | 27.31 |
| | SST1 SF3 | chrY | 0.40 | 0.42 | 0.92 | 0.49 | 0.62 | 1.01 | 0.25 | 3.16 |

**Table S8. Non-B DNA enrichment at the centromeres, suprachromosomal families, and depending on CENP-B box presence/absence**
Centromere enrichment (same data as in Fig. 7 and Fig. S16) along with haplotype, Suprachromosomal family (SF), and CENP-B motif presence/absence, sorted after SF type. Abbreviations as in Table S1.

| Species | Full chromosome name | SF | Hap | Fold enrichment | | | | | | | | CENPB in centromeres |
|---|---|---|---|---|---|---|---|---|---|---|---|---|
| | | | | APR | DR | STR | IR | MR | TRI | G4 | Z | |
| Human | chr3 | SF01 | Pri | 0.01* | 0.00* | 0.50* | 1.38* | 0.01* | 0.00* | 0.00* | 6.46* | Yes |
| Human | chr6 | SF01 | Pri | 0.00* | 0.00* | 0.00* | 0.21* | 0.00* | 0.00* | 0.00* | 3.64* | Yes |
| Chimpanzee | chr2_hap1_hsa3 | SF1 | Pri | 0.12* | 0.08* | 0.12* | 0.39* | 0.07* | 0.06* | 0.01* | 1.05 | Yes |
| Chimpanzee | chr3_hap1_hsa4 | SF1 | Pri | 0.11* | 0.00* | 0.00* | 1.11 | 4.28* | 0.08* | 0.00* | 0.05* | Yes |
| Chimpanzee | chr6_hap1_hsa7 | SF1 | Pri | 7.37* | 0.00* | 0.00* | 1.72* | 0.01* | 0.01* | 0.00* | 0.03* | Yes |
| Chimpanzee | chr7_hap1_hsa8 | SF1 | Pri | 6.92* | 0.01* | 0.00* | 1.25* | 0.35* | 0.00* | 0.00* | 0.08* | Yes |
| Chimpanzee | chr8_hap1_hsa10 | SF1 | Pri | 6.39* | 0.00* | 0.00* | 1.30* | 1.1 | 0.00* | 0.00* | 0.03* | Yes |
| Chimpanzee | chr10_hap1_hsa12 | SF1 | Pri | 2.45* | 0.00* | 0.00* | 2.50* | 0.11* | 0.00* | 0.00* | 0.00* | Yes |
| Chimpanzee | chr14_hap1_hsa13 | SF1 | Pri | 0.21* | 0.01* | 0.62 | 2.47 | 0.05* | 0.00* | 0.00* | 5.14* | Yes |
| Chimpanzee | chr15_hap1_hsa14 | SF1 | Pri | 0.27* | 0.01* | 0.09* | 0.26* | 0.05* | 0.04* | 0.02* | 0.72 | Yes |
| Chimpanzee | chr16_hap1_hsa15 | SF1 | Pri | 0.1 | 0.00* | 0.87 | 1.47* | 0.21* | 0.01* | 0.01* | 7.30* | Yes |
| Chimpanzee | chr17_hap1_hsa18 | SF1 | Pri | 0.21* | 0.01* | 0.01* | 0.75 | 0.11* | 0.00* | 0.00* | 0.09* | Yes |
| Chimpanzee | chr18_hap1_hsa16 | SF1 | Pri | 7.07* | 0.00* | 0.00* | 1.09* | 0.18* | 0.00* | 0.00* | 0.13 | Yes |
| Chimpanzee | chr20_hap1_hsa19 | SF1 | Pri | 7.23* | 0.00* | 0.01* | 1.07* | 2.83* | 0.00* | 0.00* | 0.08* | Yes |
| Chimpanzee | chr21_hap1_hsa20 | SF1 | Pri | 8.08* | 0.00* | 0.00* | 1.01 | 2.95* | 0.01* | 0.00* | 0.09 | Yes |
| Chimpanzee | chr22_hap1_hsa21 | SF1 | Pri | 0.13 | 0.00* | 0.51 | 1.44 | 0.05* | 0.00* | 0.00* | 4.26* | Yes |
| Chimpanzee | chr23_hap1_hsa22 | SF1 | Pri | 0.23 | 0.01* | 0.00* | 0.19* | 0.04* | 0.03* | 0.00* | 0.03* | Yes |
| Bonobo | chr2_mat_hsa3 | SF1 | Pri | 0.08* | 0.00* | 0.67 | 1.61* | 0.10* | 0.00* | 0.00* | 7.51* | Yes |
| Bonobo | chr8_mat_hsa10 | SF1 | Pri | 1.48 | 0.01* | 0.00* | 1.22* | 0.68* | 0.00* | 0.00* | 0.21 | Yes |
| Bonobo | chr10_mat_hsa12 | SF1 | Pri | 0.24* | 0.05* | 0.01* | 2.67* | 0.20* | 0.01* | 0.03* | 0.03* | Yes |
| Bonobo | chr15_mat_hsa14 | SF1 | Pri | 0.25 | 0.01* | 0.38 | 0.47* | 0.04* | 0.00* | 0.04* | 4.07* | Yes |
| Bonobo | chr16_pat_hsa15 | SF1 | Pri | 0.66 | 0.00* | 0.24 | 0.68 | 0.57 | 0 | 0.45 | 2.54* | Yes |
| Bonobo | chr18_pat_hsa16 | SF1 | Pri | 1.22 | 0.00* | 0.00* | 0.67 | 0.28 | 0.00* | 0.00* | 0.74 | Yes |
| Bonobo | chr20_pat_hsa19 | SF1 | Pri | 7.16* | 0.00* | 0.00* | 1.05* | 2.08* | 0.00* | 0.00* | 0.01* | Yes |
| Bonobo | chr21_mat_hsa20 | SF1 | Pri | 10.83* | 0.00* | 0.00* | 0.75 | 0.82 | 0.00* | 0.00* | 0.03* | Yes |
| Bonobo | chr23_pat_hsa22 | SF1 | Pri | 0.46 | 0.00* | 0.39 | 0.27 | 0.11 | 0 | 0 | 4.68 | Yes |
| Human | chr1 | SF1 | Pri | 0.01* | 0.00* | 0.00* | 1.30* | 0.05 | 0 | 0 | 0.07 | Yes |
| Human | chr5 | SF1 | Pri | 0.00* | 0.00* | 0.00* | 1.58* | 0.03* | 0.00* | 0.00* | 0.07* | Yes |
| Human | chr7 | SF1 | Pri | 4.31* | 0.01* | 0.00* | 0.37* | 0.01* | 0.00* | 0.00* | 0.01* | Yes |
| Human | chr10 | SF1 | Pri | 0.02* | 0.00* | 0.00* | 0.40* | 0.03* | 0.00* | 0.00* | 0.01* | Yes |
| Human | chr12 | SF1 | Pri | 0.03* | 0.00* | 0.00* | 0.76* | 0.41* | 0.17* | 0.00* | 0.03* | Yes |
| Human | chr16 | SF1 | Pri | 0.14* | 0.00* | 0.00* | 0.85 | 0.92 | 0 | 0 | 0.05 | Yes |
| Human | chr19 | SF1 | Pri | 0.02* | 0.00* | 0.00* | 1.07* | 0.14* | 0.00* | 0.00* | 0.03* | Yes |
| Gorilla | chr1_pat_hsa1 | SF1 | Pri | 0.19* | 0.00* | 0.01* | 1.54* | 0.18* | 0.01* | 0.02* | 0.12* | Yes |
| Gorilla | chr2_pat_hsa3 | SF1 | Pri | 0.18 | 0.01* | 0.01* | 0.91 | 0.13* | 0.00* | 0.06 | 0.22 | Yes |
| Gorilla | chr3_pat_hsa4 | SF1 | Pri | 0.64 | 0.01* | 0.01* | 0.98 | 0.13* | 0.00* | 0 | 0.04 | Yes |
| Gorilla | chr4_pat_hsa17x5 | SF1 | Pri | 0.36 | 0.01* | 0.00* | 1.65* | 0.04* | 0.01* | 0.01* | 1.21 | Yes |
| Gorilla | chr5_mat_hsa6 | SF1 | Pri | 0.12* | 0.00* | 0.00* | 1.31* | 0.20* | 0.00* | 0.00* | 0.23 | Yes |
| Gorilla | chr6_mat_hsa7 | SF1 | Pri | 0.32 | 0.07* | 0.04* | 1.45* | 0.6 | 0.10* | 0.18* | 0.52 | Yes |
| Gorilla | chr7_pat_hsa8 | SF1 | Pri | 0.06* | 0.00* | 0.01* | 1.68* | 0.08* | 0.01* | 0.00* | 0.18 | Yes |
| Gorilla | chr8_pat_hsa10 | SF1 | Pri | 0.06* | 0.01* | 0.02* | 1.15* | 0.04* | 0.00* | 0.00* | 0.24 | Yes |
| Gorilla | chr9_pat_hsa11 | SF1 | Pri | 0.03* | 0.00* | 0.01* | 1.61* | 0.05* | 0.00* | 0.00* | 0.43 | Yes |
| Gorilla | chr10_mat_hsa12 | SF1 | Pri | 0.08* | 0.01* | 0.01* | 1.17* | 0.24* | 0.00* | 0.00* | 0.78 | Yes |
| Gorilla | chr11_mat_hsa2b | SF1 | Pri | 0 | 0.00* | 0.00* | 1.69* | 0.06* | 0.00* | 0.04 | 0.61 | Yes |

| | | | | | | | | | | | | |
|---|---|---|---|---|---|---|---|---|---|---|---|---|
| Gorilla | chr18_pat_hsa16 | SF1 | Pri | 0.13 | 0.00* | 0.01* | 1.46* | 0.07 | 0 | 0 | 0.3 | Yes |
| Gorilla | chr20_mat_hsa19 | SF1 | Pri | 0.01* | 0.00* | 0.00* | 1.92* | 0.02* | 0.00* | 0.00* | 0.03* | Yes |
| Gorilla | chr21_pat_hsa20 | SF1 | Pri | 0.05* | 0.02* | 0.00* | 1.47* | 0.15* | 0.00* | 0.00* | 0.15 | Yes |
| Gorilla | chrX_mat_hsaX | SF1 | Pri | 0.44 | 0.01* | 0.01* | 0.76 | 0.09* | 0.02* | 0.02* | 0.07* | Yes |
| Gorilla | chrY_pat_hsaY | SF1 | Pri | 0.09 | 0.00* | 0.08 | 1.28 | 0.09 | 0.03* | 0 | 0.04 | Yes |
| Chimpanzee | chr2_hap2_hsa3 | SF1 | Alt | 0.19* | 0.02* | 0.13* | 0.47* | 0.08* | 0.00* | 0.01* | 1.14 | Yes |
| Chimpanzee | chr3_hap2_hsa4 | SF1 | Alt | 0.22* | 0.01* | 0.01* | 1.21* | 3.12* | 0.09* | 0.02* | 0.05* | Yes |
| Chimpanzee | chr6_hap2_hsa7 | SF1 | Alt | 6.55* | 0.01* | 0.00* | 1.64* | 0.07* | 0.01* | 0.00* | 0.04* | Yes |
| Chimpanzee | chr7_hap2_hsa8 | SF1 | Alt | 7.23* | 0.00* | 0.00* | 1.29* | 0.27* | 0.00* | 0.00* | 0.22 | Yes |
| Chimpanzee | chr8_hap2_hsa10 | SF1 | Alt | 6.78* | 0.00* | 0.00* | 1.36* | 1.06 | 0.00* | 0.00* | 0.01* | Yes |
| Chimpanzee | chr10_hap2_hsa12 | SF1 | Alt | 2.25* | 0.00* | 0.00* | 1.95* | 0.14* | 0.00* | 0.00* | 0.00* | Yes |
| Chimpanzee | chr14_hap2_hsa13 | SF1 | Alt | 0.26 | 0.01* | 0.54 | 2.09 | 0.19* | 0.00* | 0.00* | 4.41* | Yes |
| Chimpanzee | chr15_hap2_hsa14 | SF1 | Alt | 0.10* | 0.00* | 0.93 | 0.30* | 0.06* | 0.00* | 0.01* | 7.69* | Yes |
| Chimpanzee | chr16_hap2_hsa15 | SF1 | Alt | 0.09* | 0.00* | 0.9 | 1.57 | 0.07* | 0.01* | 0.01* | 7.37* | Yes |
| Chimpanzee | chr17_hap2_hsa18 | SF1 | Alt | 0.39* | 0.01* | 0.03* | 0.75 | 0.07* | 0.00* | 0.00* | 0.19* | Yes |
| Chimpanzee | chr18_hap2_hsa16 | SF1 | Alt | 6.99* | 0.00* | 0.00* | 1.04* | 0.25* | 0.00* | 0.00* | 0.06 | Yes |
| Chimpanzee | chr20_hap2_hsa19 | SF1 | Alt | 8.25* | 0.01* | 0.01* | 0.88 | 1.77* | 0.01* | 0.00* | 0.3 | Yes |
| Chimpanzee | chr21_hap2_hsa20 | SF1 | Alt | 7.93* | 0.02* | 0.00* | 0.95 | 2.78* | 0.01* | 0.00* | 0.21 | Yes |
| Chimpanzee | chr22_hap2_hsa21 | SF1 | Alt | 0.14 | 0.00* | 0.53 | 1.55 | 0.05* | 0.00* | 0.00* | 4.27* | Yes |
| Chimpanzee | chr23_hap2_hsa22 | SF1 | Alt | 0.12 | 0.00* | 0.00* | 0.25* | 0.02* | 0.01* | 0.00* | 0.01* | Yes |
| Bonobo | chr2_pat_hsa3 | SF1 | Alt | 0.00* | 0.00* | 0.63 | 1.26* | 0.11* | 0.00* | 0.00* | 6.66* | Yes |
| Bonobo | chr6_pat_hsa7 | SF1 | Alt | 0.6 | 0.04* | 0 | 1.23 | 0.47 | 0 | 0.11 | 0.1 | No |
| Bonobo | chr8_pat_hsa10 | SF1 | Alt | 2.06* | 0.00* | 0.00* | 1.29* | 0.86 | 0.00* | 0.00* | 0.05* | Yes |
| Bonobo | chr10_pat_hsa12 | SF1 | Alt | 0.22* | 0.00* | 0.01* | 2.64* | 0.30* | 0.04* | 0.01* | 0.01* | Yes |
| Bonobo | chr14_mat_hsa13 | SF1 | Alt | 0.48 | 0.00* | 0.16 | 3.32 | 3.99 | 0 | 0 | 1.48 | Yes |
| Bonobo | chr15_pat_hsa14 | SF1 | Alt | 0.67 | 0.00* | 0.28 | 0.44* | 0.01* | 0.00* | 0.00* | 4.56* | Yes |
| Bonobo | chr16_mat_hsa15 | SF1 | Alt | 0.43 | 0.00* | 0.36 | 0.69 | 0.38 | 0 | 0.33 | 3.36* | Yes |
| Bonobo | chr17_pat_hsa18 | SF1 | Alt | 0 | 0 | 0.81 | 0.58 | 0.00* | 0 | 0 | 8.35* | Yes |
| Bonobo | chr18_mat_hsa16 | SF1 | Alt | 1.31 | 0.00* | 0.00* | 0.65 | 0.2 | 0.01* | 0.01 | 0.48 | Yes |
| Bonobo | chr20_mat_hsa19 | SF1 | Alt | 7.39* | 0.01* | 0.00* | 1.00* | 2.73* | 0.01* | 0.00* | 0.02* | Yes |
| Bonobo | chr21_pat_hsa20 | SF1 | Alt | 10.17* | 0.00* | 0.00* | 0.8 | 1.26 | 0.00* | 0.00* | 0.13 | Yes |
| Bonobo | chr23_mat_hsa22 | SF1 | Alt | 1.24 | 0.01* | 0.29 | 0.38 | 0.13 | 0 | 0 | 3.41* | Yes |
| Gorilla | chr1_mat_hsa1 | SF1 | Alt | 0.20* | 0.00* | 0.01* | 1.47* | 0.12* | 0.01* | 0.01* | 0.23 | Yes |
| Gorilla | chr2_mat_hsa3 | SF1 | Alt | 0.15* | 0.01* | 0.01* | 0.97 | 0.12* | 0.00* | 0.05* | 0.2 | Yes |
| Gorilla | chr3_mat_hsa4 | SF1 | Alt | 0.18 | 0.00* | 0.00* | 0.99 | 0.08* | 0.00* | 0.00* | 0.13 | Yes |
| Gorilla | chr4_mat_hsa17x5 | SF1 | Alt | 0.12* | 0.00* | 0.00* | 1.87* | 0.04* | 0.00* | 0.00* | 0.35 | Yes |
| Gorilla | chr5_pat_hsa6 | SF1 | Alt | 0.10* | 0.00* | 0.00* | 0.83 | 0.38* | 0.00* | 0.00* | 0.07* | Yes |
| Gorilla | chr6_pat_hsa7 | SF1 | Alt | 0.3 | 0.06* | 0.03* | 1.58* | 0.81 | 0.10* | 0.17 | 0.45 | Yes |
| Gorilla | chr7_mat_hsa8 | SF1 | Alt | 0.11* | 0.00* | 0.00* | 1.67* | 0.06* | 0.02* | 0.01* | 0.19 | Yes |
| Gorilla | chr8_mat_hsa10 | SF1 | Alt | 0.09* | 0.00* | 0.01* | 1.26* | 0.06* | 0.00* | 0.01* | 0.14 | Yes |
| Gorilla | chr9_mat_hsa11 | SF1 | Alt | 0.16* | 0.01* | 0.00* | 1.59* | 0.04* | 0.00* | 0.00* | 0.82 | Yes |
| Gorilla | chr10_pat_hsa12 | SF1 | Alt | 0.08* | 0.00* | 0.00* | 1.29* | 0.6 | 0.00* | 0.00* | 0.33 | Yes |
| Gorilla | chr11_pat_hsa2b | SF1 | Alt | 0 | 0.00* | 0.00* | 1.67* | 0.06* | 0 | 0.04 | 0.63 | Yes |
| Gorilla | chr18_mat_hsa16 | SF1 | Alt | 0.13 | 0.00* | 0.01* | 1.48* | 0.07 | 0 | 0 | 0.31 | Yes |
| Gorilla | chr20_pat_hsa19 | SF1 | Alt | 0.01* | 0.00* | 0.00* | 1.99* | 0.12* | 0.00* | 0.00* | 0.07* | Yes |
| Gorilla | chr21_mat_hsa20 | SF1 | Alt | 0.06* | 0.01* | 0.00* | 1.50* | 0.18* | 0.00* | 0.00* | 0.13 | Yes |
| Bonobo | chr6_mat_hsa7 | SF1,SF01 | Pri | 0.77 | 0.06* | 0 | 1.22 | 0.11 | 0 | 0.18 | 0.16 | No |
| Chimpanzee | chr11_hap1_hsa9 | SF2 | Pri | 0.61 | 0.55 | 0.17* | 0.45* | 0.12* | 0.09* | 0.16* | 0.06* | Yes |
| Chimpanzee | chr13_hap1_hsa2b | SF2 | Pri | 1.52* | 0.04* | 0.00* | 0.46* | 0.07* | 0.00* | 0.00* | 0.00* | Yes |
| Bonobo | chr11_mat_hsa9 | SF2 | Pri | 0.34* | 0.26* | 0.08* | 0.19 | 0.45 | 0.08 | 0.1 | 0.21 | Yes |
| Human | chr2 | SF2 | Pri | 0.11* | 0.00* | 0.01* | 0.07* | 0.03* | 0.00* | 0.01* | 0.00* | Yes |

| | | | | | | | | | | | | |
|---|---|---|---|---|---|---|---|---|---|---|---|---|
| Human | chr4 | SF2 | Pri | 0.23* | 0.02* | 0.00* | 0.72 | 0.02* | 0.00* | 0.00* | 0.00* | Yes |
| Human | chr8 | SF2 | Pri | 0.01* | 0.00* | 0.00* | 0.25* | 0.01* | 0.00* | 0.00* | 0.00* | Yes |
| Human | chr9 | SF2 | Pri | 0.06* | 0.02* | 0.00* | 0.28* | 0.05 | 0.00* | 0 | 0.00* | Yes |
| Human | chr13 | SF2 | Pri | 2.91* | 0.00* | 0.00* | 1.04 | 0.81 | 0.00* | 0.00* | 0 | Yes |
| Human | chr14 | SF2 | Pri | 0.01* | 0.00* | 0.00* | 0.29* | 0.11* | 0.00* | 0.00* | 0.01* | Yes |
| Human | chr15 | SF2 | Pri | 1.88 | 0.02* | 0.00* | 1.02 | 0.02 | 0 | 0 | 0 | Yes |
| Human | chr18 | SF2 | Pri | 1.04* | 0.00* | 0.00* | 0.21* | 0.02* | 0.00* | 0.00* | 0.00* | Yes |
| Human | chr20 | SF2 | Pri | 0.01* | 0.00* | 0.00* | 0.20* | 0.9 | 0.00* | 0.00* | 0.00* | Yes |
| Human | chr21 | SF2 | Pri | 1.75 | 0.00* | 0.01* | 1.15 | 0.24* | 0.00* | 0.00* | 0.00* | Yes |
| Human | chr22 | SF2 | Pri | 0.04* | 0.00* | 0.00* | 0.31* | 0.04* | 0.00* | 0.00* | 0.06* | Yes |
| Gorilla | chr12_pat_hsa2a | SF2 | Pri | 0.2 | 0.02* | 0.00* | 0.15* | 0.11 | 0.01 | 0 | 0 | Yes |
| Gorilla | chr13_pat_hsa9 | SF2 | Pri | 0.41 | 0.02* | 0.01* | 0.26 | 0.13 | 0.02 | 0.05* | 0.02 | Yes |
| Gorilla | chr14_pat_hsa13 | SF2 | Pri | 0.43 | 0.04* | 0.03* | 0.43* | 0.15 | 0.09 | 0.1 | 0.08 | Yes |
| Gorilla | chr15_pat_hsa14 | SF2 | Pri | 1.05 | 0.02* | 0.00* | 0.18 | 0.16 | 0.01 | 0 | 0.01 | Yes |
| Gorilla | chr16_pat_hsa15 | SF2 | Pri | 0.12 | 0.00* | 0.00* | 0.22* | 0.08 | 0.01 | 0 | 0 | Yes |
| Gorilla | chr17_mat_hsa18 | SF2 | Pri | 0.64 | 0.16* | 0.07* | 0.68 | 0.16 | 0.1 | 0.19* | 0.08 | Yes |
| Gorilla | chr19_pat_hsa5x17 | SF2 | Pri | 0.13 | 0.33 | 0.00* | 0.35 | 0.12* | 0.00* | 0.01* | 0.00* | Yes |
| Gorilla | chr22_mat_hsa21 | SF2 | Pri | 1.88* | 0.01* | 0.00* | 0.26 | 0.21* | 0.00* | 0.00* | 0.00* | Yes |
| Gorilla | chr23_mat_hsa22 | SF2 | Pri | 3.08* | 0.01* | 0.00* | 0.31 | 0.08* | 0.01* | 0.00* | 0.00* | Yes |
| Chimpanzee | chr11_hap2_hsa9 | SF2 | Alt | 0.31 | 0.27* | 0.10* | 0.30* | 0.20* | 0.05* | 0.09* | 0.04* | Yes |
| Chimpanzee | chr13_hap2_hsa2b | SF2 | Alt | 3.12* | 0.00* | 0.00* | 0.72 | 0.06* | 0.00* | 0.00* | 0.00* | Yes |
| Bonobo | chr11_pat_hsa9 | SF2 | Alt | 0.20* | 0.01* | 0.00* | 0.12 | 0.26 | 0.00* | 0.00* | 0.03* | Yes |
| Gorilla | chr12_mat_hsa2a | SF2 | Alt | 0.24 | 0.05* | 0.00* | 0.12* | 0.05 | 0.02 | 0.01* | 0 | Yes |
| Gorilla | chr13_mat_hsa9 | SF2 | Alt | 0.3 | 0.01* | 0.01* | 0.22 | 0.1 | 0.02 | 0.04* | 0.02 | Yes |
| Gorilla | chr14_mat_hsa13 | SF2 | Alt | 0.42 | 0.04* | 0.02* | 0.4 | 0.14 | 0.09 | 0.09 | 0.06 | Yes |
| Gorilla | chr15_mat_hsa14 | SF2 | Alt | 0.97 | 0.03* | 0.00* | 0.16 | 0.19 | 0.01 | 0 | 0.01 | Yes |
| Gorilla | chr16_mat_hsa15 | SF2 | Alt | 0.56 | 0.17 | 0.06* | 0.36* | 0.16 | 0.14 | 0.2 | 0.1 | Yes |
| Gorilla | chr17_pat_hsa18 | SF2 | Alt | 0.55 | 0.13* | 0.04* | 0.72 | 0.26 | 0.07 | 0.13 | 0.06 | Yes |
| Gorilla | chr19_mat_hsa5x17 | SF2 | Alt | 0.47 | 0.28 | 0.00* | 0.28 | 0.21* | 0.00* | 0.04 | 0.00* | Yes |
| Gorilla | chr22_pat_hsa21 | SF2 | Alt | 2.05* | 0.07* | 0.00* | 0.22 | 0.17* | 0.00* | 0.00* | 0.00* | Yes |
| Gorilla | chr23_pat_hsa22 | SF2 | Alt | 2.55 | 0.00* | 0.00* | 0.27 | 0.31 | 0.01* | 0.00* | 0.00* | Yes |
| Bonobo | chr13_mat_hsa2b | SF2,SF02 | Pri | 1.31 | 0.38 | 0.01 | 0.79 | 0.61 | 0 | 0 | 0 | Yes |
| Bonobo | chr13_pat_hsa2b | SF2,SF02 | Alt | 1.41 | 0.02* | 0.02 | 0.82 | 1.06 | 0 | 0 | 0 | Yes |
| Chimpanzee | chr1_hap1_hsa1 | SF3 | Pri | 0.14* | 0.04* | 0.00* | 0.89 | 0.11* | 0.00* | 0.01* | 0.00* | Yes |
| Chimpanzee | chr5_hap1_hsa6 | SF3 | Pri | 0.32* | 0.17* | 0.00* | 0.72* | 0.00* | 0.00* | 0.00* | 0.00* | Yes |
| Chimpanzee | chr9_hap1_hsa11 | SF3 | Pri | 0.26* | 0.02* | 0.01* | 1.20* | 0.45* | 0.00* | 0.02* | 0.02* | Yes |
| Chimpanzee | chr12_hap1_hsa2a | SF3 | Pri | 0.01* | 0.51 | 0.00* | 0.53 | 0.08* | 0.00* | 0.00* | 0.00* | Yes |
| Chimpanzee | chr19_hap1_hsa17 | SF3 | Pri | 0.07* | 0.17* | 0.00* | 0.99 | 0.49 | 0.00* | 0.00* | 0.00* | Yes |
| Chimpanzee | chrX_hap1_hsaX | SF3 | Pri | 0.04* | 0.00* | 0.01* | 1.57* | 0.00* | 0.00* | 0.01* | 0.01* | Yes |
| Bonobo | chr5_pat_hsa6 | SF3 | Pri | 0.38 | 1.21 | 0 | 0.73 | 0.00* | 0.00* | 0 | 0 | Yes |
| Bonobo | chr9_pat_hsa11 | SF3 | Pri | 0.23* | 0.00* | 0.01* | 0.8 | 1.19* | 0.01* | 0.01* | 0.00* | Yes |
| Bonobo | chr12_mat_hsa2a | SF3 | Pri | 0.41 | 0.27 | 0.00* | 0.73 | 0.16* | 0 | 0 | 0 | Yes |
| Bonobo | chr19_pat_hsa17 | SF3 | Pri | 1.5 | 0.01* | 0.01* | 1.01* | 0.05* | 0.00* | 0.03* | 0.00* | Yes |
| Bonobo | chrX_mat_hsaX | SF3 | Pri | 0.02* | 0.00* | 0.00* | 1.17* | 0.08* | 0.01* | 0.00* | 0.01* | Yes |
| Human | chr11 | SF3 | Pri | 4.98* | 0.00* | 0.00* | 3.46* | 0.00* | 0.00* | 0.00* | 0.00* | Yes |
| Human | chr17 | SF3 | Pri | 0.01* | 0.01* | 0.00* | 1.26* | 0.02* | 0.00* | 0.00* | 0.00* | Yes |
| Human | chrX | SF3 | Pri | 0.15* | 0.01* | 0.00* | 3.11* | 0.05* | 0.00* | 0.00* | 0.00* | Yes |
| Chimpanzee | chr1_hap2_hsa1 | SF3 | Alt | 0.36* | 0.07* | 0.00* | 0.93 | 0.07* | 0.00* | 0.00* | 0.00* | Yes |
| Chimpanzee | chr5_hap2_hsa6 | SF3 | Alt | 0.52* | 0.04* | 0.00* | 0.70* | 0.00* | 0.00* | 0.00* | 0.00* | Yes |
| Chimpanzee | chr9_hap2_hsa11 | SF3 | Alt | 0.38 | 0.01* | 0.01* | 1.06 | 0.52* | 0.00* | 0.02* | 0.02* | Yes |
| Chimpanzee | chr12_hap2_hsa2a | SF3 | Alt | 0.02* | 0.16* | 0.00* | 0.52* | 0.04* | 0.00* | 0.00* | 0.00* | Yes |

| | | | | | | | | | | | |
|---|---|---|---|---|---|---|---|---|---|---|---|
| Chimpanzee | chr19_hap2_hsa17 | SF3 | Alt | 0.07* | 0.07* | 0.00* | 0.96 | 0.44 | 0.00* | 0.00* | 0.00* | Yes |
| Bonobo | chr5_mat_hsa6 | SF3 | Alt | 0.52 | 0.41 | 0.02 | 0.96 | 0.11* | 0 | 0 | 0.09 | Yes |
| Bonobo | chr9_mat_hsa11 | SF3 | Alt | 0.12* | 0.00* | 0.00* | 0.83 | 0.50* | 0.00* | 0.00* | 0.00* | Yes |
| Bonobo | chr12_pat_hsa2a | SF3 | Alt | 0.32 | 0.00* | 0.00* | 0.73 | 0.05* | 0.00* | 0 | 0 | Yes |
| Bonobo | chr19_mat_hsa17 | SF3 | Alt | 0.36 | 0.03* | 0.00* | 0.93 | 0.04* | 0.00* | 0.02* | 0.00* | Yes |
| Chimpanzee | chrY_hap2_hsaY | SF4 | Pri | 0.7 | 0.01* | 0.15* | 0.25 | 1.23 | 0 | 0 | 1.14 | No |
| Bonobo | chr14_pat_hsa13 | SF4 | Pri | 1.62* | 0.33 | 0.15 | 0.48* | 0.71 | 0.94 | 0.55 | 0.04 | No |
| Bonobo | chr17_mat_hsa18 | SF4 | Pri | 1.46 | 0.11 | 0.11 | 0.43 | 0.48 | 0.77 | 0.4 | 0 | No |
| Bonobo | chr22_pat_hsa21 | SF4 | Pri | 0.96 | 0.23 | 0.04 | 0.91 | 0.51 | 0.19 | 0.08 | 0.32 | Yes |
| Bonobo | chrY_pat_hsaY | SF4 | Pri | 2.02 | 0.12* | 0.12* | 0.41 | 1.99 | 0.07 | 0.08 | 0.26 | No |
| Human | chrY | SF4 | Pri | 4.35 | 0.00* | 0.00* | 0.49 | 0.15 | 1.22 | 0 | 0 | No |
| S. orangutan | chr10_hap1_hsa12 | SF4 | Pri | 2.56* | 0.14* | 0.13* | 0.97 | 0.36* | 0.08* | 0.39 | 0.52 | Yes |
| Bonobo | chr22_mat_hsa21 | SF4 | Alt | 1.1 | 0.05 | 0.06 | 0.73 | 0.44* | 0.29 | 0.1 | 0.33 | Yes |
| S. orangutan | chr10_hap2_hsa12 | SF4 | Alt | 3.32* | 0.11* | 0.14* | 0.94 | 0.32* | 0.05* | 0.31 | 0.4 | Yes |
| Chimpanzee | chr4_hap1_hsa5 | SF5 | Pri | 0.21* | 0.48 | 0.02* | 0.36* | 0.08* | 0.01* | 0.04* | 0.05* | Yes |
| Bonobo | chr1_pat_hsa1 | SF5 | Pri | 1.65* | 0.01* | 0.06* | 1.22* | 0.58 | 0.00* | 0.17 | 0.14* | Yes |
| Bonobo | chr3_pat_hsa4 | SF5 | Pri | 0.13* | 0.03* | 0.19* | 0.67* | 0.08* | 0.08* | 0.34 | 0.19* | No |
| Bonobo | chr4_mat_hsa5 | SF5 | Pri | 1.11 | 0.06* | 0.07* | 0.92 | 0.46* | 0.10* | 0.16* | 0.10* | Yes |
| Bonobo | chr7_pat_hsa8 | SF5 | Pri | 1.4 | 0.07* | 0.03 | 1.11 | 0.10* | 0 | 0 | 0.12 | Yes |
| B. orangutan | chr1_hap1_hsa1 | SF5 | Pri | 0.66 | 0.03* | 0.02* | 1.07 | 1.34* | 0.04* | 0.02* | 0.12* | Yes |
| B. orangutan | chr2_hap1_hsa3 | SF5 | Pri | 0.10* | 0.02* | 0.05* | 1.88* | 0.39 | 0.05* | 0.02* | 0.01* | Yes |
| B. orangutan | chr3_hap1_hsa4 | SF5 | Pri | 0.12* | 0.00* | 0.02* | 2.07* | 2.30* | 0.00* | 0.01* | 0.21* | Yes |
| B. orangutan | chr4_hap1_hsa5 | SF5 | Pri | 0.17* | 0.02* | 0.10* | 0.91 | 0.16* | 0.14* | 0.08* | 0.04* | Yes |
| B. orangutan | chr5_hap1_hsa6 | SF5 | Pri | 0.33* | 0.02* | 0.01* | 0.77* | 0.39* | 0.05* | 0.02* | 0.06* | Yes |
| B. orangutan | chr6_hap1_hsa7 | SF5 | Pri | 0.64 | 0.57 | 0.76 | 1.22* | 0.69 | 0.82 | 0.59 | 0.47* | Yes |
| B. orangutan | chr7_hap1_hsa8 | SF5 | Pri | 1.71* | 0.09* | 0.01* | 1.30* | 1.50* | 0.02* | 0.02* | 0.14* | Yes |
| B. orangutan | chr8_hap1_hsa10 | SF5 | Pri | 0.53 | 0.67 | 0.41 | 0.49 | 4.20* | 0.05* | 0.03* | 0.04* | Yes |
| B. orangutan | chr9_hap1_hsa11 | SF5 | Pri | 0.96 | 2.93 | 1.58 | 0.89 | 0.77 | 0.12* | 0.09* | 0.05* | Yes |
| B. orangutan | chr11_hap1_hsa2b | SF5 | Pri | 0.17* | 0.02* | 0.03* | 1.15 | 0.14* | 0.04* | 0.07* | 0.04* | Yes |
| B. orangutan | chr12_hap1_hsa2a | SF5 | Pri | 1.11 | 0.57 | 0.47* | 0.94 | 0.89 | 0.11* | 0.23 | 0.22 | Yes |
| B. orangutan | chr13_hap1_hsa9 | SF5 | Pri | 0.74 | 0.59 | 0.44* | 0.88 | 0.24* | 0.04* | 0.14* | 0.1 | Yes |
| B. orangutan | chr14_hap1_hsa13 | SF5 | Pri | 0.25* | 0.02* | 0.02* | 1.07 | 0.23* | 0.03* | 0.06* | 0.07 | Yes |
| B. orangutan | chr15_hap1_hsa14 | SF5 | Pri | 0.30* | 0.03* | 0.05* | 0.74 | 0.37 | 0.04* | 0.15* | 0.14 | Yes |
| B. orangutan | chr16_hap1_hsa15 | SF5 | Pri | 0.23* | 0.05* | 0.02* | 0.99 | 0.56* | 0.04* | 0.07* | 0.11 | Yes |
| B. orangutan | chr17_hap1_hsa18 | SF5 | Pri | 1.19 | 0.43 | 0.33* | 0.91 | 0.79 | 0.08* | 0.15* | 0.14 | Yes |
| B. orangutan | chr18_hap1_hsa16 | SF5 | Pri | 0.06* | 0.00* | 0.07* | 1.22 | 1.50* | 0.00* | 0.00* | 1.09 | No |
| B. orangutan | chr19_hap1_hsa17 | SF5 | Pri | 0.3 | 0.04* | 0.04* | 0.58 | 1.73* | 0.02* | 0.01* | 0.02* | Yes |
| B. orangutan | chr20_hap1_hsa19 | SF5 | Pri | 0.16* | 0.01* | 0.01* | 2.37* | 1.56 | 0.02* | 0.01* | 0.02* | Yes |
| B. orangutan | chr21_hap1_hsa20 | SF5 | Pri | 0.03* | 0.05* | 0.04* | 1.08* | 0.12* | 0.03* | 0.06* | 0.00* | No |
| B. orangutan | chr22_hap1_hsa21 | SF5 | Pri | 0.62 | 0.06* | 0.10* | 1.12 | 0.63 | 0.08* | 0.24 | 0.29 | Yes |
| B. orangutan | chr23_hap1_hsa22 | SF5 | Pri | 0.33* | 0.02* | 0.04* | 1.19* | 0.34* | 0.06* | 0.07* | 0.07 | Yes |
| B. orangutan | chrX_hap1_hsaX | SF5 | Pri | 0.26* | 0.02* | 0.07* | 1.92* | 2.57* | 0.02* | 0.03* | 0.05* | Yes |
| B. orangutan | chrY_hap2_hsaY | SF5 | Pri | 0.59* | 0.02* | 0.05* | 1.50* | 1.2 | 0 | 0.00* | 0 | No |
| S. orangutan | chr1_hap1_hsa1 | SF5 | Pri | 0.7 | 0.03* | 0.02* | 0.92 | 2.38* | 0.04* | 0.02* | 0.07* | Yes |
| S. orangutan | chr2_hap1_hsa3 | SF5 | Pri | 0.61 | 0.14 | 0.06* | 2.03* | 1.24* | 0.03* | 0.01* | 0.01* | Yes |
| S. orangutan | chr3_hap1_hsa4 | SF5 | Pri | 0.2 | 0.00* | 0.03* | 2.63* | 2.81* | 0.00* | 0.00* | 0.5 | Yes |
| S. orangutan | chr4_hap1_hsa5 | SF5 | Pri | 0.15* | 0.01* | 0.11* | 0.89 | 0.15* | 0.05* | 0.03* | 0.04* | Yes |
| S. orangutan | chr5_hap1_hsa6 | SF5 | Pri | 0.33* | 0.01* | 0.00* | 0.95 | 0.72* | 0.01* | 0.01* | 0.11* | Yes |
| S. orangutan | chr6_hap1_hsa7 | SF5 | Pri | 0.10* | 0.02* | 0.03* | 1.36* | 0.11* | 0.04* | 0.04* | 0.01* | Yes |
| S. orangutan | chr7_hap1_hsa8 | SF5 | Pri | 1.40* | 0.12* | 0.01* | 0.93 | 0.9 | 0.02* | 0.01* | 0.09* | Yes |
| S. orangutan | chr8_hap1_hsa10 | SF5 | Pri | 0.33 | 0.71 | 0.41 | 0.59 | 3.26* | 0.21* | 0.05* | 0.15* | Yes |

| | | | | | | | | | | | | |
|---|---|---|---|---|---|---|---|---|---|---|---|---|
| S. orangutan | chr9_hap1_hsa11 | SF5 | Pri | 0.81 | 2.49 | 1.29 | 0.9 | 0.94 | 0.09* | 0.07* | 0.04* | Yes |
| S. orangutan | chr11_hap1_hsa2b | SF5 | Pri | 0.24* | 0.03* | 0.02* | 1.19 | 0.24* | 0.00* | 0.05* | 0.07* | Yes |
| S. orangutan | chr12_hap1_hsa2a | SF5 | Pri | 1.04 | 0.46 | 0.37* | 0.74 | 0.44* | 0.11* | 0.22 | 0.19 | Yes |
| S. orangutan | chr13_hap1_hsa9 | SF5 | Pri | 0.91 | 0.76 | 0.55* | 0.92 | 0.32* | 0.07* | 0.21* | 0.15 | Yes |
| S. orangutan | chr14_hap1_hsa13 | SF5 | Pri | 0.12* | 0.01* | 0.00* | 1.11 | 0.08* | 0.03* | 0.01* | 0.00* | No |
| S. orangutan | chr15_hap1_hsa14 | SF5 | Pri | 0.08* | 0.15* | 0.00* | 0.77 | 0.08* | 0.00* | 0.00* | 0.01* | No |
| S. orangutan | chr16_hap1_hsa15 | SF5 | Pri | 0.22* | 0.02* | 0.02* | 1.18* | 0.15* | 0.02* | 0.05* | 0.05* | Yes |
| S. orangutan | chr17_hap1_hsa18 | SF5 | Pri | 0.06* | 0.08* | 0.03* | 0.61 | 0.14* | 0.03* | 0.04* | 0.04* | No |
| S. orangutan | chr18_hap1_hsa16 | SF5 | Pri | 0.03* | 0.04* | 0.04* | 1.17* | 1.50* | 0.01* | 0.01* | 0.9 | No |
| S. orangutan | chr19_hap1_hsa17 | SF5 | Pri | 0.12* | 0.05* | 0.05* | 0.79 | 2.28* | 0.00* | 0.00* | 0.02* | Yes |
| S. orangutan | chr20_hap1_hsa19 | SF5 | Pri | 2.31* | 0.02* | 0.01* | 2.52* | 1.38 | 0.02* | 0.02* | 0.05* | Yes |
| S. orangutan | chr21_hap1_hsa20 | SF5 | Pri | 0.11 | 0.02* | 0.01* | 1.16* | 0.16* | 0.01* | 0.02* | 0.02* | Yes |
| S. orangutan | chr22_hap1_hsa21 | SF5 | Pri | 0.14* | 0.03* | 0.03* | 1.1 | 0.14* | 0.08* | 0.05* | 0.05* | Yes |
| S. orangutan | chr23_hap1_hsa22 | SF5 | Pri | 0.21* | 0.03* | 0.01* | 1.23* | 0.38* | 0.02* | 0.03* | 0.04* | Yes |
| S. orangutan | chrX_hap1_hsaX | SF5 | Pri | 0.24* | 0.05* | 0.18* | 1.81* | 2.26* | 0.04* | 0.04* | 1.15 | Yes |
| S. orangutan | chrY_hap2_hsaY | SF5 | Pri | 0.55* | 0.00* | 0.05* | 1.55* | 1.24 | 0 | 0 | 0 | No |
| Chimpanzee | chr4_hap2_hsa5 | SF5 | Alt | 0.19* | 0.06* | 0.02* | 0.43* | 0.12* | 0.02* | 0.05* | 0.05* | Yes |
| Bonobo | chr1_mat_hsa1 | SF5 | Alt | 2.11* | 0.12* | 0.05* | 1.24* | 0.52 | 0.00* | 0.14 | 0.11* | Yes |
| Bonobo | chr3_mat_hsa4 | SF5 | Alt | 0.25* | 0.06* | 0.16* | 0.66* | 0.15* | 0.19* | 0.10* | 0.18* | No |
| Bonobo | chr4_pat_hsa5 | SF5 | Alt | 0.94 | 0.16* | 0.11* | 0.99 | 0.54* | 0.11* | 0.35 | 0.33* | Yes |
| Bonobo | chr7_mat_hsa8 | SF5 | Alt | 1.57 | 0.10* | 0 | 1.28 | 0.06* | 0 | 0 | 0 | Yes |
| B. orangutan | chr1_hap2_hsa1 | SF5 | Alt | 0.71 | 0.02* | 0.02* | 1.06 | 1.39* | 0.04* | 0.02* | 0.11* | Yes |
| B. orangutan | chr2_hap2_hsa3 | SF5 | Alt | 0.10* | 0.13* | 0.12* | 1.87* | 0.46 | 0.08* | 0.02* | 0.01* | Yes |
| B. orangutan | chr3_hap2_hsa4 | SF5 | Alt | 0.54 | 0.01* | 0.05* | 1.66* | 1.50* | 0.00* | 0.01* | 0.7 | Yes |
| B. orangutan | chr4_hap2_hsa5 | SF5 | Alt | 0.21* | 0.01* | 0.02* | 1.11 | 0.15* | 0.05* | 0.02* | 0.09* | Yes |
| B. orangutan | chr5_hap2_hsa6 | SF5 | Alt | 0.26* | 0.01* | 0.01* | 0.9 | 0.51* | 0.06* | 0.01* | 0.07* | Yes |
| B. orangutan | chr6_hap2_hsa7 | SF5 | Alt | 0.99 | 0.83 | 0.96 | 1.22* | 0.75 | 0.9 | 0.64 | 0.52 | Yes |
| B. orangutan | chr7_hap2_hsa8 | SF5 | Alt | 1.63* | 0.06* | 0.01* | 1.32* | 1.56* | 0.01* | 0.01* | 0.45* | Yes |
| B. orangutan | chr8_hap2_hsa10 | SF5 | Alt | 0.32 | 0.66 | 0.41 | 0.44 | 4.61* | 0.10* | 0.03* | 0.03* | Yes |
| B. orangutan | chr9_hap2_hsa11 | SF5 | Alt | 0.13* | 0.01* | 0.02* | 1.05 | 0.11* | 0.01* | 0.03* | 0.03* | Yes |
| B. orangutan | chr10_hap2_hsa12 | SF5 | Alt | 0.17* | 0.08* | 0.05* | 0.87* | 0.20* | 0.13* | 0.05* | 0.00* | No |
| B. orangutan | chr11_hap2_hsa2b | SF5 | Alt | 0.17* | 0.01* | 0.02* | 1.21* | 0.24* | 0.02* | 0.04* | 0.06* | Yes |
| B. orangutan | chr12_hap2_hsa2a | SF5 | Alt | 1.33 | 0.5 | 0.43* | 0.98 | 1.15 | 0.14* | 0.25 | 0.23 | Yes |
| B. orangutan | chr13_hap2_hsa9 | SF5 | Alt | 0.75 | 0.58 | 0.43* | 0.89 | 0.24* | 0.04* | 0.14* | 0.1 | Yes |
| B. orangutan | chr14_hap2_hsa13 | SF5 | Alt | 0.30* | 0.02* | 0.03* | 1.13 | 0.36* | 0.02* | 0.08* | 0.11 | Yes |
| B. orangutan | chr15_hap2_hsa14 | SF5 | Alt | 0.30* | 0.02* | 0.03* | 1 | 0.31* | 0.02* | 0.06* | 0.08 | Yes |
| B. orangutan | chr16_hap2_hsa15 | SF5 | Alt | 0.31* | 0.05* | 0.03* | 0.95 | 0.50* | 0.05* | 0.09* | 0.14 | Yes |
| B. orangutan | chr17_hap2_hsa18 | SF5 | Alt | 1.28 | 0.5 | 0.39* | 1 | 1.25 | 0.07* | 0.19 | 0.24 | Yes |
| B. orangutan | chr18_hap2_hsa16 | SF5 | Alt | 0.3 | 0.01* | 0.03* | 1.04 | 1.69* | 0.01* | 0.00* | 0.66 | Yes |
| B. orangutan | chr19_hap2_hsa17 | SF5 | Alt | 0.25* | 0.04* | 0.06* | 0.65* | 1.66* | 0.04* | 0.02* | 0.03* | Yes |
| B. orangutan | chr20_hap2_hsa19 | SF5 | Alt | 0.3 | 0.01* | 0.01* | 2.26* | 1.65 | 0.02* | 0.02* | 0.06* | Yes |
| B. orangutan | chr21_hap2_hsa20 | SF5 | Alt | 0.04* | 0.05* | 0.01* | 1.10* | 0.10* | 0.02* | 0.02* | 0.00* | No |
| B. orangutan | chr22_hap2_hsa21 | SF5 | Alt | 0.62 | 0.06* | 0.10* | 1.14 | 0.64 | 0.08* | 0.23 | 0.29 | Yes |
| B. orangutan | chr23_hap2_hsa22 | SF5 | Alt | 0.28* | 0.03* | 0.04* | 1.22* | 0.29 | 0.07 | 0.07 | 0.07 | Yes |
| S. orangutan | chr1_hap2_hsa1 | SF5 | Alt | 0.29* | 0.01* | 0.01* | 1.41* | 0.61 | 0.02* | 0.01* | 0.07* | Yes |
| S. orangutan | chr2_hap2_hsa3 | SF5 | Alt | 0.66 | 0.04* | 0.01* | 1.95* | 1.20* | 0.04* | 0.01* | 0.01* | Yes |
| S. orangutan | chr3_hap2_hsa4 | SF5 | Alt | 0.37 | 0.02* | 0.03* | 1.78* | 1.61* | 0.02* | 0.01* | 0.32 | Yes |
| S. orangutan | chr4_hap2_hsa5 | SF5 | Alt | 0.20* | 0.01* | 0.13* | 0.87 | 0.10* | 0.02* | 0.01* | 0.04* | Yes |
| S. orangutan | chr5_hap2_hsa6 | SF5 | Alt | 0.15* | 0.08* | 0.00* | 0.62* | 0.11* | 0.00* | 0.00* | 0.01* | Yes |
| S. orangutan | chr6_hap2_hsa7 | SF5 | Alt | 0.05* | 0.57 | 0.02* | 1.49* | 0.09* | 0.00* | 0.00* | 0.00* | Yes |
| S. orangutan | chr7_hap2_hsa8 | SF5 | Alt | 1.62* | 0.16* | 0.01* | 0.93 | 1.57* | 0.01* | 0.01* | 0.02* | Yes |

| | | | | | | | | | | | | |
|---|---|---|---|---|---|---|---|---|---|---|---|---|
| S. orangutan | chr8_hap2_hsa10 | SF5 | Alt | 0.35 | 0.79 | 0.41 | 0.46 | 3.05* | 0.41 | 0.03* | 0.06* | Yes |
| S. orangutan | chr9_hap2_hsa11 | SF5 | Alt | 0.88 | 3.17 | 1.51 | 0.82 | 0.85 | 0.06* | 0.08* | 0.04* | Yes |
| S. orangutan | chr11_hap2_hsa2b | SF5 | Alt | 0.18* | 0.04* | 0.02* | 1.28* | 0.17* | 0.01* | 0.04* | 0.05* | Yes |
| S. orangutan | chr12_hap2_hsa2a | SF5 | Alt | 1.04 | 0.66 | 0.47 | 0.7 | 0.50* | 0.08* | 0.16* | 0.14 | Yes |
| S. orangutan | chr13_hap2_hsa9 | SF5 | Alt | 1 | 0.93 | 0.61* | 0.91 | 0.32* | 0.07* | 0.20* | 0.15* | Yes |
| S. orangutan | chr14_hap2_hsa13 | SF5 | Alt | 0.07* | 0.01* | 0.01* | 1.24 | 0.13* | 0.02* | 0.00* | 0.05* | No |
| S. orangutan | chr15_hap2_hsa14 | SF5 | Alt | 0.07* | 0.04* | 0.02* | 0.72 | 0.03* | 0.00* | 0.00* | 0.16 | No |
| S. orangutan | chr16_hap2_hsa15 | SF5 | Alt | 0.24* | 0.02* | 0.04* | 1.13* | 0.22* | 0.02* | 0.06* | 0.06* | Yes |
| S. orangutan | chr17_hap2_hsa18 | SF5 | Alt | 0.07* | 0.12* | 0.03* | 0.61 | 0.15* | 0.04* | 0.04* | 0.03* | No |
| S. orangutan | chr18_hap2_hsa16 | SF5 | Alt | 0.23* | 0.05* | 0.06* | 1.16 | 0.91 | 0.00* | 0.01* | 0.79 | No |
| S. orangutan | chr19_hap2_hsa17 | SF5 | Alt | 0.08* | 0.03* | 0.03* | 0.85 | 3.10* | 0.01* | 0.01* | 0.03* | Yes |
| S. orangutan | chr20_hap2_hsa19 | SF5 | Alt | 0.6 | 0.01* | 0.00* | 2.21* | 1.44 | 0.00* | 0.00* | 0.08 | Yes |
| S. orangutan | chr21_hap2_hsa20 | SF5 | Alt | 0.12 | 0.03* | 0.01* | 1.15* | 0.16* | 0.01* | 0.02* | 0.02* | Yes |
| S. orangutan | chr22_hap2_hsa21 | SF5 | Alt | 0.12* | 0.07* | 0.03* | 1.43* | 0.18* | 0.05* | 0.03* | 0.05* | Yes |
| S. orangutan | chr23_hap2_hsa22 | SF5 | Alt | 0.18* | 0.03* | 0.01* | 1.33* | 0.31* | 0.02* | 0.03* | 0.04* | Yes |
